# Biophysical modeling reveals how recording geometry shapes representations of cortical activity

**DOI:** 10.64898/2026.09.17.752414

**Authors:** A. A. Díaz-Montiel, J. A. Willis, G. Unal, A.M. Abrego, Y. Kajikawa, J. C. Mosher, C. E. Schroeder, S. Dura-Bernal, J. P. Seymour, S. A. Neymotin

## Abstract

**Objective:** Determine how recording geometry shapes the spatial organization of cortical activity represented in extracellular signals.

**Approach:** We combined a biophysical macaque auditory thalamocortical model with forward models of linear laminar (UPROBE) and Directional and Scalable (DiSc) electrodes. Four conditions combined graded thalamic drive (Peak, Mid, Low, and Off-best, in descending order) with distinct fixed connectivity realizations; each was evaluated across 25 background-input realizations. Population-specific extracellular signals were calculated from compartmental transmembrane currents, and experimental tone and broadband-noise (BBN) responses were analyzed for both probe geometries.

**Main results:** Extracellular response patterns and relative contributions of neuronal populations differed across model conditions and signal measures. Layer 2/3 intratelencephalic neurons comprised 39.38% of cortical cells but contributed 54.86% of the normalized multiunit activity (MUA)-like envelope magnitude in the Peak condition; their contributions ranged from 9.76% to 54.86% across the four conditions. Early MUA-like envelope amplitude was 0.00676 a.u. (arbitrary units) higher in Peak than in Mid (95% bootstrap confidence interval 0.00061-0.01272), although these conditions differed in both drive and connectivity. In experimental nonhuman primate (NHP) recordings, Pearson correlations between BBN and the mean tone-evoked spatial response profile averaged 0.802 across six DiSc and 0.875 across eight UPROBE penetrations.

**Significance:** Biophysical source and electrode models help distinguish neural source organization from the spatial information retained by recording geometry. Experimental correlations support shared spatial response structure across tones and noise, subject to penetration dependence and fixed stimulus-order effects.

## Introduction

Understanding how neural circuit activity gives rise to extracellular electrophysiological signals remains a central challenge in systems neuroscience. Recordings of local field potential (LFP) and multiunit activity (MUA) across cortical layers, together with estimates of current source density (CSD), provide related views of population-level neural activity. MUA predominantly reflects local suprathreshold spiking, whereas extracellular field potentials arise from the superposition of transmembrane currents generated by synaptic activity, intrinsic membrane conductances, and action potentials; CSD analysis estimates the spatial distribution of the underlying current sinks and sources^1–4^. The magnitude and spatial structure of these extracellular signals depend not only on neuronal activity itself, but also on cellular morphology, cytoarchitectural organization, synchrony, volume conduction, and the spatial relationship between neuronal current sources and the recording sites^1,2,5,6^. Consequently, LFP, CSD and MUA measures provide related but non-equivalent representations of cortical activity.

The relationship between neural activity and the measured extracellular signal is further shaped by the recording apparatus. Computational studies of silicon microelectrodes have shown that neuronal position, contact dimensions, and electrode configuration influence recorded amplitudes and sensitivity^7^, while physical models demonstrate that recording-site dimensions affect the measured extracellular potential^8^. High-density and spatially oversampled electrode arrays provide increasingly fine spatial sampling of extracellular activity^9,10^, although different sampling considerations may apply when the objective is to maximize single-unit yield^11^. Forward models incorporating electrode sensitivity have further demonstrated that device placement and configuration influence the information recoverable from underlying cortical LFP sources^12^. These considerations are increasingly relevant as neural engineering develops interfaces with greater spatial density and more diverse geometries^13^. Recording geometry should therefore be considered part of the measurement transformation rather than merely a passive means of observing a pre-existing extracellular signal.

Biophysically detailed computational models provide a means of linking these experimentally accessible signals to their underlying cellular mechanisms^2,6^. Forward models can compute extracellular potentials from transmembrane currents in multicompartment neurons and neuronal populations, enabling simultaneous prediction of membrane dynamics, spiking, LFP, CSD, and other extracellular observables^2,14–17^. Such approaches have been used both in detailed cortical microcircuits and in hybrid population models to reproduce experimentally observed field potentials and investigate the network processes that generate them^14–16^. Recently, we developed a data-driven multiscale model of macaque primary auditory cortex (A1), medial geniculate body, and thalamic reticular nucleus that reproduces physiological firing rates, laminar LFP/CSD patterns, and oscillatory activity observed in vivo^18^. Because the same simulated network generates cellular activity and multiple extracellular observables, such models make it possible to manipulate a defined circuit process and follow its consequences through successive stages of signal generation.

This ability is particularly useful for distinguishing robust organization in the underlying neural activity from its expression in individual extracellular signals. Biological neurons and circuits exhibit substantial variability in cellular and synaptic parameters, and similar network-level outputs can arise from different underlying parameter combinations^19,20^. Accordingly, validation need not require point-by-point reproduction of every waveform from an individual preparation; models can instead be evaluated against experimentally constrained observables and robust relationships in population activity^18–21^. Controlled manipulation of a circuit variable provides one way to establish such relationships. In the auditory thalamocortical system, constructing cortical-column conditions spanning graded thalamic recruitment therefore provides a way to order afferent drive and follow its expression in cortical population activity and the resulting LFP, CSD, and MUA representations.

Biophysical circuit models have extensively characterized how cellular and network activity gives rise to extracellular signals within specified virtual or experimental recording arrangements^14–18,22^. Less is known about how organization predicted at the circuit level is transformed when common underlying cortical sources are sampled through substantially different recording geometries. This distinction is increasingly important as neural interfaces expand from conventional linear laminar probes toward high-density and multidimensional electrode configurations with different spatial sampling properties^9,10,12,13,23–25^. One example is the Directional and Scalable (DiSc) microelectrode array, a depth electrode that uses small contacts distributed around the circumference of an insulating shaft to provide directional sensitivity to surrounding neural sources^12,26–28^. Comparing substantially different geometries under experimentally controlled conditions is difficult because separate devices cannot generally sample precisely the same biological sources. Forward modeling provides a way around this limitation by allowing recording geometry to be varied while holding the underlying source activity constant.

We therefore hypothesized that organizational relationships generated by a biophysically realistic circuit model would be expressed across extracellular signals and transformed in geometry-dependent ways during spatial sampling. We tested this hypothesis in successive stages using the macaque auditory thalamocortical model^18^. First, we evaluated four separately instantiated cortical-column conditions spanning graded thalamic recruitment to determine whether the imposed response levels were associated with corresponding organization across LFP, CSD, and MUA and to identify the neuronal populations contributing to these signals. We then constructed controlled spatial configurations of independently simulated cortical columns and sampled the same underlying sources using virtual linear laminar and DiSc arrays, allowing the effects of recording geometry to be isolated from differences in neuronal dynamics.

Finally, we tested whether the forms of spatial organization predicted for each geometry were supported by experimental macaque auditory-cortex recordings obtained with the corresponding electrode configurations. Rather than requiring point-by-point correspondence between simulated and experimental responses, we evaluated whether the predicted organizational relationships were identifiable across heterogeneous penetrations exhibiting variation in response magnitude and frequency preference. This framework separates successive contributions of circuit activity, extracellular signal generation, and recording geometry, providing a principled basis for interpreting how neural-interface design determines which dimensions of cortical organization are accessible experimentally.

## Methods

### Biophysical model

Numerical simulations were performed using a version of the macaque auditory thalamocortical model based on our previously published model^18^, implemented in NetPyNE^29^ and NEURON^30^. The model retains the general architecture of the published version but incorporates updated parameter settings. The original publication describes the model architecture and its physiological validation, while the complete model implementation, parameter values, and simulation configurations used in the present study are available in the repositories described below in **Modeling and analysis software**.

The model comprises approximately 12K biophysically detailed neurons distributed across six cortical layers together with thalamic relay populations and over 25 million synapses, incorporating experimentally constrained neuronal morphologies, intrinsic membrane properties, synaptic dynamics, and local and long-range connectivity. The network reproduces a broad range of anatomical and physiological observations from the macaque auditory cortex, including laminar organization, spontaneous activity, sensory-evoked responses, and extracellular electrophysiological signals. Because the present study uses this published model with only minor modifications to its cellular or network architecture, detailed descriptions of the neuronal populations, connectivity rules, parameter optimization, and model validation are not repeated here.

To investigate how graded thalamic recruitment is expressed in extracellular electrophysiological signals, we generated four separately instantiated cortical-column conditions, each combining a fixed network connectivity realization with one level of externally generated excitatory synaptic input to the thalamocortical (TC) and high-threshold thalamocortical (HTC) relay populations. Stimulus events were delivered to somatic AMPA synapses using NEURON NetStim event generators, and input strength was set by the peak AMPA synaptic conductance. The four operational response levels (Peak, Mid, Low, and Off-best) used TC/HTC peak conductances of 20, 16, 8, and 3 µS, respectively; input to the matrix thalamocortical (TCM) relay population was held at 5 µS across conditions. One matched baseline without stimulus-driven thalamic input was included for each connectivity realization.

Each condition represented a separate cortical-column realization generated using a fixed network connectivity random seed (*connseed* in the saved model configurations); the seeds for Peak, Mid, Low, and Off-best were 1,2,3, and 4, respectively. These operational labels describe representative levels within an imposed column-response profile rather than separately simulated acoustic frequencies, empirically fitted tuning curves, or anatomically registered A1 locations. For each fixed column/drive condition, 25 repeated simulations were generated by varying the random seed governing stochastic background/noise inputs (*stimseed* in the saved model configurations), while preserving the column connectivity and deterministic thalamic stimulus timing. One matched baseline was analyzed for each condition using the same connectivity realization but without stimulus-driven thalamic input. Thus, the final analysis comprised 100 stimulated simulations and four matched baseline simulations (104 outputs).

Thalamic stimulation began at 3000 ms, with deterministic synaptic input events delivered at 100-ms intervals without temporal jitter or synaptic delay. Simulations ended at 3600 ms. The resulting network activity and extracellular responses were evaluated using the analysis-specific time windows described below.

Simulations were integrated using a fixed time step of 0.05 ms, with selected state variables recorded every 0.1 ms (10 kHz). Throughout each simulation, membrane voltages, neuronal spiking activity, and extracellular quantities required for subsequent analyses were stored for offline processing.

### Extracellular signal generation

For the graded-recruitment and population-contribution analyses (**Figs. 2-3**), extracellular potentials were computed using the forward-modeling framework implemented in NetPyNE, which uses the line-source approximation to sum contributions from transmembrane currents across all compartments of all modeled neurons^31^. Potentials were computed at virtual recording contacts spaced 100 µm apart along the cortical depth axis, using a homogeneous conductivity of 0.3 S/m.

For the device analyses (**Fig. 4**), cortical compartment-level transmembrane currents from the same biophysical cortical column simulations were spatially arranged and projected through FEM-derived lead fields to compute device-specific extracellular voltages (using conductivity of 0.25 S/m; see **Device simulation & multi-column organizations**). Values obtained from the two forward models are not on a common scale. CSD was subsequently computed from the multichannel LFP recordings along the cortical depth axis (see below **Graded thalamic recruitment analysis**). For the graded-recruitment and population-contribution analyses, MUA was represented by a model-derived high-frequency envelope computed from the broadband simulated extracellular potential using the signal-processing procedure described below. Simulation time-series were aligned to stimulus onset before analysis.

For the device comparison in **Fig. 4**, the same underlying multi-column source configurations were sampled using virtual implementations of the DiSc array and the linear UPROBE. The same signal-processing and analysis pipelines were applied across both recording geometries unless otherwise stated.

### Graded thalamic recruitment analysis

Evoked responses were quantified for the four fixed column/drive conditions described above. SEM was computed across the 25 stochastic background-input realizations within each condition and therefore reflects variability across these realizations, not across network connectivity realizations. Because connectivity realization and thalamic input strength differed together across conditions, between-condition comparisons characterize the four operational column/drive states rather than the independent effect of input strength in a common network. Signals were aligned to thalamic stimulus onset, and LFP, CSD, and MUA responses were baseline corrected by subtracting the temporal mean during the -50 to 0 ms prestimulus interval. The early evoked response was defined as the 0-25 ms post-stimulus interval.

CSD was estimated from the depth-resolved LFP as the negative second spatial derivative along the cortical depth axis. Before CSD estimation, LFP signals were arranged as depth × time, the temporal mean was removed independently from each depth channel, and Gaussian smoothing (σ = 0.5 channels) was applied along the depth dimension. For the equally spaced recording contacts, CSD was calculated using the finite-difference approximation

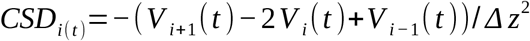

where *V _i_*(*t*) denotes the LFP at depth index i and *Δz* = 100 µm is the inter-electrode spacing. Because the second spatial derivative is defined only for interior recording locations, the **Figure 2** CSD heat map and response metric used all interior recording depths. CSD values were scaled for visualization and are therefore reported in arbitrary units rather than as absolute physical current-density measurements.

The model-derived MUA envelope (hereafter MUA) was computed from the broadband simulated extracellular potential at each recording contact. Each contact’s signal was high-pass filtered above 300 Hz with a fourth-order zero-phase Butterworth filter, full-wave rectified, and smoothed with a Gaussian kernel (*σ* =1 ms)^32^. Each envelope was baseline corrected by subtracting its -50 to 0 ms prestimulus mean: negative values after correction were clipped to zero. The contact envelopes were then averaged to obtain one MUA trace per simulation. No explicit upper-frequency filter cutoff was applied; the 10 kHz signal has a 5 kHz Nyquist limit.

For temporal response analysis, baseline-corrected traces were averaged across the 25 stochastic background-input realizations within each condition. Temporal responses are reported as mean ± SEM across these realizations. Each condition’s matched baseline was processed using the same procedures and is shown as a reference; the matched-baseline trace was not subtracted from the stimulated trace. The four-condition response profiles were summarized descriptively. The separate Peak–Mid comparison described below treated realizations as independent between conditions, because each condition combined a different fixed connectivity realization with a different thalamic input strength.

Early contact-averaged MUA-envelope amplitudes were compared between Peak and Mid using an independent within-condition percentile bootstrap of the mean difference (100,000 resamples), a two-sided Welch test and Hedges g. Each condition included 25 background-input realizations. The conditions had different fixed connectivity realizations as well as different input strengths, so the comparison does not isolate either factor. Realizations were treated as independent between conditions because matching random-seed labels do not imply matching background inputs across different networks. **Fig. 2** response amplitudes were calculated from signals interpolated onto a common time grid, whereas **Fig. 3** population contributions were calculated using the original sampled signal values.

### Population-specific contribution analysis

Population-specific contributions to extracellular signals were quantified in the Peak, Mid, Low and Off-best conditions, with 25 stochastic background-input realizations per condition. Neurons were grouped into ten cortical population classes: L2/3 IT (IT2 and IT3), L4 SS (ITS4), L4 IT (ITP4), L5 IT (IT5A and IT5B), L5 PT (PT5B), mixed deep-layer IT/CT (IT6, CT5A, CT5B, and CT6), PV (PV2, PV3, PV4, PV5A, PV5B, and PV6), SST (SOM2, SOM3, SOM4, SOM5A, SOM5B, and SOM6), VIP (VIP2, VIP3, VIP4, VIP5A, VIP5B, and VIP6), and NGF (NGF1, NGF2, NGF3, NGF4, NGF5A, NGF5B, and NGF6). Population contributions were quantified over the 0-25 ms post-stimulus interval.

These ten classes provide a compact descriptive grouping, not an exhaustive cortical taxonomy. Excitatory classes retain laminar/projection distinctions, whereas inhibitory classes pool layers and therefore hide within-class laminar variation. IT denotes intratelencephalic; SS, spiny stellate; PT, pyramidal tract; CT, corticothalamic; PV, parvalbumin-expressing; SST/SOM, somatostatin-expressing; VIP, vasoactive intestinal peptide-expressing; and NGF, neurogliaform. Native suffixes denote layers. Signed population LFPs are summed within each class before filtering, rectification and envelope extraction; nonlinear MUA-envelope fractions are not an exact additive decomposition of total MUA.

Population-specific LFPs were reconstructed separately for each neuronal population and stochastic input realization from saved compartment transmembrane currents and geometry using the line-source forward calculation described above. The 43 native population voltages were summed with their signs intact, and their total agreed with the saved whole-network LFP. When multiple populations belonged to the same class, their LFPs were summed before response quantification, reflecting the linear additivity of extracellular potentials. Each recording-contact trace was baseline corrected independently by subtracting its mean during the half-open [-50, 0) ms prestimulus interval. Population-specific LFP response magnitude was quantified as the root-mean-square (RMS) amplitude across recording contacts and time samples during the half-open [0, 25) ms post-stimulus interval.

Population-specific CSD was calculated from the corresponding grouped LFP. The 20 recording contacts were ordered by depth at 100-µm spacing over a 1.9-mm span. The negative second spatial derivative was obtained by applying the coordinate-based gradient twice, with second-order boundary derivatives; all 20 contacts were retained. This population-contribution calculation differs from the smoothed interior-contact finite difference used for **Fig. 2**. A constant tissue-conductivity scaling factor was omitted because contributions were normalized within each realization. CSD was baseline corrected independently at each depth over [-50, 0) ms, and response magnitude was the RMS across all depths and samples in [0, 25) ms.

For the population-specific MUA envelope analysis, population-specific LFP contributions belonging to the same cortical population class were first summed to reconstruct a grouped extracellular signal for each class. Summation was performed before envelope extraction because the underlying extracellular potential is linearly additive, whereas rectification and the resulting envelope transformation are nonlinear. Each grouped signal was processed at each contact by fourth-order zero-phase Butterworth high-pass filtering at 300 Hz, absolute-value rectification and Gaussian smoothing with σ = 1 ms, truncated at three standard deviations. The envelope at each contact was baseline corrected over [-50, 0) ms and negative values were clipped to zero. Population-specific response magnitude was quantified as the mean baseline-corrected MUA envelope amplitude across recording contacts and time samples within the half-open [0, 25) ms post-stimulus window. For each extracellular signal representation and stochastic input realization, the response magnitude of each cortical population class was divided by the summed response magnitudes of all ten population classes and multiplied by 100, such that normalized contributions summed to 100% within each realization. Normalization was therefore performed before across-realization averaging, preventing realizations with larger absolute extracellular responses from receiving greater weight in the population percentages. For each population and signal modality, normalized contributions were subsequently averaged across the 25 stochastic background-input realizations within each condition and are reported as arithmetic mean ± SEM.

For each signal and condition, we report normalized percentages, absolute response magnitudes, individual-realization distributions and mean ± SEM across the 25 background-input realizations. The ten cortical classes are a subset of the 43 native populations. Their normalized MUA-like envelope magnitudes are not an additive decomposition of the whole-network MUA because rectification and clipping are nonlinear, even though the underlying population voltages sum linearly. Comparisons across conditions are descriptive: connectivity and input weights differ together, and realizations are not paired across states. No significance tests were performed for the population-contribution comparisons.

### Device simulation & multi-column organizations

The DiSc and UPROBE devices (**Fig. 1D**) were modeled in ANSYS Electronics Desktop (AEDT, version 2025b) to achieve a quasistatic lead field using finite element modeling. Both device simulations had identical environments and material constants. A 15 mm edge cube of conductive, isotropic “brain” tissue was used with a conductivity of 0.25 S/m and edges defined as a 0 V reference. Both probes had identical properties of insulating probe bodies (10^-10^ S/m) and conductive contacts (10^9^ S/m) but varied probe characteristics as described in **Table 1**. With completed models, quasistatic field solutions were achieved, exported on a 100 μm voxel scale, and loaded into Python for source simulation as described in our prior work^12,29^. Uprobe, both virtual and physical, consists of 23 laminar contacts in a single column while the virtual DiSc consists of 8 equally spaced laminar columns of 16 contacts (128 contacts total) with a half-pitch offset for every other column, creating a diamond pattern. This offset creates two vertical pitches with different starting points, resulting in a 15.5x pitch for total contact span. The single-column pitch is 300 µm, while the neighbouring column creates a distributed pitch of 150 µm. The 128-contact virtual DiSc is not the physical 64-channel DiSc configuration used for experimental recordings (See Macaque data collection). Both virtual probes are shown in **Fig. 4A**.

**Table 1.** UPROBE and DiSc device characteristics.

|  | **UPROBE** | **DiSc (virtual)** | **DiSc (recorded)** |
|---|---|---|---|
| **Number of recording contacts** | 23 | 128 | 64 |
| **Probe diameter (µm)** | 450 | 800 | 800 |
| **Assumed probe damage radius (µm)** | 75 | 100 | 100 |
| **Contact dimensions (µm)** | 25 | 120 | 86 × 136 |
| **Contact area (µm<sup>2</sup>)** | 491 | 11,310 | 11,696 |
| **Contact vertical pitch (µm)** | 100 | 300 1-column;<br>150 multi-column | 600 1-column;<br>300 multi-column |
| **Contact vertical span (µm)** | 2,200 | 4,650 | 4,500 |
| **Contact horiz. pitch (µm; deg.)** | N/A | 314; 45° | 314; 45° |

Individual simulated A1 columns were loaded at cell-compartment resolution using transmembrane currents. After spatial orientation and placement relative to each device, these compartment-level current sources were combined with the corresponding FEM lead fields to compute device voltages for each cell. Voltage results were preserved at a cell type and layer resolution to enable additional investigation. Given 25 available realizations of each thalamic recruitment condition (Peak, Mid, etc.), the placement was randomized for each of 3 technical replicates per cortical region configuration. In each replicate, superposition of all individual columns was used to create the combined “tissue” state as well as accelerate computation. For **Supplement S9**, each UPROBE replicate was additionally rebuilt with the contact face rotated 90° about the shank axis to point toward the region 1 center. To keep this tractable, both orientations of all nine replicates were built from a fixed pool of 20 of the 100 column realizations, each column being assigned to a pool realization of its own condition, so that the two orientations of a replicate record identical columns and realizations and are compared pairwise (per-contact LFP pattern cosine similarity to the full-realization replicates 0.999, gain 0.90, n = 9).

The primary arrangements used a rigid grid without cortical-column rotation or tilt. We then tested randomized rotations and tilts (<5°) in three replicates against their unmodified counterparts. Average displacement of cells was roughly 1 voxel due to the comparative scale of voxels and cortical columns. Voltage correlations between perturbed and unmodified arrangements exceeded 0.98 in all three comparisons, indicating little sensitivity to these small geometric perturbations.

For the device analysis, LFP, broadband high frequency activity (BHA), and MUA were obtained from the device voltages using the 1-300 Hz, 80-150 Hz, and 300-5000 Hz bands, respectively, with fourth order Butterworth filters. LFP was additionally common average referenced across contacts, and MUA was rectified. CSD was computed from the LFP using central finite differences along cortical depth. Because the probes have different vertical contact spacing, we emphasize relative CSD patterns rather than absolute magnitudes. An alternative view, shown in **Fig. 4D**, demonstrates a method to artificially enhance the DiSc pitch further by spatial averaging between close electrodes of neighboring columns. This yields 31 interpolated contacts per column at a 150 μm pitch. Each interpolated column lies midway between its parent columns and due to the method of generation, all interpolated columns have the exact same depth span.

After formatting each region-replicate into a single vector of per-contact response amplitudes, cosine distances are computed between all replicate pairs for each device independently. Cosine distance is invariant to uniform scaling, isolating the spatial arrangement of signal relative to contacts which is a key distinguishing factor of source configurations ^33^. For each device, the resulting 9x9 matrix of pairwise replicate distances were computed between regions and averaged across replicates, forming a dissimilarity matrix embedded into two dimensions by metric multidimensional scaling (MDS; via SciKit-Learn)^34^ as seen in **Fig. 4E**. A separability measure was computed from the within- and between-region cosine distances for each device, taken as the mean distance over replicate pairs sharing a region and over pairs from different regions. Silhouette coefficients were computed from the same matrix labeled by region. The silhouette reports cluster tightness and separation jointly on a bounded range [-1,1], enabling direct comparison between devices, and establishes a basis for recording separability. The separability ratio compares the same quantities but with all replicates averaged before division, and is unbound above. Because regions 1 and 3 are mirrored about the probe while region 2 is centralized, an angular-radial index was made to test region (R) assignment by leave-one-out nearest-centroid classification under cosine difference (d) using *d* (*R* 1 *, R* 3)*÷ mean*( *d* (*R* 1 *, R* 2) *, d* (*R* 2 *, R* 3)).

### Macaque data collection

Extracellular electrophysiological recordings were obtained from the auditory cortex of two macaque monkeys. All recordings were obtained while the animals were awake and sitting quietly in a primate chair. All animal procedures, surgical preparation, and experimental protocols were conducted under protocol AP2023-728, approved by the Institutional Animal Care and Use Committee of the Nathan Kline Institute, in compliance with the Guide for the Care and Use of Laboratory Animals (National Research Council of the National Academies).

Recordings were acquired using two complementary recording geometries: DiSc and UPROBE. The 0.8-mm-diameter DiSc probe contained 128 fabricated sites, of which 64 rectangular contacts (86 × 136 µm) were connected for recording and arranged in eight directional columns at 45° angular intervals, with a 600-µm vertical pitch within each connected directional column and 300-µm spacing between successive connected rows. This shares the same staggered half-pitch arrangement as the virtual DiSc: the fabricated sites have a 300-µm within-column pitch and 150-µm spacing across successive staggered rows. The 64 connected contacts span 4,500 µm vertically between the uppermost and lowermost contact centers, compared with the 4,650-µm span of the virtual 128-contact geometry. The analysis included six DiSc penetrations and eight conventional laminar UPROBE penetrations. Penetrations, rather than animals, were the units of the group analysis. Together, these recordings provided complementary measurements of extracellular activity using radial (DiSc) and linear laminar (UPROBE) sampling geometries. For UPROBE recordings, signals were referenced to saline in the recording chamber, preamplified 10-fold, analog band-pass filtered at 0.1-500 Hz for field potential and 500-5000 Hz for MUA, and the MUA signal was rectified before digitization. Both field-potential and rectified-MUA signals were sampled at 2 kHz using a customized National Instrument A/D and LabVIEW acquisition system.

During recordings with both electrode types, we inserted the probe from the lateral-parietal surface and advanced it in 2-mm steps (UPROBE) or 2–4-mm steps (DiSc). At each step, we examined responses to broadband noise. The probe was advanced until polarity inversion of response to the noise sound was observed. After recording, we assigned cortical layers offline using spatiotemporal CSD and MUA response profiles. We identified the granular layer at channels showing the earliest MUA responses and a concomitant CSD sink below the field-potential (FP) polarity inversion. We identified supragranular layers from current sink/source pairs flanking the FP polarity inversion. We assigned channels below the granular-layer sink to infragranular layers.

Acoustic stimuli comprised fourteen pure tones (354, 500, 707, 1000, 1414, 2000, 2828, 4000, 5656, 8000, 11312, 16000, 22624 and 32000 Hz) and broadband noise (BBN). Both devices used a repeated fixed playback sequence. DiSc event labels identified the presented stimuli directly; UPROBE event identifiers were converted to acoustic identities according to their positions in that sequence. All events, including incomplete final cycles, were retained. Frequencies are presented in ascending order in the figures. Frequency-dependent responses therefore remain associated with sequence position and preceding stimulus history. Stimuli lasted 50 ms at 60 dB SPL and were presented at approximately 624.5-ms intervals.

### Experimental data analysis

Experimental MUA was processed separately for the two probe systems. DiSc signals recorded at 20 kHz were notch filtered at 60 Hz, band-pass filtered at 300–3000 Hz, rectified and smoothed with a Gaussian kernel (σ = 5 ms); the analyzed signals were sampled at 2 kHz. UPROBE measurements were device-supplied rectified MUA sampled at 2 kHz and received no additional digital filtering or rectification. At each contact, response energy was the mean absolute amplitude during 0–50 ms minus that during −50 to −10 ms. Both interval endpoints were included, giving 101 response samples and 81 baseline samples. DiSc energies were averaged across per-trial differences; UPROBE energies were averaged across trials and time samples. Negative values were retained. DiSc amplitudes are reported in source units and UPROBE amplitudes in stored counts; these units do not support absolute amplitude comparisons between devices. UPROBE stimulus-event times were identified from the recorded trigger signals, using MUA trigger onsets when necessary to ensure correct trial alignment.

### DiSc multidimensional response organization

For each of six DiSc penetrations, fifteen MUA energy maps included all 64 contacts. Channel index = 8 × direction index + depth index, represented as an 8 × 8 depth-by-direction matrix, with each index ranging from 0 to 7. Physical angular origin and absolute cortical depths were unavailable, so maps are shown in contact coordinates. Full frequency maps and direction profiles at a single depth selected from the mean tone response are provided in **Supplement S5**. Color limits were shared across stimuli within each penetration, without per-tone normalization.

BBN-versus-individual-tone Pearson correlations were computed across corresponding elements of the complete 64-contact maps. DiSc tone magnitude was the mean signed energy of the strongest 25% of finite contacts, rounded up and selected separately for each tone (16 of 64 contacts in every recording). Trial SEM for that magnitude is conditional on the selected contact set. Correlation intervals used 2,000 trial-bootstrap resamples within each stimulus category, independently resampling tone and BBN trials while holding contacts fixed. These within-penetration intervals differ from the penetration-bootstrap group intervals.

For each penetration, we calculated Pearson correlation between the BBN response profile and the elementwise mean of the fourteen tone profiles across all paired finite contacts. Group summaries report the arithmetic mean correlation and a 95% percentile interval from 10,000 bootstrap resamples of penetrations (NumPy default_rng, seed 0). A two-sided exact sign-flip test enumerated all 2^n^ sign combinations of Fisher-z-transformed penetration correlations under a zero-mean, sign-symmetric null. Penetrations were the analysis units; repeated penetrations from the same animal may be dependent, so these summaries do not establish independent animal-level replication. A clearly separated response maximum required b > 0 and (b − s)/|b| ≥ 0.20, where b and s are the strongest and second-strongest tone magnitudes. This criterion is descriptive, not an inferential test of frequency tuning. Illustrative penetrations were chosen from those meeting this criterion; all penetrations were retained in the group analysis.

### UPROBE laminar response organization

UPROBE analyses included all 23 neural contacts (indices 0–22) for each of eight penetrations. Correlations compared BBN with each individual tone profile and with the mean of all fourteen tone profiles. Tone magnitude was calculated within one contiguous region selected from the signed mean-tone profile and used for all tones in that penetration. The region contained 40% of finite contacts, rounded up (ten contacts in every recording), and maximized the sum of the mean-tone profile among fully finite contiguous windows; ties selected the first window. Tone magnitude was the mean of the nonnegative parts of the contact-mean energies within that region. Within-penetration correlation intervals were calculated by resampling trials within stimulus categories as described above, and magnitude uncertainty is reported as trial SEM conditional on the selected region and the set of contacts with positive mean energy. Individual trial energies were not clipped. These error bars exclude uncertainty in region selection and the clipping boundary. Full frequency-response profiles are provided in **Supplement S6**. The strongest tested tone describes preference within the tested set rather than a finely resolved characteristic frequency or registered tonotopic location.

### Modeling and analysis software

The https://github.com/TBBL-UTHealth/NetPyNE-DiSc-A1 repository contains finite-element models of the DiSc and UPROBE electrodes and associated simulation and analysis code used to examine the effects of recording geometry (**Fig. 4**). The https://github.com/NathanKlineInstitute/A1-DiSc-JNE repository provides the biophysical macaque auditory thalamocortical model, simulation configurations, and code and data for reproducing the analyses in **Figs. 2,3**. A compact, runnable version of the model is also deposited in https://modeldb.science/2041703 ^35^. The https://github.com/NathanKlineInstitute/A1-DiSc-NHP repository contains processed nonhuman primate (NHP) electrophysiological data and analysis code for reproducing the experimental results in **Fig. 5**.

## Results

### Biophysical auditory thalamocortical model links neuronal activity to extracellular signals

To investigate how controlled variations in thalamic input are transformed into extracellular electrophysiological signals, we used our biophysically realistic model of the macaque auditory thalamocortical system^18^ (**Fig. 1A**). Implemented in NetPyNE^29^/NEURON^30^, a framework for constructing and simulating data-driven multiscale neuronal networks, the model comprises a cortical column spanning layers 1-6 together with anatomically and physiologically constrained medial geniculate body and thalamic reticular populations. The model contains more than 12,000 neurons and approximately 25 million synapses and was previously shown to reproduce multiple experimentally observed features of macaque auditory thalamocortical activity, including spontaneous and evoked firing, laminar LFP/CSD organization, and oscillatory dynamics^18^.

**Figure 1.**
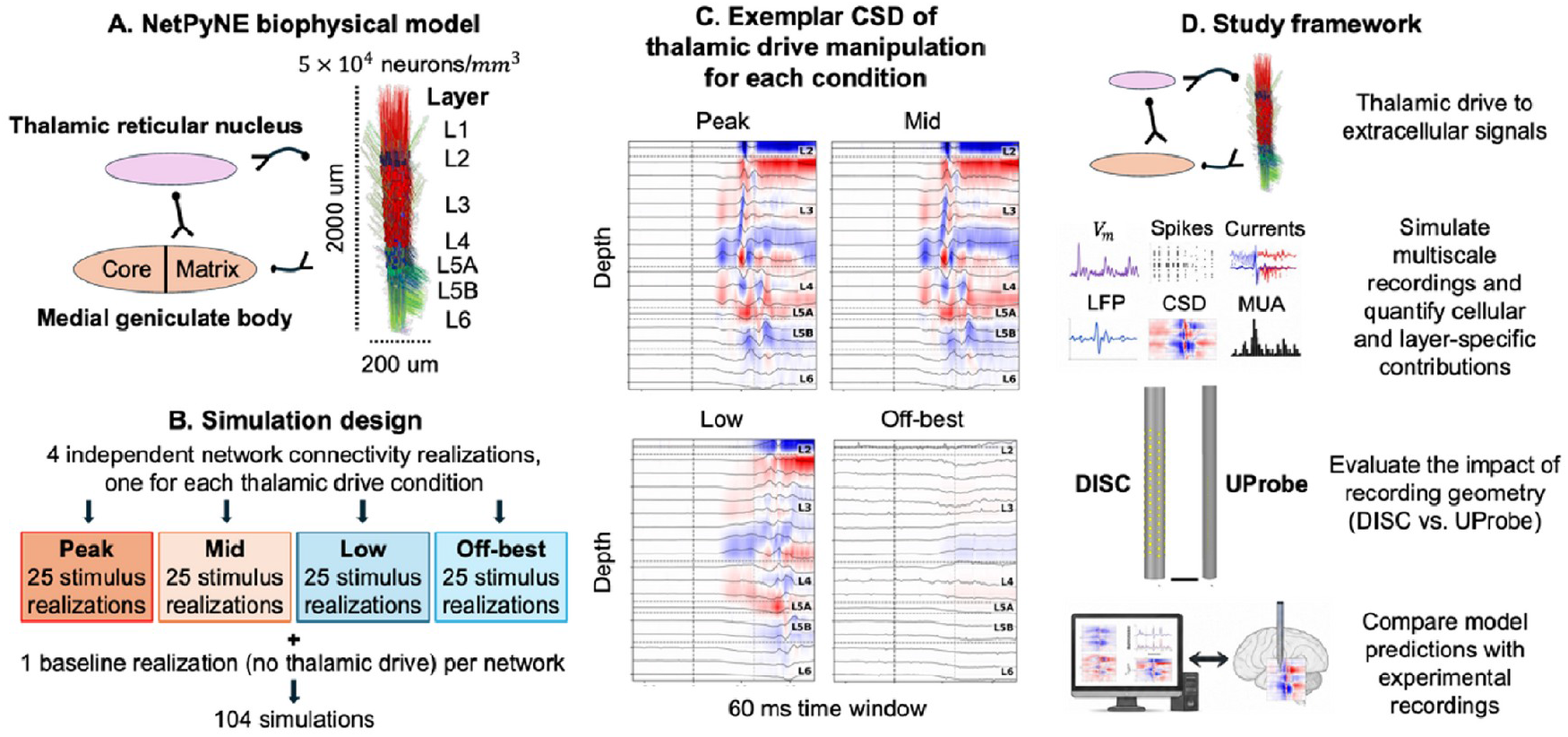
Computational and experimental framework. **(A)** Biophysical model of the macaque auditory thalamocortical system. **(B)** Four fixed cortical-column/drive conditions (Peak, Mid, Low and Off-best), each evaluated across 25 background-input realizations and one unstimulated reference, giving 104 simulation outputs. The “stimulus realizations” in **B** denote simulations with different stochastic background inputs; stimulus timing was deterministic. **(C)** Example laminar local field potential (LFP) and current-source density (CSD) responses. **(D)** Modeled membrane potentials, spikes and currents link neuronal activity to LFP, CSD and multiunit activity (MUA), virtual electrode recordings and experimental comparisons. Enlarged model, response and device illustrations are provided in **Supplement S1**.

To generate representative levels of cortical recruitment, we assigned four separately instantiated cortical-column realizations to progressively weaker levels of externally driven excitation delivered to the core TC and HTC relay populations, while input to the matrix TCM population remained fixed. The resulting operational conditions (Peak, Mid, Low, and Off-best) spanned progressively weaker thalamic recruitment (**Fig. 1C**). These states were used as representative components of an imposed multi-column response profile, with Peak denoting the most strongly recruited column and Mid, Low, and Off-best denoting progressively weaker recruitment. The labels do not represent separately simulated acoustic frequencies, empirically fitted tuning curves, or anatomically registered A1 locations. Each column/drive condition used a different connectivity realization, as expected for cortical columns at different locations, and was evaluated across 25 stochastic background-input realizations, together with one matched baseline without stimulus-driven thalamic input, yielding 104 analyzed outputs in total (**Fig. 1B**).

Each simulation provided simultaneous access to multiple levels of the same underlying network dynamics, including membrane potentials, neuronal spiking, transmembrane currents, LFP, CSD, MUA, and derived measures of extracellular response magnitude (**Fig. 1D**). This ability to derive extracellular observables from the same biophysical activity that generates cellular and population dynamics is a central advantage of detailed forward-modeling approaches, which provide a model-internal link between neuronal current sources and their resulting extracellular fields^1,15,36–38^. Because these quantities are known simultaneously within the simulation, differences in extracellular signals can be related to the specified column/drive conditions and to the neuronal populations generating them.

We therefore used the model not simply to reproduce a particular extracellular waveform, but as a mechanistic reference for the subsequent analyses. First, we asked whether the four column/drive conditions exhibited a consistent graded organization across complementary extracellular signals (**Fig. 2**). We then determined whether this common response organization arose from the same or different cellular contributors (**Fig. 3**), examined how multi-column cortical activity was transformed by distinct recording geometries (**Fig. 4**), and finally tested whether the resulting geometry-appropriate organizational features remained detectable in both DiSc and linear array laminar recordings (**Fig. 5**).

**Figure 2.**
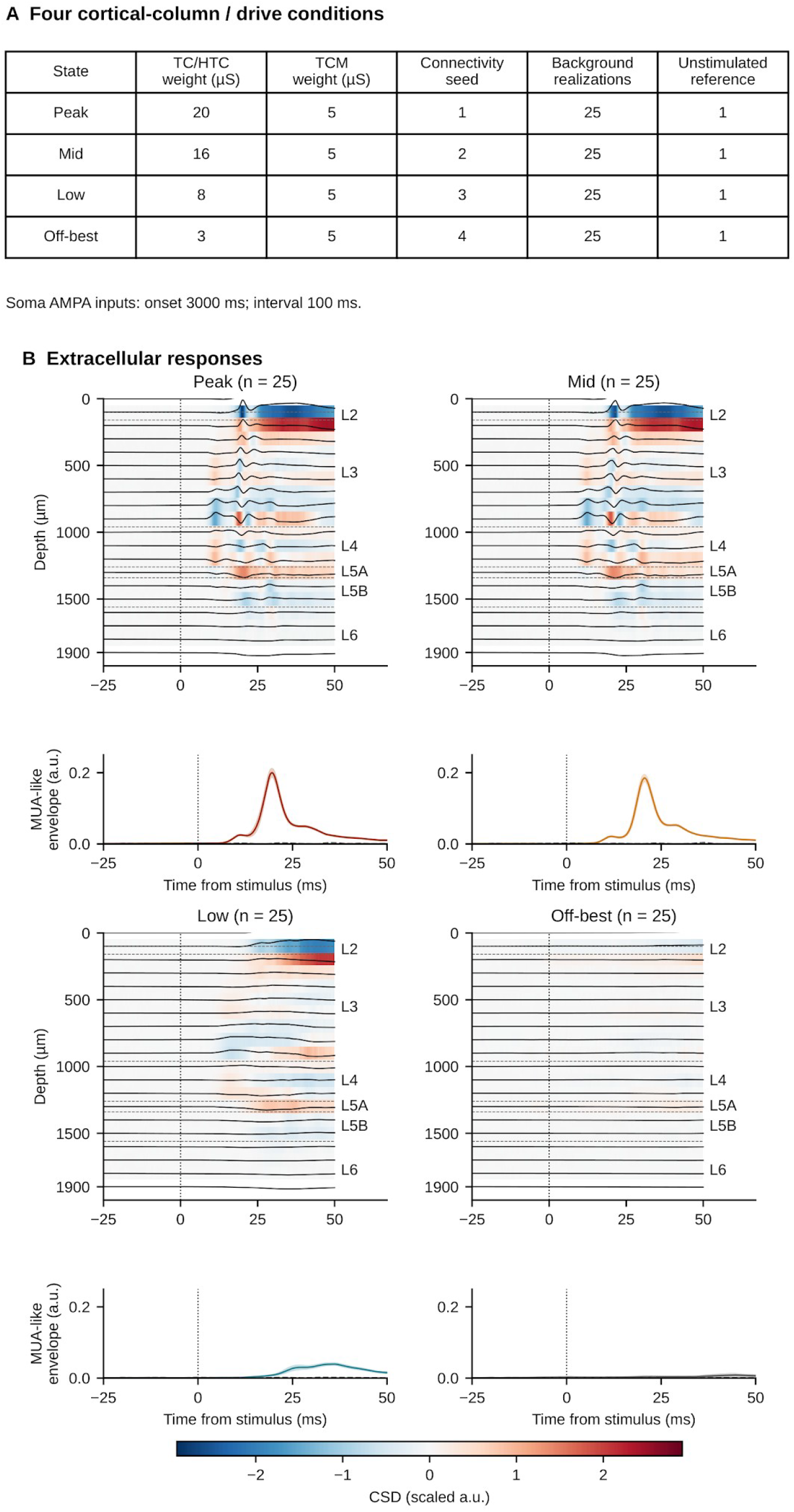
Extracellular responses across four cortical-column/drive conditions. **(A)** Thalamic input weights, fixed connectivity seeds and realization counts. Thalamocortical (TC) and high-threshold thalamocortical (HTC) populations received different input weights across conditions, whereas matrix thalamocortical (TCM) input remained fixed. **(B)** Mean responses across 25 background-input realizations per condition. Heat maps show baseline-corrected current-source density (CSD); black traces show local field potentials (LFPs) with a common scale. Lower traces show the model-derived multiunit activity (MUA)-like envelope averaged across all 20 contacts, with mean ± standard error of the mean (SEM). Gray dashed traces denote one separately processed unstimulated reference per condition. Contacts are spaced 100 µm apart over 1.9 mm; horizontal dashed anatomical guides use the model’s separate 2-mm cortical depth. Time zero denotes thalamic input onset. CSD and MUA-like amplitudes are shown in arbitrary units (a.u.). Thalamic spike rasters and rate profiles, the Peak–Mid contrast, and population rates are shown in **Supplements S2– S4**.

### LFP, CSD, and MUA preserve the imposed response hierarchy

To determine how complementary extracellular measurements represent the imposed column-response profile, we examined the four column/drive conditions across LFP, CSD, and MUA. Previous simultaneous recordings in macaque auditory cortex have shown that these signals can preserve common features of sensory organization while differing in their spatial extent and frequency selectivity. Differences among these signals reflect the distinct physiological and spatial processes represented by each signal: MUA predominantly reflects local suprathreshold population output activity, CSD estimates the local distribution of transmembrane current sinks and sources, whereas LFP can additionally contain contributions extending beyond its site of generation^4–6^. More recent laminar recordings have similarly demonstrated distinct spatial distributions of MUA, CSD, and field-potential activity across macaque auditory cortex, including broader field-potential distributions attributable in part to volume conduction^22,39^.

The four column/drive conditions exhibited a graded progression of extracellular response strength (**Fig. 2**). Representative simulations showed progressively larger stimulus-evoked responses from Off-best through Peak, whereas matched baseline simulations lacked comparable responses (**Fig. 2B**). Within this imposed response profile, the ordering of Peak, Mid, Low, and Off-best conditions was broadly preserved across LFP, CSD, and MUA. However, the expression of this common underlying recruitment differed substantially among signal representations, particularly in response polarity, temporal profile, spatial distribution, and waveform morphology. This dissociation between shared underlying population activity and its extracellular expression is consistent with recent computational analyses of macaque auditory cortex showing that similar inferred population dynamics can underlie substantially different laminar extracellular current distributions across recording sites and animals^22^.

This graded organization remained reproducible across stochastic background-input realizations. Averaging 25 stochastic background-input realizations within each column/drive condition produced consistent mean response profiles, with the corresponding SEM illustrating between-realization variability (**Fig. 2B**). The separation of the condition-averaged responses was maintained across realizations, with the Peak and Mid conditions producing the largest evoked responses, Low producing weaker responses, and Off-best remaining closest to baseline. This graded response pattern was evident across the simultaneously generated LFP, CSD, and MUA signals, although the magnitude and temporal profile of the LFP and CSD responses varied with cortical recording depth. These results extend the previous validation of the underlying macaque auditory thalamocortical model, which reproduced experimentally constrained firing, LFP, and CSD dynamics^18^, by showing that the imposed column-response conditions are represented coherently across several simultaneously generated extracellular observables.

The early MUA-envelope amplitude was 0.054263 ± 0.002382 a.u. for Peak and 0.047504 ± 0.002058 a.u. for Mid (mean ± SEM, n = 25 background realizations per fixed state). Peak minus Mid was 0.006759 a.u. (95% bootstrap CI [0.000612, 0.012721]; Hedges g = 0.598; two-sided Welch p = 0.03695). This quantifies a contrast between two fixed connectivity/drive states. Population spike-rate summaries do not show universal monotonicity across all classes; TCM and inhibitory responses provide counterexamples to a simple uniform recruitment account (**Supplement S4**).

Together, these results establish that the four condition-specific column simulations exhibit a graded hierarchy of cortical responses across the extracellular signal representations examined. Although this cross-signal ordering is expected because each signal was derived from the same activity within a given simulation, LFP, CSD, and MUA nevertheless differ in their biophysical basis and measured waveform structure. These differences motivate the subsequent analysis of how this shared response organization is shaped by the cellular populations contributing to each extracellular signal.

### Signal representations differ in population weighting

The biophysical model enables direct quantification of the population-specific contributions to each extracellular representation. We therefore quantified contributions from excitatory and inhibitory populations distributed across the cortical column during the 0-25 ms post-stimulus response window (**Fig. 3A-C**), providing a cellular decomposition that is difficult to obtain experimentally, where the neuronal populations contributing to extracellular signals must generally be inferred indirectly.

**Figure 3.**
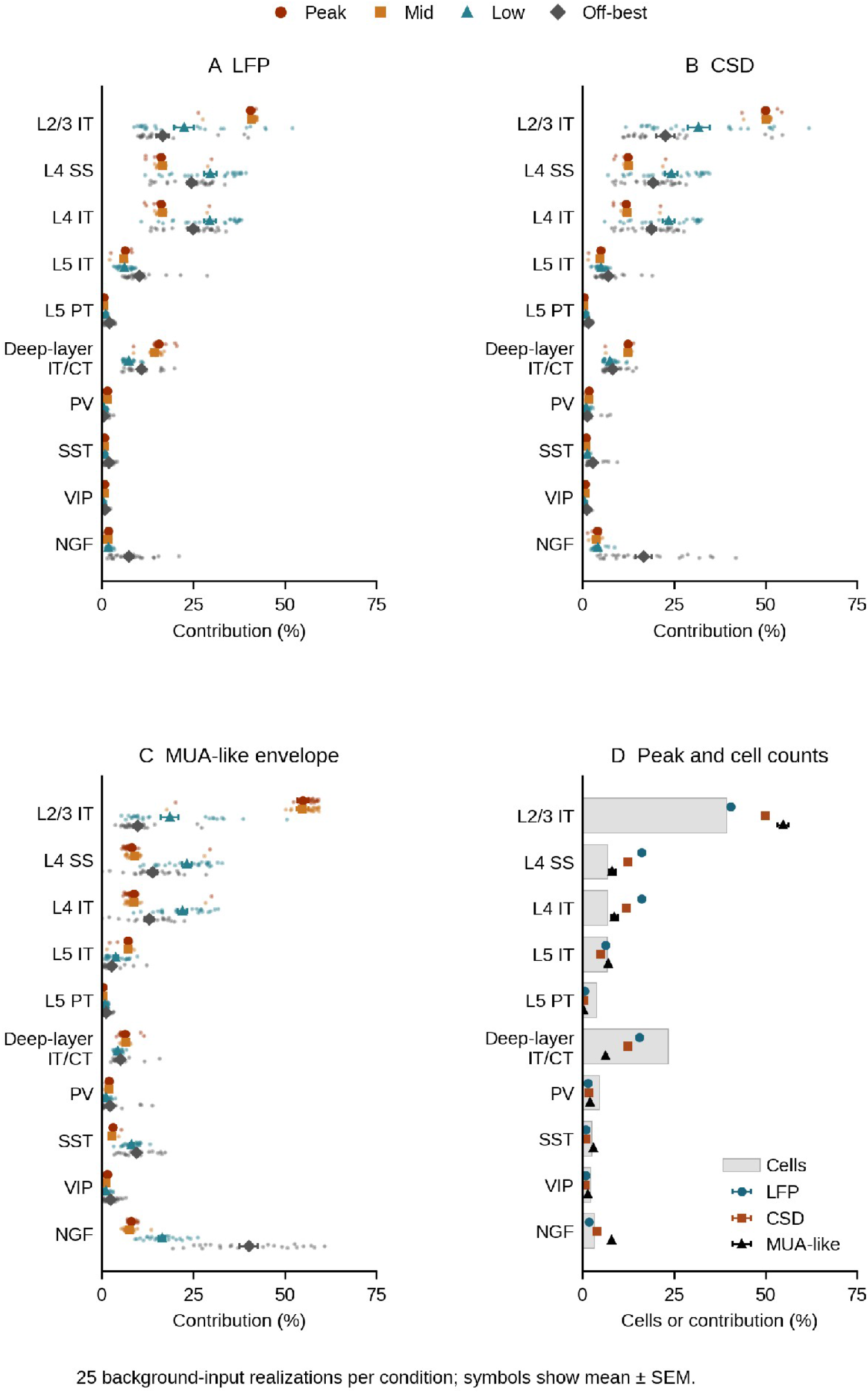
Population contributions across cortical-column/drive conditions. **(A-C)** Normalized contributions to local field potential (LFP), current-source density (CSD), and the multiunit activity (MUA)-like envelope for ten cortical classes in Peak, Mid, Low and Off-best. Small points show individual background-input realizations; larger symbols and horizontal error bars show mean ± standard error of the mean (SEM), with n = 25 in every condition. Response magnitudes were calculated over [0, 25) ms after baseline correction over [-50, 0) ms and normalized across the ten classes within each realization. **(D)** Cortical cell fractions (gray bars) and Peak signal fractions (symbols, mean ± SEM). The model contains 12,187 cells in these cortical classes. Native population voltages were summed within classes before signal processing. MUA-like percentages describe normalized envelope magnitudes, not an additive decomposition of whole-network MUA. Connectivity and input weights differ between conditions. Absolute magnitudes and all individual values are provided in **Supplement S8** and the accompanying data tables.

The resulting population profiles revealed a shared organization across signal modalities together with systematic differences in the relative weighting of individual populations (**Fig. 3A-D**). In Peak, LFP contributions were dominated by excitatory populations, particularly neurons in superficial and granular layers together with the aggregated deep-layer IT/CT group. This pattern is consistent with known effects of neuronal morphology and source-electrode geometry on LFP generation^1,2,5,36^.

While CSD approximation exhibited a broadly similar population organization, critically, it altered the relative weighting of individual populations (**Fig. 3B,D**). In Peak, superficial excitatory populations became more prominent relative to several granular and deeper-layer populations. This redistribution is consistent with spatial differentiation in CSD, which emphasizes localized extracellular gradients^3,6^.

In Peak, the MUA representation produced a further redistribution of population contributions (**Fig. 3C,D**). Superficial excitatory populations remained prominent, while the relative contributions of mixed deep-layer IT/CT and L5 PT decreased and several inhibitory contributions increased. The more local sensitivity of high-frequency extracellular activity provides a plausible basis for this altered weighting^1,4,39^, while broadband high frequency activity (BHA) may occupy an intermediate spatial regime^40^. The modeled MUA signal should therefore be interpreted as a high-frequency extracellular representation rather than a direct measurement of population spike counts.

The Peak comparison makes these modality-dependent redistributions particularly apparent (**Fig. 3D**). Although the same major cortical populations were represented across LFP, CSD, and MUA, their relative prominence changed systematically across signal modalities. Superficial excitatory populations became increasingly prominent across the modeled transformations (LFP → CSD → MUA), whereas several granular and deeper-layer excitatory populations became comparatively less prominent, and inhibitory populations showed distinct signal-dependent changes. Thus, the cross-signal transformation preserved shared population-level organization while changing the relative emphasis of specific populations. These percentages should be interpreted as model-derived relative contributions rather than direct quantitative estimates of each population’s contribution in local cortex.

Specifically, L2/3 IT comprises 39.38% of the 12,187 modeled cortical cells and contributes mean normalized magnitudes of 40.45% for LFP, 50.01% for CSD and 54.86% for the MUA envelope across the 25 Peak realizations. NGF comprises 3.25% of cells and 8.04% of normalized MUA-envelope magnitude. Comparisons retain L4 SS, L4 IT, L5 IT, L5 PT and mixed deep-layer IT/CT as distinct classes. Signal fractions are not simply proportional to cell counts; these descriptive comparisons do not isolate morphology, current distributions, timing or source-contact geometry as separate causes. The corresponding L2/3 IT MUA-like envelope fractions were 54.86% in Peak, 54.78% in Mid, 18.54% in Low, 9.76% in Off-best. Across the four conditions, L2/3 IT LFP fractions ranged from 16.57% to 41.02% and CSD fractions from 22.69% to 50.24% (**Fig. 3A–C**). These differences describe variation among the four fixed network/input conditions, rather than an isolated effect of thalamic drive.

Across signals, Peak and Mid had similar mean weighting profiles. L2/3 IT had the largest mean LFP and CSD fractions in both states, whereas L4 SS and L4 IT each accounted for about 30% of LFP in Low and 24-25% in Off-best. L2/3 IT remained the largest individual CSD class in Low and Off-best, although its mean fraction fell to 31.76% and 22.69%. MUA-like weighting changed more strongly: L4 SS and L4 IT were the largest classes in Low (23.20% and 22.01%), and NGF was largest in Off-best (40.16%). SST envelope fractions increased from 2.97% in Peak to 9.46% in Off-best, whereas PV and VIP remained below 2.5% in every state. L5 IT, L5 PT and mixed deep-layer IT/CT also differed by signal and condition, without a common monotonic ordering. The larger NGF envelope fraction in Off-best did not reflect a larger absolute envelope than Peak: its mean magnitude was 0.00169 versus 0.00643 model mV (**Supplement S8**).

Together, **Figs. 2-3** show that a common response ordering can coexist with signal-specific population weighting, motivating the next test of how that organization is transformed by recording geometry.

### Recording geometry determines the accessible spatial representation

Having established that complementary extracellular signals preserve common organizational features despite differing in the relative contributions of neuronal populations (**Figs. 2-3**), we next asked how those signals are transformed by the recording geometry itself. Extracellular potentials form spatially distributed voltage fields whose measured amplitude and structure depend on the position of the recording site relative to their neuronal generators^5,7,8^. Computational and physical models have further demonstrated that electrode dimensions, contact position, and recording-site geometry influence the potentials ultimately measured from an otherwise unchanged extracellular source^5,7,8^. Thus, an implanted electrode does not provide a geometry-independent readout of neural activity, but rather, it samples and spatially integrates the extracellular field according to the physical arrangement of its recording contacts.

To isolate this measurement transformation from biological variability, we constructed synthetic multi-column cortical configurations from independently simulated auditory cortical columns (**Fig. 4A,B**). Each column represented a complete biophysical simulation performed under one of four thalamic-recruitment conditions (Peak, Mid, Low, or Off-best), preserving its corresponding membrane dynamics, transmembrane currents, extracellular potentials, and spiking activity. Columns were placed on a 0.2 mm grid and assigned a recruitment condition according to their lateral offset from the center of peak recruitment, generating three region configurations with displacements of -1 mm, 0 mm, or +1 mm along the x axis. Each configuration was composed of 200 cortical columns for DiSc and 216 columns for UPROBE, using the same column trials between devices with additional columns to fill in the space around the smaller UPROBE device. Three technical replicates of each configuration were generated for each geometry by resampling the available simulated columns from a set of at least 25 per condition. Thus, the simplification introduced at this stage lies in the imposed spatial organization of otherwise biophysically simulated cortical columns rather than in their underlying neuronal dynamics. Though cortical columns are organized into active regions, each column is simulated in isolation resulting in no lateral connectivity.

**Figure 4.**
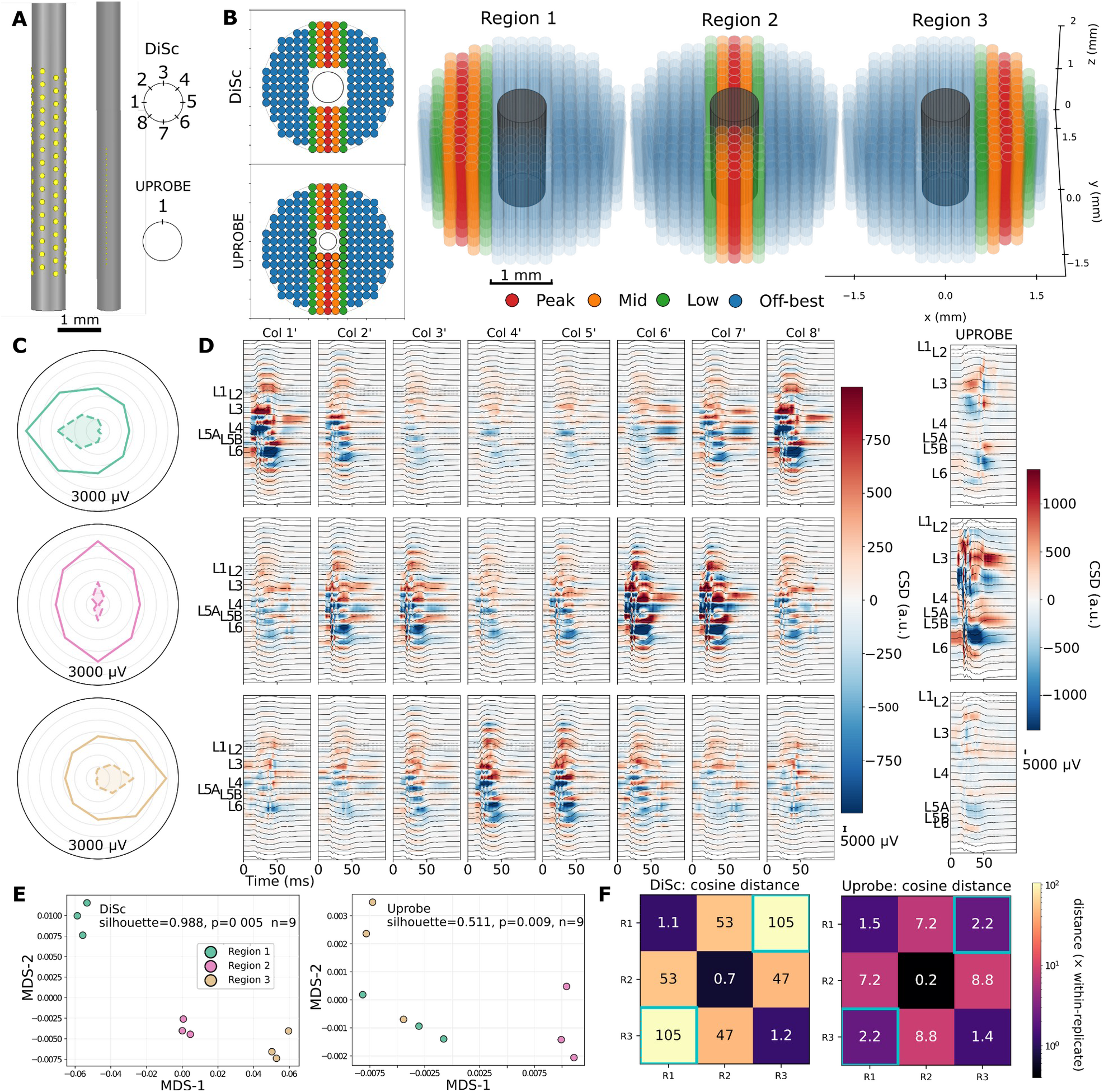
Recording geometry transforms common multi-column cortical activity into radial sensitivity in DiSc and laminar sensitivity in UPROBE. **(A)** Virtual device geometries. Left: side view of the 128-channel virtual DiSc and 23-channel UPROBE at a common scale (scale bar, 1 mm). Right: top-down (x-y plane) contact layouts of each denoted by dashes. The DiSc circular diagram defines the column numbering (Col 1-8, 45° apart) and the angular reference used in B-D. The UPROBE diagram shows its single column and direction of contact facing (+y, orthogonal to the region displacement axis; a-x-facing orientation comparison is available in **Supplement S9**). **(B)** Three idealized multi-column configurations constructed from independently simulated columns. Cortical columns are depicted with red, orange, green, and blue cylinders while the representative device shanks are in grey. The left side top-down view demonstrates the device sizes (black circles) and clearance regions. The right side is a front-top 45° view of the columns and DiSc device (trimmed length to fit in the tissue depth). **(C)** Directional response profiles for DiSc in each region. Each polar plot shows the wideband root mean square (RMS) voltage within the response window for each of the eight columns, plotted at that column’s angular position. Shading of the smaller area shows the minimum-subtracted response. Angular response is defined in A and radial units are µV. **(D)** Representative LFP traces and CSD maps for each device and region (x-axis time in ms; y-axis cortical depths marked by layer; LFP traces in µV; CSD in arbitrary units). For DiSc, between-column spatial averaging increased the displayed sampling from 16 to 31 laminar contacts per recording column and reduced effective pitch from 300 to 150 µm; interpolated columns are labeled Col N′. **(E)** Two-dimensional metric multidimensional scaling (MDS) of cosine distances among complete per-contact LFP amplitude vectors, using UPROBE and non-interpolated DiSc, across the three regions with three technical replicates. Axes are MDS dimensions 1 and 2 (unitless). **(F)** Region-pair cosine distances (multiples of the mean within-region distance) for DiSc and UPROBE, with confusion between regions 1 and 3 highlighted in teal. n = 3 technical replicates per configuration per device.

The same multi-column sources were then sampled using virtual implementations of two substantially different recording geometries: a conventional linear UPROBE with 23 contacts at a 100 μm pitch and the circumferential DiSc array with 128 contacts at a 300 μm vertical pitch and 8 staggered columns of 16 channels (**Fig. 4A-B**). The virtual UPROBE matches the physical probe while the simulated DiSc model provides twice the contacts and a longer depth span. Previous computational work with silicon microelectrodes has shown that changes in recording position and electrode configuration can alter the amplitude and detectability of extracellular neuronal sources^7^, while physical models of recording contacts predict spatial averaging of coherent extracellular potentials over the exposed electrode surface^8^. Recent finite-element analyses have further demonstrated that electrode placement and spatial configuration affect the information about cortical LFP sources recoverable by an array^12^. The additional 16 cortical columns in each UPROBE replicate lie close to the probe since our model assumes a larger damage radius for a larger shaft diameter (**Table 1**)^41^. As such, comparisons of absolute response amplitudes are not perfectly attributable to sampling geometry, but comparisons are valid between spatial patterns.

Representative recordings demonstrated that these two geometries transformed the same cortical activation patterns into distinct measured representations (**Fig. 4D-E**). Decomposing each DiSc array of per-contact response amplitudes into depth and directional components using a two-way sum-of-squares partition attributed 55.2% of its variance to depth, 31.9% to directionality, and 12.9% to their interaction, whereas UPROBE sampling on a single axis carried its entire variance along depth by definition. The directional component of DiSc varied with the active region, accounting for approximately 40.4% of the variance for the mirrored regions 1 and 3, but only 14.9% for a centered source peak in region 2. DiSc directional signals for regions 1 and 3 point to the center of each peak region within 1° and have standard deviations under 4°. Directional vectors are accordingly strong for regions 1 and 3, at 0.18 and 0.17, while the central region 2 is weaker at 0.03, supporting the directional modulation in amplitude. This directional sensitivity is consistent with the design principle of circumferentially distributed recording contacts, which provide multiple angular observations of the extracellular field around a common probe shaft^26^.

Separating each probe’s response to the three configurations into a gain component and a spatial-pattern component distinguished them further. Across regions UPROBE varied 1.98-fold in amplitude (13.1 times its within-region amplitude variability) with little pattern variation given by a between-region cosine distance of 0.012. DiSc amplitude varied by only 1.08-fold (1.8 times within-region variability) but the pattern variation was much higher at a between-region cosine distance of 0.084. The linear UPROBE encoded each configuration primarily in amplitude while DiSc encoded predominantly in spatial arrangement of signal across its contacts. This illustrates a broader principle already exploited by spatially oversampled and three-dimensional neural probes: the dimensionality and arrangement of recording sites determine which spatial components of neural activity are directly observable^10,23,24,42^.

The geometry dependence was not restricted to a single signal representation. Depth-dependent responses differed between DiSc and UPROBE across LFP, CSD, broadband high-frequency activity (BHA), and MUA frequency ranges, and the effect of cortical source location was expressed differently by the two devices in every band. Displacing peak recruitment changed UPROBE response amplitude by 2.0- to 3.6-fold depending on the band, compared to 1.1- to 1.4-fold for DiSc. Changes in the shape of depth profiles were larger for DiSc in all four bands, by 4.1 to 24.0 times the within-region distance, against 1.5 to 6.1 times for UPROBE. The relative sensitivity of the two geometries also varied across depth rather than differing by a constant factor. A fixed difference in lead field scaling would displace amplitude uniformly, but the DiSc-to-UPROBE ratio instead varied across depth within each band, by 2.2-fold for LFP, 16.4-fold for resulting CSD, 2.9-fold for MUA, and 3.6-fold for BHA. Contact spacing also directly impacts signal quality distribution and CSD computation. UPROBE has 21 of 23 contacts within the cortex while each DiSc column has only 7-8 of 16. As such, even after normalizing by the pitch differences, UPROBE recovers 2.1 times greater laminar current density than DiSc, indicating that the coarser 0.3 mm pitch attenuates the gradient beyond simple restoration.

To determine whether the complete multichannel representations retained information about the spatial organization, we exploited symmetry of regions 1 and 3 (**Fig. 4E-F**). A single-shank device geometry can only distinguish depth, leaving separation to be based on laminar pattern and magnitude. Cosine distances between the complete per-contact LFP arrays bore this out and quantified the value of directionality. For DiSc, the mirrored pair has the largest between-region cosine distance of 0.13 against 0.062 for the mean of the two pairs, and within-region distance mean of 0.0012. For UPROBE the ordering reverses such that the mirrored pair gives its smallest between-region distance, 0.0044, below its radial pairs at 0.016 and only 2.2 times its within-region distance mean of 0.0020. Expressed as the angular-radial index, this contrast is 2.11 for DiSc against 0.28 for UPROBE, a reversal in ordering rather than a difference in magnitude. Pooling all pairs, the mean between-region distance was 0.084 for DiSc and 0.012 for UPROBE, yielding a separability ratio of 68.4 for DiSc and 6.1 for UPROBE. In a leave-one-out nearest-centroid assignment of replicates, all 9 DiSc replicates were grouped correctly while UPROBE missed two, mistaking samples of the mirrored regions 1 and 3. Silhouette scores on the array data achieved 0.98 for DiSc and 0.51 for UPROBE. Because the UPROBE orientation was on the line of symmetry for regions 1 and 3, rotation of the contact face toward region 1 to create a non-symmetric arrangement was explored in **Supplement S9**. Rotation redistributed gain between the lateral regions (rotated / original LFP gain 1.33 in region 1, 0.80 in region 2 and 0.88 in region 3; mirror-pair gain ratio 0.64 versus 0.97) but left the laminar pattern unable to resolve the mirrored sources, which remained the closest pair of regions with an unchanged silhouette (0.47 versus 0.48 within the same comparison), whereas DiSc separates the mirrored regions without a preferred orientation.

Together these findings show that DiSc and UPROBE preserved different spatial components of the same multicolumn source activity, motivating the subsequent test of geometry-appropriate organization in experimental recordings (**Fig. 5**).

**Figure 5.**
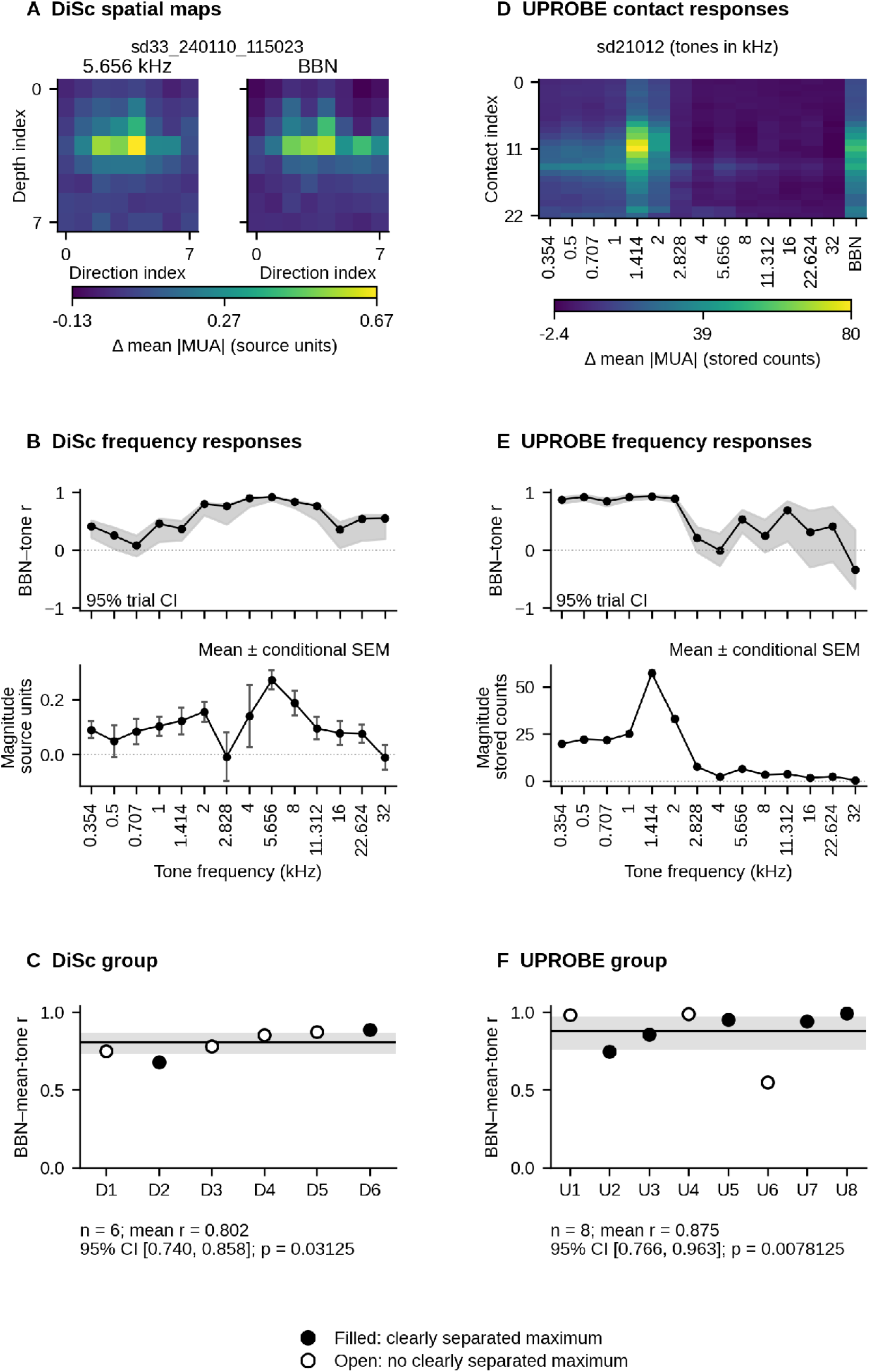
Spatial organization of tone- and noise-evoked multiunit activity (MUA). **(A)** Response maps for the strongest tested tone (5.656 kHz) and broadband noise (BBN) in DiSc penetration D6 (sd33_240110_115023). The 64 contacts of the recorded DiSc span eight direction and eight depth indices; both maps share a color scale. **(B)** BBN–tone correlations across all 64 contacts (upper) and mean signed response magnitude across the strongest 16 contacts selected separately for each tone (lower). **(C)** BBN– mean-tone correlations across six DiSc penetrations. **(D)** Tone and BBN response energies across 23 contacts in UPROBE penetration U5 (sd21012), whose strongest tested tone was 1.414 kHz. **(E)** BBN–tone correlations across all 23 contacts (upper) and tone magnitudes within ten contiguous contacts selected from the mean-tone profile (lower). (F) BBN–mean-tone correlations across eight UPROBE penetrations. Both illustrative penetrations met the separation criterion; all penetrations were retained in group summaries. In **B** and **E**, shading shows 95% trial-bootstrap confidence intervals and error bars show trial standard error of the mean (SEM), conditional on the selected contacts and, for UPROBE, the positive-mean contact set (Methods). In **C** and **F**, filled circles indicate a clearly separated strongest-tone response, b > 0 and (b − s)/|b| ≥ 0.20 (**Methods**); open circles indicate that this criterion was not met. Lines and shading show group means and 95% intervals from 10,000 penetration-bootstrap resamples; p values use exact two-sided Fisher-z sign flips. Clearly separated maxima occurred in 2/6 DiSc and 5/8 UPROBE penetrations. Response energy is baseline-subtracted mean absolute MUA amplitude. DiSc values use source units and UPROBE values use stored counts; amplitudes are not calibrated across devices. Coordinates are contact indices. Frequencies are displayed in ascending order, although stimuli were presented in a fixed, repeated sequence.

### Experimental recordings show geometry-appropriate organization

To determine whether the geometry-dependent forms of response organization identified in the simulations were detectable experimentally, we analyzed MUA responses obtained with recorded DiSc and linear array laminar probes (**Fig. 5**). Frequency-dependent spatial organization is a well-established property of primate auditory cortex: electrophysiological mapping has demonstrated systematic tonotopic gradients across primary and neighboring auditory fields, while laminar recordings reveal structured depth-dependent patterns of synaptic and spiking activity within individual cortical penetrations^39,43–46^. We therefore asked whether these established forms of auditory cortical organization were expressed in the geometry-appropriate representations predicted by the simulations: multidimensional organization across depth and circumferential direction for DiSc, and laminar organization along cortical depth for UPROBE. Importantly, these analyses tested preservation of organizational structure across heterogeneous penetrations rather than point-by-point reproduction of individual simulated responses.

The six DiSc penetrations exhibited substantial variability in tone-evoked response organization (**Supplement S5**). A penetration with a clearly separated response maximum at 5.656 kHz is shown in **Fig. 5A,B**. Tone-response magnitude was quantified from the mean MUA response across the strongest 25% of contacts for each tone, and the “site-preferred” (highest-response) tone was considered clearly separated when its separation from the second-highest response was at least 20% of the highest response (**Methods**). This classification therefore reflected the relative separation of the two strongest tone responses, rather than absolute response magnitude or broadband-noise/tone similarity. Some penetrations contained a clearly separated highest-response tone, whereas others showed several frequencies with comparable response magnitudes. Such variability is compatible with the known organization of primate auditory cortex, in which local neuronal populations exhibit frequency selectivity within broader tonotopic gradients rather than identical tuning across recording locations^39,43,46^. Because each DiSc recording preserves both depth and circumferential dimensions, we quantified broadband-noise/tone correspondence by correlating the complete 8 × 8 MUA energy maps rather than comparing response magnitude alone. This allowed us to test whether broadband noise recruited spatial organization shared across tone-evoked responses despite differences in frequency-dependent response magnitude.

Across the six DiSc penetrations, BBN maps were positively correlated with the mean of the fourteen tone maps (mean Pearson r = 0.802201; 95% penetration-bootstrap CI [0.739651, 0.857810]; exact two-sided Fisher-z sign-flip p = 0.03125; Fig. 5C). Two penetrations had clearly separated response maxima. These comparisons used all 64 contacts and indicate shared depth-by-direction response organization across noise and tones.

UPROBE recordings provided a complementary test along cortical depth. Laminar CSD and MUA recordings have long been used to characterize the sequence and distribution of auditory cortical activation because simultaneous sampling across layers resolves depth-dependent input and population-spiking profiles^39,45,46^. The eight penetrations differed substantially in strongest tested-tone identity and in whether a single tone produced a clearly separated response maximum (**Supplement S6**). **Figure 5D,E** shows penetration U5, which exhibited a clearly separated response maximum at 1.414 kHz and frequency-dependent contact profiles. Nevertheless, correlations between broadband noise and individual tone profiles were frequently high even when response magnitudes differed across frequencies. We therefore compared the broadband-noise depth profile with the mean profile across tones to test whether spectrally distinct stimuli recruited a shared laminar response organization.

Across the eight UPROBE penetrations, broadband-noise depth profiles were consistently correlated with the corresponding mean tone-evoked profiles (mean Pearson *r* = 0.874893, penetration-bootstrap 95% confidence interval [0.765694, 0.963257]; exact two-sided Fisher-z sign-flip *p* = 0.0078125; **Fig. 5F**). Five penetrations (U2, U3, U5, U7 and U8) had clearly separated strongest-tone responses. High correspondence was again observed in penetrations with both clearly separated and ambiguous highest-response tones. Thus, the laminar distribution of MUA was substantially preserved across broadband-noise and tone stimulation even when frequency dependence and absolute response magnitude varied across penetrations. This result is consistent with previous laminar response profile studies demonstrating reproducible depth-dependent organization of auditory responses despite variation in tuning, stimulus condition, and recording location^22,39,46^.

Together, the experimental recordings showed positive correspondence between BBN and mean-tone spatial responses for both probe geometries. DiSc retained variation across direction and depth, whereas UPROBE represented variation along its contact axis. Together, these findings demonstrate that both DiSc and UPROBE recordings capture shared spatial organization of auditory cortical responses to broadband noise and pure tones, with DiSc revealing depth- and direction-dependent structure and UPROBE revealing laminar organization.

## Discussion

Our results support a two-stage account of how cortical activity becomes an extracellular measurement. First, the imposed Peak-to-Off-best ordering was retained across LFP, CSD, and MUA even though the signals differed in waveform structure and population weighting (**Figs. 2–3**). Second, sampling the same multi-column sources with different virtual arrays produced geometry-specific representations: UPROBE emphasized laminar variation, whereas DiSc additionally captured circumferential source organization and more clearly separated mirrored configurations (**Fig. 4**). Macaque recordings showed corresponding geometry-appropriate organization, with broadband-noise responses correlated with mean tone-evoked maps for DiSc and laminar profiles for UPROBE (**Fig. 5**). These findings identify recording geometry as a second transformation, following biophysical signal generation, that determines which dimensions of cortical activity remain observable.

### Biophysical models provide mechanistic access to hidden neural processes

A central challenge in interpreting extracellular electrophysiology is that the measured signal is separated from its cellular generators by multiple levels of biological organization. LFPs arise from the superposition of transmembrane currents distributed across neuronal populations, while their magnitude and spatial extent depend on neuronal morphology, synaptic placement, synchrony, and the relative position of current sources and recording sites^1,2,5^. CSD provides a more spatially localized estimate of current sinks and sources, but remains a derived representation of population transmembrane activity, whereas MUA predominantly reflects nearby suprathreshold spiking^3,4,6^. Consequently, no single extracellular measurement uniquely specifies the cellular processes that generated it.

Biophysically detailed models provide one way to bridge this gap by explicitly linking experimentally constrained cellular and circuit properties to simultaneously generated physiological observables. Forward-modeling frameworks such as VERTEX and LFPy have demonstrated how transmembrane currents generated by detailed neurons and network models can be translated into extracellular potentials across arbitrary virtual recording sites^14–16,29^. Large-scale mechanistic models extend this strategy by integrating experimentally constrained morphologies, intrinsic conductances, synaptic dynamics, connectivity, and long-range inputs within a common network whose membrane activity, spiking, LFP, CSD, EEG, and related signals can be interrogated simultaneously^18,36,47^. This level of access transforms the model from a device for reproducing experimental traces into an explicit set of hypotheses about the chain of transformations linking circuit input, neuronal dynamics, extracellular field generation, and measured activity.

The present study uses that capability to examine four separately instantiated cortical-column conditions rather than fitting separate models to LFP, CSD, and MUA. Each condition combined a fixed connectivity realization with a prescribed level of externally driven thalamic recruitment, representing cortical columns at different locations. Stochastic background input was varied across repeated simulations within each condition, to represent trial-to-trial variability. Because each condition used a different connectivity realization, the Peak-to-Off-best hierarchy characterizes the imposed column-response conditions rather than an independent estimate of thalamic-drive effects across replicated networks. The expected ordering of response magnitude served as a controlled internal reference rather than the principal finding: the informative result was that this ordering remained identifiable across LFP, CSD, and MUA despite differences in waveform structure and population weighting (**Figs. 2-3**). In this respect, our results extend the previous validation of the macaque auditory thalamocortical model, which reproduced experimentally constrained spontaneous and evoked dynamics^18^, by using the model as a mechanistic reference in which the intermediate variables linking thalamic input to extracellular activity remain observable.

### Shared organizational features emerge despite distinct signal-generation mechanisms

The preservation of a common recruitment hierarchy across LFP, CSD, and MUA does not imply that these measurements are physiologically equivalent. Simultaneous recordings in macaque auditory cortex have previously demonstrated that MUA, CSD, and LFP can share frequency-related organization while differing markedly in spatial extent, with MUA generally exhibiting the most localized representation and LFP the broadest^6^. More recent laminar recordings likewise demonstrate substantial differences in the spatial distribution of spiking, current-source density, and field-potential activity across auditory cortical depth^39,40^. These experimental observations are consistent with biophysical modeling showing that extracellular field generation depends on the morphology and membrane properties of the contributing neuronal populations^36^, and that different laminar populations can contribute very differently to the field recorded at a given location^15^.

Our population-resolved simulations extend this literature by showing that similar macroscopic recruitment structures can persist despite differences in the relative weighting of the cellular populations contributing to each signal (**Figs. 2, 3A-D**). Within Peak, population-specific contributions differed across LFP, CSD, and MUA, with changes in the relative representation of granular, deeper-layer excitatory, and inhibitory populations across signal modalities (**Fig. 3D**). This result is consistent with the known influence of neuronal morphology, active membrane currents, and population geometry on extracellular field generation^1,2,36^. At the same time, it shows that different cellular mixtures need not lead to qualitatively different representations of network recruitment. The population percentages should therefore be interpreted as model-derived relative contributions rather than direct quantitative estimates of the contribution of individual neuronal populations in local cortex.

The four-condition comparison further shows that population weighting depends on network state as well as signal definition. Across Peak, Mid, Low and Off-best, the mean L2/3 IT MUA-like fraction ranged from 9.76% to 54.86%. The largest mean envelope fraction shifted from L2/3 IT in Peak and Mid to L4 SS in Low and NGF in Off-best. Absolute signal magnitudes and normalized fractions answer different questions: a larger fraction need not indicate a larger absolute response. Because each condition combines a different connectivity realization with a different input strength, these observations cannot identify either factor as the sole cause of the weighting differences.

These signal-dependent redistributions are consistent with the distinct biophysical and analytical transformations applied to each representation. LFP magnitude and spatial structure depend on neuronal morphology, synaptic organization, active membrane conductances, and source–electrode geometry^1,2,5,36^. Spatial differentiation in CSD emphasizes localized gradients in extracellular potentials and can therefore alter the relative weighting of population contributions^3,6^. The high-frequency MUA envelope preferentially emphasizes local fast activity associated with neuronal firing^1,4,39^, whereas broadband high-frequency activity (BHA) may occupy an intermediate spatial regime with a volume-conducted component^40^. Consequently, populations contributing to more than one signal need not retain the same relative prominence after transformation from LFP to CSD or MUA.

That stronger imposed thalamic input produced larger responses was expected. The important result was that the common ordering remained evident after distinct biophysical and analytical transformations, even as polarity, temporal profile, spatial distribution, and population weighting differed across LFP, CSD, and MUA (**Figs. 2-3**). Concordance among these signals therefore should not be taken to mean that they report the same biological process. Rather, different transformations can preserve a common lower-dimensional feature of network organization, while redistributing the contributions of individual populations. Comparable ideas have emerged from auditory modeling studies in which similar inferred population dynamics could underlie markedly different extracellular current distributions across cortical sites and animals^22^. The relevant correspondence can therefore occur at the level of network organization rather than waveform identity or cellular-generator identity.

This observation provides an important bridge to the subsequent device simulations. If common circuit organization can survive the first transformation, from cellular activity into distinct extracellular signals, the next question is whether it also survives a second transformation imposed by how those extracellular fields are physically sampled.

### Recording geometry constitutes a second transformation of extracellular signals

The dependence of neural recordings on electrode geometry is well established. Computational studies have shown that the amplitude and detectability of extracellular sources depend on the location of neurons relative to recording contacts and on electrode configuration^7^. Physical models further demonstrate that finite recording contacts spatially average the extracellular voltage field over their exposed area, meaning that contact size itself can alter the measured representation^8^. More generally, the spatial reach of LFP depends strongly on source organization and measurement location^5^. These observations establish that the measured signal cannot be considered independent of recording-device geometry.

Modern neural-interface development has increasingly exploited this relationship. Closely packed silicon probes use spatial oversampling to capture the extracellular footprint of neighboring sources^10^, while high-channel-count arrays and three-dimensional probes extend observations across increasingly large and multidimensional tissue volumes^23,24^. High-density Neuropixels probes similarly sample activity across extended cortical depth, and recent ultra-high-density implementations demonstrate that finer spatial sampling can improve the detection and spatial identification of extracellular neural sources^9^. The DiSc design approaches the same problem from a complementary direction by distributing recording contacts circumferentially around a depth electrode to recover directional information that would not be directly available from contacts arranged only along a linear axis^26^. Broader reviews of emerging electrode technologies similarly emphasize that increasing spatial integration and changing probe geometry fundamentally expands the dimensions of neuronal activity that can be observed^25^.

Our device simulations isolate this measurement transformation from the biological source more explicitly than is generally possible experimentally (**Fig. 4**). The DiSc and UPROBE virtual devices sampled the same multi-column transmembrane-current sources, meaning that differences in their outputs cannot be attributed to different neuronal dynamics. Under these controlled conditions, the linear probe preferentially retained laminar variation along depth, whereas DiSc additionally retained circumferential information about lateral source organization. The distinction should therefore not be framed as one geometry recording a more “accurate” version of the cortical activity than the other. Rather, the two geometries provide different projections of the same underlying spatial field, making them complementary for addressing different questions in the auditory system. Linear probes are particularly suited to resolving laminar response organization and propagation across cortical depth, whereas DiSc provides additional access to lateral spatial organization surrounding the recording site (**Fig. 4**). Modeling provides a means to interrogate both representations against the same underlying cortical activity and thereby determine which features of that activity each recording geometry preserves.

This interpretation is consistent with recent work treating electrode placement and configuration as an information-recovery problem. We recently demonstrated^12^ that the information available from cortical LFP sources depends systematically on electrode positioning and sensitivity profiles, while spatially oversampled arrays likewise exploit redundancy across neighboring contacts to improve source discrimination^10^. The present work extends this principle from electrode placement within a single general geometry to a comparison between fundamentally different sampling geometries. Similar considerations may apply to studies of transient sensory responses, where alternative recording configurations could provide access to additional spatial or population-level features of the underlying neural activity. The important quantity is therefore not simply signal amplitude or signal-to-noise ratio, but which dimensions of the underlying cortical organization remain identifiable after spatial sampling. Isolating gain and spatial pattern contribution for each geometry highlights the difference that UPROBE observes differences primarily by amplitude while DiSc operates primarily by spatial distribution across contacts. Because DiSc encodes by spatial pattern in addition to amplitude, mirrored cortical sources that are degenerate along an axis remain separable, with angular-radial indices of 2.11 versus 0.28 and overall separability ratios of 68.4 versus 6.1, resulting in silhouette scores of 0.98 versus 0.51 (**Fig. 4E-F**). Turning the UPROBE contact face toward one lateral source changes its gain for that source but not the separability of its laminar pattern (**Supplement S9**), so contact direction alone does not substitute for circumferential sampling.

The geometry dependence also varied across LFP, CSD, BHA, and MUA representations, suggesting that recording geometry and biological signal generation should not be considered independent stages in practice. Different extracellular components possess different spatial scales^6,19^, and a given contact arrangement will therefore interact differently with each of them. Measurement geometry can consequently alter not only where activity appears strongest but also which organizational features are emphasized by the resulting multichannel representation.

### Experimental recordings support organizational rather than point-by-point model validation

The experimental recordings were intended as a test of this organizational framework rather than as an attempt to reproduce individual simulated traces. This distinction is particularly important in auditory cortex, where systematic organization coexists with substantial local variability. Macaque auditory cortex contains established tonotopic gradients across primary and neighboring auditory fields^43,44^, while individual penetrations exhibit heterogeneous frequency preference and response magnitude. Laminar organization is likewise reproducible at the population level while varying across cortical location, stimulus condition, and preparation^39,46^. Tone and noise stimulation additionally recruit overlapping but non-identical neuronal populations; in primate auditory cortex, noise can evoke stronger or broader responses than pure tones while retaining relationships to local characteristic frequency^48,49^.

The UPROBE recordings provided experimental support for the model-derived framework along cortical depth (**Fig. 5**). Broadband-noise depth profiles were strongly correlated with the mean tone-evoked profile across all eight penetrations (mean *r* = 0.87), including penetrations without a clearly separated highest-response tone. Thus, stimulus identity and response magnitude could vary while a common laminar organization remained detectable. This result is consistent with prior macaque studies showing stable laminar organization across sensory-evoked responses^39,46^ and with modeling work showing that comparable population dynamics can persist despite differences in the precise extracellular current distributions measured across sites^22^.

The DiSc recordings provided complementary support for the model-derived framework across a multidimensional spatial representation (**Fig. 5**). Broadband-noise maps were consistently correlated with the mean tone-evoked spatial map across the six penetrations (mean *r* = 0.80), including penetrations both with and without a clearly separated highest-response tone. Thus, despite heterogeneity in frequency-dependent response magnitude and the degree to which individual tones dominated the response, shared depth × circumferential organization remained detectable across stimulus conditions. This correspondence extends the experimental comparison beyond laminar organization by showing that multidimensional spatial structure captured by DiSc can likewise be preserved across spectrally distinct stimuli.

The heterogeneity observed across penetrations is important for how validation of complex neural models is framed. Biological circuits can vary substantially across animals and preparations while maintaining related functional outputs, and multiple parameter combinations can produce similar network activity^19,20,50,51^. Accordingly, the experimental analyses targeted higher-order organizational features rather than one-to-one correspondence between an individual simulated column and an individual cortical penetrations. This does not mean that any model reproducing a broad qualitative feature should be considered valid. Rather, evaluation should instead target explicitly defined observables at the level appropriate to the scientific claim.

Formal work on model validation in computational neuroscience makes a similar distinction. Gutzen *et al.*^21^ argue for validating network models using statistical descriptors of population dynamics rather than requiring identity of individual trajectories, while emphasizing that the relevant validation metric must be specified according to the model’s intended scope. In the present study, the relevant observables were deliberately defined at the level of graded recruitment, laminar response organization, and multidimensional spatial structure. The experimental comparison therefore evaluates whether those particular relationships remain detectable under biological variability; it does not establish that the model uniquely reconstructs the cellular state of any individual penetration.

### Organizational features provide a useful but bounded framework for model validation

Biophysically detailed models inevitably combine experimentally constrained mechanisms with assumptions, approximations, and parameter choices. Even large-scale models containing detailed morphologies, conductances, synaptic dynamics, and connectivity cannot represent every biological feature of the modeled circuit^47^. Moreover, the existence of multiple parameterizations capable of producing related network behavior means that correspondence with one experimental observable does not uniquely identify the biological mechanism that generated it^19,20,50^. Model validation should therefore be interpreted in relation to a defined scope of applicability, rather than as confirmation that every internal parameter of the model is biologically correct. The growing literature on reproducibility and validation in computational neuroscience similarly emphasizes explicit model assumptions, comparison with experimental reference data, robustness across conditions, and validation at appropriate levels of network behavior^21,52^.

Our results suggest that organizational features can provide useful validation targets when exact waveform matching is neither expected nor scientifically necessary. The condition-specific simulation results showed that graded recruitment can remain recognizable despite different signal generators, while the device simulations showed that spatial organization can remain recognizable after geometry-dependent transformation. The experimental analyses then asked whether heterogeneous recordings exhibiting substantial variation in frequency-dependent response magnitude and tone-response dominance expressed organizational relationships consistent with those identified in the simulations. This hierarchy provides a more constrained claim than simply asserting that simulated and experimental recordings “look similar”: the model generates explicit relationships, and the experimental data assess whether corresponding forms of organization are evident under biological and measurement variability, without implying equivalence or the absence of potentially meaningful differences.

At the same time, organizational agreement cannot by itself establish a unique mechanistic explanation. Different mechanisms can produce similar outputs^19,20,53–55^, and different recording geometries can transform distinct underlying source distributions into partially similar measurements. The utility of organizational validation therefore depends on combining multiple constraints. In the present study, those constraints include the simultaneous behavior of LFP, CSD, and MUA; population-resolved cellular contributions; known recording geometry; and experimental laminar or multidimensional organization. Increasing the number and independence of such observables should progressively restrict the space of plausible mechanisms, which is one of the central motivations for large-scale mechanistic modeling^47^.

### Limitations and scope

Several limitations define the scope of these conclusions. First, the Peak, Mid, Low, and Off-best conditions are operational cortical-column response levels rather than explicit simulations of individual acoustic frequencies, empirically fitted tuning curves, or anatomically registered A1 locations. Each condition combined one fixed connectivity realization with one thalamic input strength; therefore, between-condition differences reflect the combined column/drive states and do not independently estimate thalamic-drive effects across connectivity realizations.

Second, the multi-column configurations used for the device analysis were constructed by spatially arranging independently simulated cortical columns. This design was intentional because it allowed source organization to be manipulated while preserving the full biophysical dynamics of each constituent column, but it does not reproduce horizontal synaptic interactions, shared inputs, or continuous tonotopic connectivity among neighboring cortical columns. Extending the model to an interconnected multi-column network would substantially increase computational cost and, importantly, would require additional experimental constraints to validate large-scale intercolumnar connectivity and dynamics that were not available when the model was constructed. The **Fig. 4** results should therefore be interpreted as an isolation of measurement geometry under controlled source configurations rather than as a complete simulation of a spatially extended auditory cortical sheet. These source configurations are also limited in terms of available permutations, as 25 cortical column variations were used in unique arrangements for each technical replicate. In addition, device lead field voxelization leads to imperfect cortical column placement and summing of cell membrane currents, acting essentially as a spatial filter. Ideal implementation requires an approach such as custom mesh generation to perfectly isolate all membrane currents but also requires substantially greater resources and processing. These issues will be addressed in upcoming studies.

Third, the experimental sample is limited to six DiSc and eight UPROBE penetrations. The noise– mean-tone correspondence was consistent across the available recordings for both geometries, but its generality across animals, cortical fields, stimulus sets, and behavioral states remains to be established; for example, selective attention can affect both frequency tuning and side-band inhibition^45^. Moreover, the correlation analysis tested for the presence of shared spatial organization between broadband-noise and tone-evoked responses, not for their statistical equivalence or the absence of stimulus-dependent differences. The observed positive correlations therefore should not be interpreted as evidence that broadband noise and tones produce identical spatial response organization. Because broadband noise distributes energy across much of the frequency range represented by the tone set, substantial correspondence with the mean tone response may be expected, whereas frequency-specific inhibition and nonlinear spectral interactions could produce differences not captured by the present analysis. Averaging across tones may further emphasize spatial components shared across frequencies while attenuating frequency-specific structure. The DiSc dataset is particularly limited for evaluating such variability in tone-response organization, as only two penetrations exhibited a clearly separated highest-response tone. This separation describes the tested response magnitudes rather than establishing statistically significant frequency selectivity. Larger datasets and analyses explicitly designed to test noise–tone differences or equivalence will be required to determine which components of these geometry-appropriate organizational relationships generalize across stimulus conditions. The experimental measurements have additional limitations. Repeated penetrations may be dependent, and penetration-level statistical summaries do not establish replication across independent animals. Absolute cortical depths and angular registration were unavailable, preventing assignment of the response maps to cortical tonotopy. The devices used different preprocessing and amplitude units, precluding absolute cross-device amplitude comparisons. Finally, the repeated fixed playback sequence confounded frequency with sequence position and preceding stimulus history, and trigger alignment did not establish acoustic arrival time.

Fourth, the experimental comparison was intentionally performed at the level of response organization rather than direct simulation-to-penetration matching. No attempt was made to fit model parameters separately to individual experimental recordings, and anatomical registration was insufficient to assert that a given simulated source configuration corresponded directly to a particular biological penetration. In addition, the recorded and simulated DiSc differ in contact count and span in order to provide a better simulated comparison of devices, while still performing as strongly as possible in implantation. Consequently, the present results support the preservation of geometry-appropriate organizational features, not one-to-one prediction of experimental waveforms, amplitudes, or cellular states.

Finally, the conclusions inherit the biological assumptions and approximations of the underlying macaque auditory thalamocortical model^18^. As additional anatomical, physiological, and cell-type-specific data become available, future model iterations may alter the quantitative contribution of individual populations while leaving some higher-order organizational relationships intact. Testing which conclusions survive such model refinement will itself provide a useful measure of their robustness.

The framework could be extended to multishank, mesh or other distributed arrays through geometry-specific forward operators validated for their contacts and conductivity assumptions. Modeling traveling waves would additionally require interacting cortical regions with justified horizontal connections, delays and common inputs. The present independent-column arrangements do not simulate intercolumnar propagation or estimate effective connectivity. Such applications remain prospective and require a separately validated dynamical and inference framework.

### Implications for interpreting extracellular electrophysiology

Neural-interface technologies are rapidly diversifying in channel count, contact density, spatial extent, and dimensionality^10,23,24,26,56^. These developments provide access to increasingly rich neural datasets, but they also make it less appropriate to treat recordings from different devices as interchangeable observations of the same latent signal. High-density and multidimensional arrays do not merely record *more* activity; they sample different spatial combinations of the underlying extracellular field and can therefore expose different aspects of circuit organization.

The framework presented here provides one strategy for interpreting those differences. By separating the generation of extracellular activity from its subsequent sampling, a common biophysical model can serve as an intermediate reference between circuit mechanisms and multiple recording technologies. In this formulation, disagreement between two probes need not imply disagreement about the neural source. Instead, it may reveal that the probes preserve different spatial projections of that source. Conversely, agreement across distinct geometries becomes especially informative when a predicted organizational relationship survives both biological and measurement transformations.

More broadly, our results argue for evaluating extracellular recordings at multiple levels simultaneously. Precise waveforms remain important when the scientific question concerns timing, oscillatory phase, synaptic kinetics, or local current flow. Cellular contributions are essential when assigning mechanisms. Spatial organization becomes critical when comparing recording interfaces. Population-level relationships can provide robust validation targets when biological variability precludes literal replication of an individual recording. No single level should replace the others; their combination provides progressively stronger constraints on mechanistic interpretation.

Thus, the principal contribution of the present framework is not that a biophysical model reproduces one particular experimental waveform or that one recording geometry is superior to another. Rather, the model makes it possible to follow the same underlying cortical activity through successive transformations, from circuit recruitment to signal-specific cellular generation, to geometry-dependent measurement, and to ask which features remain stable at each stage. This perspective provides a foundation for comparing emerging neural interfaces within a shared mechanistic framework and for using increasingly detailed circuit models not simply to imitate electrophysiological recordings, but to explain why different recordings of the same neural system can look different while still preserving common organization.

## Conclusion

Biophysically detailed forward modeling provides a means of separating the biological generation of extracellular activity from its geometry-dependent measurement. By following an imposed set of cortical-column response conditions through extracellular signal generation and virtual electrode sampling, we showed that organizational relationships can be expressed across LFP, CSD, and MUA while different recording geometries represent different spatial dimensions of common underlying cortical sources. Experimental macaque auditory-cortex recordings provided evidence for the corresponding geometry-appropriate organization across heterogeneous penetrations, supporting the use of organizational relationships, rather than exact waveform correspondence, as validation targets for multiscale models.

These findings emphasize that an extracellular recording represents neither the biological source nor the recording device in isolation, but their interaction. Recording geometry therefore determines not simply how strongly neural activity is measured, but which dimensions of circuit organization are accessible to measurement. Integrating biophysical circuit models with forward models of neural interfaces offers a framework for interpreting measurements across recording technologies and, prospectively, for selecting or designing electrode configurations according to the biological organization and scientific questions they are intended to resolve.

## Acknowledgements

Research supported by NIH RF1NS133972, UG3NS125487, R01NS133972, R01DC012947, R01DC019979, R01MH134118-01, NIH R01NS128924-01, P50MH109429, ARL Cooperative Agreement W911NF-22-2-0139, and NYS DOH SCIRB C38328GG. This research is dedicated to the memory of William W. Lytton (8/6/2026). We thank Nikita Novikov, Ethan Irby, and Scott McElroy for assistance in development of the biophysical models; Michael Hines, Ted Carnevale, and Robert McDougal for NEURON simulator and ModelDB support.

## Notes

### Competing Interest Statement

The authors have declared no competing interest.

### Summary of Updates

Expanded analysis of neuronal population contributions. We extended the model analysis across all four cortical network and input conditions, examining how different neuronal populations contribute to local field potentials, current source density, and multiunit activity. We added quantitative comparisons, statistical analyses, and supplementary results to clarify how these contributions depend on network state and signal modality. Additional analyses of electrode geometry. We expanded the simulations to examine the effects of cortical column rotations, small tilts, and electrode orientation. These analyses further characterize the spatial sensitivity of directional DiSc arrays compared with conventional laminar probes. We also clarified the electrode specifications, recording configurations, and modeling assumptions. Strengthened experimental analyses. We revised the analyses of macaque auditory cortex recordings from six DiSc and eight conventional laminar probe penetrations. The updated results include more complete frequency-response maps, spatial correlations, quantitative response comparisons, and uncertainty estimates. We also clarified the criteria used to characterize frequency-dependent responses and the limitations of their interpretation. Improved presentation and reproducibility. We revised the abstract, Methods, Results, and Discussion; improved figure readability, labeling, and legends; and expanded the supplementary figures and numerical data.

## References

1. Buzsáki, G., Anastassiou, C. A. & Koch, C. The origin of extracellular fields and currents: EEG, ECoG, LFP and spikes. Nat. Rev. Neurosci. https://www.nature.com/articles/nrn3241 (2012).

2. Einevoll, G. T., Kayser, C., Logothetis, N. K. & Panzeri, S. Modelling and analysis of local field potentials for studying the function of cortical circuits. Nat. Rev. Neurosci. 14, 770–785 (2013).

3. Nicholson, C. & Freeman, J. A. Theory of current source-density analysis and determination of conductivity tensor for anuran cerebellum. J. Neurophysiol. 38, 356–368 (1975).

4. Kajikawa, Y., Smiley, J. F. & Schroeder, C. E. Primary Generators of Visually Evoked Field Potentials Recorded in the Macaque Auditory Cortex. J. Neurosci. 37, 10139–10153 (2017).

5. Lindén, H. et al. Modeling the spatial reach of the LFP. Neuron 72, 859–872 (2011).

6. Kajikawa, Y. & Schroeder, C. E. How local is the local field potential? Neuron 72, 847–858 (2011).

7. Moffitt, M. A. & McIntyre, C. C. Model-based analysis of cortical recording with silicon microelectrodes. Clin. Neurophysiol. 116, 2240–2250 (2005).

8. Nelson, M. J. & Pouget, P. Physical model of coherent potentials measured with different electrode recording site sizes. J. Neurophysiol. 107, 1291–1300 (2012).

9. Ye, Z. et al. Ultra-high-density Neuropixels probes improve detection and identification in neuronal recordings. Neuron 113, 3966–3982.e12 (2025).

10. Scholvin, J. et al. Close-packed silicon microelectrodes for scalable spatially oversampled neural recording. IEEE Trans. Biomed. Eng. 63, 120–130 (2016).

11. Meszéna, D. et al. Optimal inter-electrode distances for maximizing single unit yield per electrode in neural recordings. Microsyst. Nanoeng. 12, 41 (2026).

12. Willis, J. A. et al. Optimizing electrode placement and information capacity for local field potentials in cortex. bioRxivorg 2025.04. 25.650658 (2025) doi:10.1101/2025.04.25.650658.

13. Grill, W. M. et al. A roadmap to navigate the future of neural engineering. J. Neural Eng. 23, 041501 (2026).

14. Tomsett, R. J. et al. Virtual Electrode Recording Tool for EXtracellular potentials (VERTEX): comparing multi-electrode recordings from simulated and biological mammalian cortical tissue. Brain Struct. Funct. 220, 2333–2353 (2015).

15. Hagen, E. et al. Hybrid scheme for modeling local field potentials from point-neuron networks. Cereb. Cortex 26, 4461–4496 (2016).

16. Hagen, E., Næss, S., Ness, T. V. & Einevoll, G. T. Multimodal Modeling of Neural Network Activity: Computing LFP, ECoG, EEG, and MEG Signals With LFPy 2.0. Front. Neuroinform. 12, 92 (2018).

17. Anastassiou, C. A., Perin, R., Hill, S. L., Markram, H. & Koch, C. A biophysically detailed model of neocortical local field potentials predicts the critical role of active membrane currents. Neuron https://www.cell.com/neuron/fulltext/S0896-6273(13)00443-1 (2013).

18. Dura-Bernal, S. et al. Data-driven multiscale model of macaque auditory thalamocortical circuits reproduces in vivo dynamics. Cell Rep. 42, 113378 (2023).

19. Prinz, A., Bucher, D. & Marder, E. Similar network activity from disparate circuit parameters. Nat. Neurosci. 7, 1345–1352 (2004).

20. Marder, E. & Taylor, A. L. Multiple models to capture the variability in biological neurons and networks. Nat. Neurosci. 14, 133–138 (2011).

21. Gutzen, R. et al. Reproducible neural network simulations: Statistical methods for model validation on the level of network activity data. Front. Neuroinform. 12, 90 (2018).

22. Chien, V. S. C., Wang, P., Maess, B., Fishman, Y. & Knösche, T. R. Laminar neural dynamics of auditory evoked responses: Computational modeling of local field potentials in auditory cortex of non-human primates. Neuroimage 281, 120364 (2023).

23. Rios, G., Lubenov, E. V., Chi, D., Roukes, M. L. & Siapas, A. G. Nanofabricated neural probes for dense 3-D recordings of brain activity. Nano Lett. 16, 6857–6862 (2016).

24. Jun, J. J. et al. Fully integrated silicon probes for high-density recording of neural activity. Nature 551, 232–236 (2017).

25. Hong, G. & Lieber, C. M. Novel electrode technologies for neural recordings. Nat. Rev. Neurosci. 20, 330–345 (2019).

26. Abrego, A. et al. Sensing local field potentials with a directional and scalable depth electrode array. J. Neural Eng. 20, 016041 (2023).

27. Shores, R. et al. Microelectrode arrays enable directional stereo-EEG during kainate-mediated seizures. bioRxivorg (2026) doi:10.64898/2026.06.11.731746.

28. Medani, T. et al. High-resolution directional depth electrodes: Open-source FEM lead-field modeling, characterization, and validation. arXiv [physics.med-ph] (2025).

29. Dura-Bernal, S. et al. NetPyNE, a tool for data-driven multiscale modeling of brain circuits. Elife 8, (2019).

30. Carnevale, N. T. & Hines, M. L. The NEURON Book. (Cambridge University Press, 2006).

31. Koch, C. & Segev, I. Methods in Neuronal Modeling: From Ions to Networks. (MIT Press, London, England, 1998).

32. Lakatos, P. et al. Global dynamics of selective attention and its lapses in primary auditory cortex. Nat. Neurosci. 10.1038/nn.4386 (2016) doi:10.1038/nn.4386.

33. Henin, S. et al. Learning hierarchical sequence representations across human cortex and hippocampus. Sci. Adv. 7, eabc4530 (2021).

34. Pedregosa, F., Varoquaux, G. & Gramfort, A. Scikit-learn: Machine learning in Python. the Journal of machine https://www.jmlr.org/papers/volume12/pedregosa11a/pedregosa11a.pdf?ref=https:/ (2011).

35. Hines, M. L., Morse, T., Migliore, M., Carnevale, N. T. & Shepherd, G. M. ModelDB: A Database to Support Computational Neuroscience. J. Comput. Neurosci. 17, 7–11 (2004).

36. Reimann, M. W. et al. A biophysically detailed model of neocortical local field potentials predicts the critical role of active membrane currents. Neuron 79, 375–390 (2013).

37. Neymotin, S. A. et al. Human Neocortical Neurosolver (HNN), a new software tool for interpreting the cellular and network origin of human MEG/EEG data. Elife 9, 740597 (2020).

38. Kohl, C., Parviainen, T. & Jones, S. R. Neural mechanisms underlying human auditory evoked responses revealed by Human Neocortical Neurosolver. Brain Topogr. 35, 19–35 (2022).

39. Kajikawa, Y., Mackey, C. A. & O’Connell, M. N. Laminar pattern of sensory-evoked dynamic high-frequency oscillatory activity in the macaque auditory cortex. Cereb. Cortex 34, bhae338 (2024).

40. Leszczyński, M. et al. Dissociation of broadband high-frequency activity and neuronal firing in the neocortex. Science Advances 6, eabb0977 (2020).

41. Gerbella, M. et al. Histological assessment of a chronically implanted cylindrically-shaped, polymer-based neural probe in the monkey. J. Neural Eng. 18, 024001 (2021).

42. Shobe, J. L., Claar, L. D., Parhami, S., Bakhurin, K. I. & Masmanidis, S. C. Brain activity mapping at multiple scales with silicon microprobes containing 1,024 electrodes. J. Neurophysiol. 114, 2043–2052 (2015).

43. Morel, A., Garraghty, P. E. & Kaas, J. H. Tonotopic organization, architectonic fields, and connections of auditory cortex in macaque monkeys. J. Comp. Neurol. 335, 437–459 (1993).

44. Kaas, J. H. & Hackett, T. A. Subdivisions of auditory cortex and processing streams in primates. Proc. Natl. Acad. Sci. U. S. A. 97, 11793–11799 (2000).

45. O’Connell, M. N., Barczak, A., Schroeder, C. E. & Lakatos, P. Layer specific sharpening of frequency tuning by selective attention in primary auditory cortex. J. Neurosci. 34, 16496–16508 (2014).

46. Schroeder, C. E. et al. Somatosensory input to auditory association cortex in the macaque monkey. J. Neurophysiol. 85, 1322–1327 (2001).

47. Dura-Bernal, S. et al. Large-scale mechanistic models of brain circuits with biophysically and morphologically detailed neurons. 44, (2024).

48. Kajikawa, Y., de La Mothe, L., Blumell, S. & Hackett, T. A. A comparison of neuron response properties in areas A1 and CM of the marmoset monkey auditory cortex: tones and broadband noise. J. Neurophysiol. 93, 22–34 (2005).

49. Kajikawa, Y. et al. Auditory cortical tuning to band-pass noise in primate A1 and CM: a comparison to pure tones. Neurosci. Res. 70, 401–407 (2011).

50. Marder, E. & Goaillard, J.-M. Variability, compensation and homeostasis in neuron and network function. Nat. Rev. Neurosci. 7, 563–574 (2006).

51. Goaillard, J.-M., Taylor, A. L., Schulz, D. J. & Marder, E. Functional consequences of animal-to-animal variation in circuit parameters. Nat. Neurosci. 12, 1424–1430 (2009).

52. McDougal, R. A., Bulanova, A. S. & Lytton, W. W. Reproducibility in computational neuroscience models and simulations. IEEE Trans. Biomed. Eng. 63, 2021–2035 (2016).

53. Neymotin, S. A. et al. Optimizing computer models of corticospinal neurons to replicate in vitro dynamics. J. Neurophysiol. 117, 148–162 (2017).

54. Neymotin, S. A., Dura-Bernal, S., Lakatos, P., Sanger, T. D. & Lytton, W. W. Multitarget Multiscale Simulation for Pharmacological Treatment of Dystonia in Motor Cortex. Front. Pharmacol. 7, 157 (2016).

55. Neymotin, S. A., Dura-Bernal, S., Moreno, H. & Lytton, W. W. Computer modeling for pharmacological treatments for dystonia. Drug Discov. Today Dis. Models 19, 51–57 (2016).

56. de la Mothe, L. A., Blumell, S., Kajikawa, Y. & Hackett, T. A. Cortical connections of the auditory cortex in marmoset monkeys: core and medial belt regions. J. Comp. Neurol. 496, 27–71 (2006).

