## Supplementary figures for "Biophysical modeling reveals how recording geometry shapes representations of cortical activity"

DiSc auditory cortical modeling · Figures S1–S9

Each figure legend immediately precedes its corresponding panels. The original vector-based figures are preserved, and the individual panels retain their published axes, scales, and labels.

#### Contents


**S1** Model and device illustration details (5 source pages)

**S2** Thalamic input population responses (3 panel pages)

**S3** Peak–Mid early MUA-like envelope contrast

**S4** Population spike-rate comparisons

**S5** Complete DiSc frequency maps and direction profiles (12 panel pages)

**S6** Complete UPROBE frequency-response profiles (8 panel pages)

**S7** UPROBE frequency-order comparison

**S8** Absolute population-response magnitudes (3 panel pages)

**S9** UPROBE contact orientation variation and region separability

### S1. Model and device illustration details

Enlarged components of Figure 1: model anatomy and simulation design, Peak/Mid and Low/Off-best response illustrations, model observables, and device/experimental framework.

(a) anatomical model and recruitment schematic

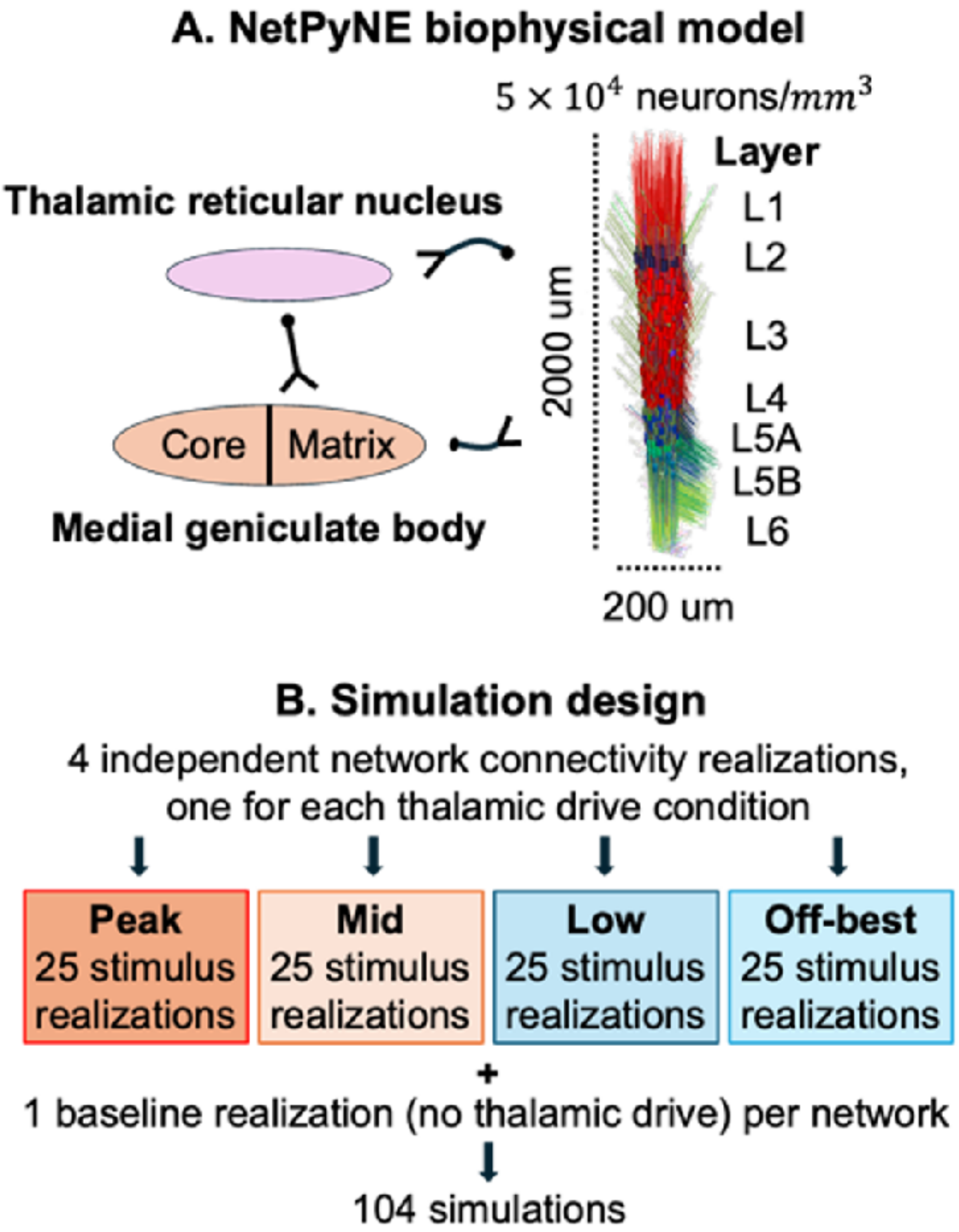

(b) example responses: Peak and Mid

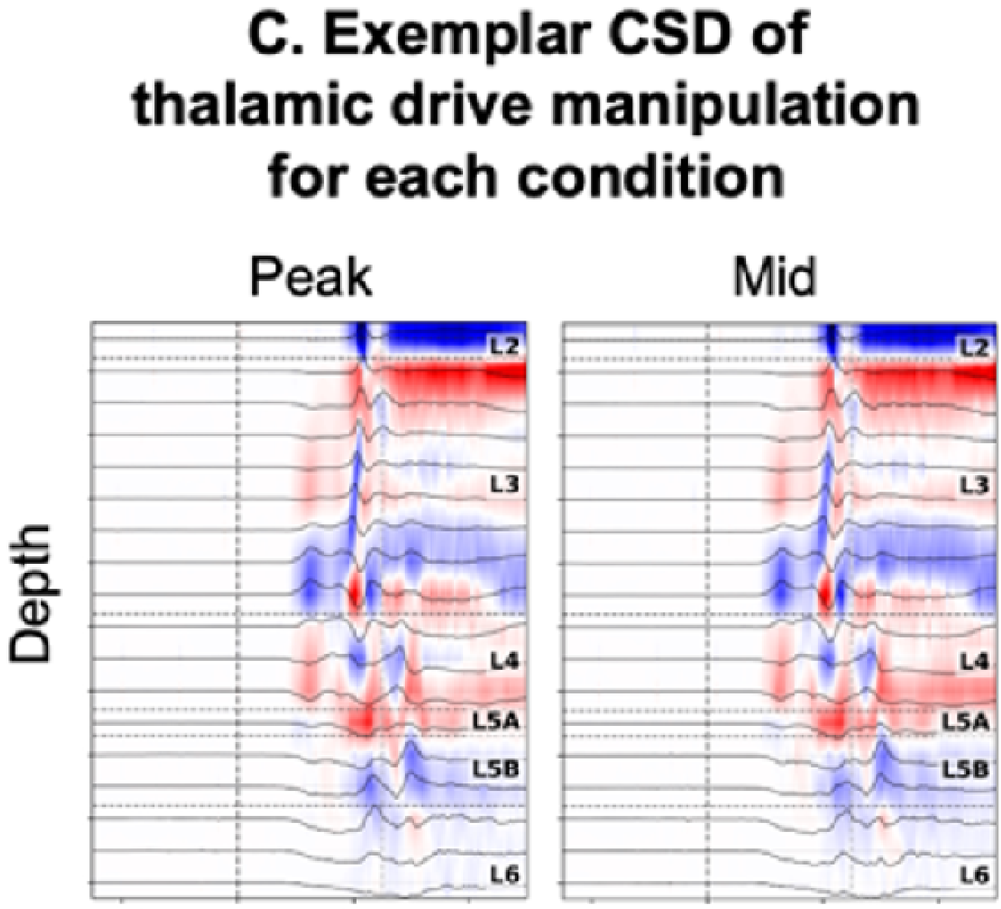

(b continued) example responses: Low and Off-best

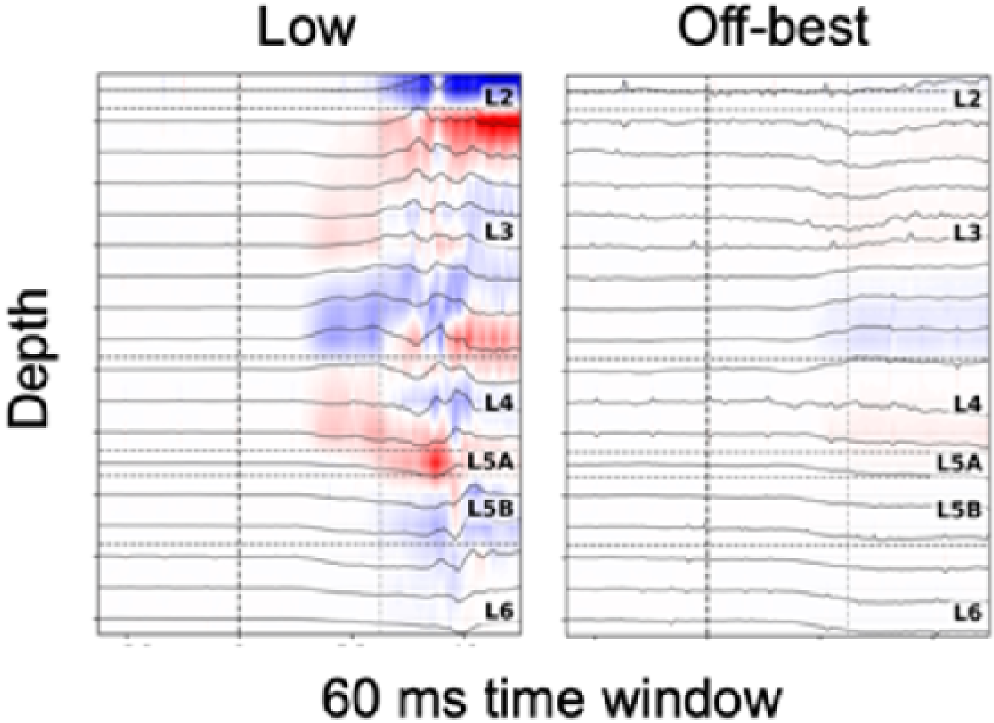

(c) model observables

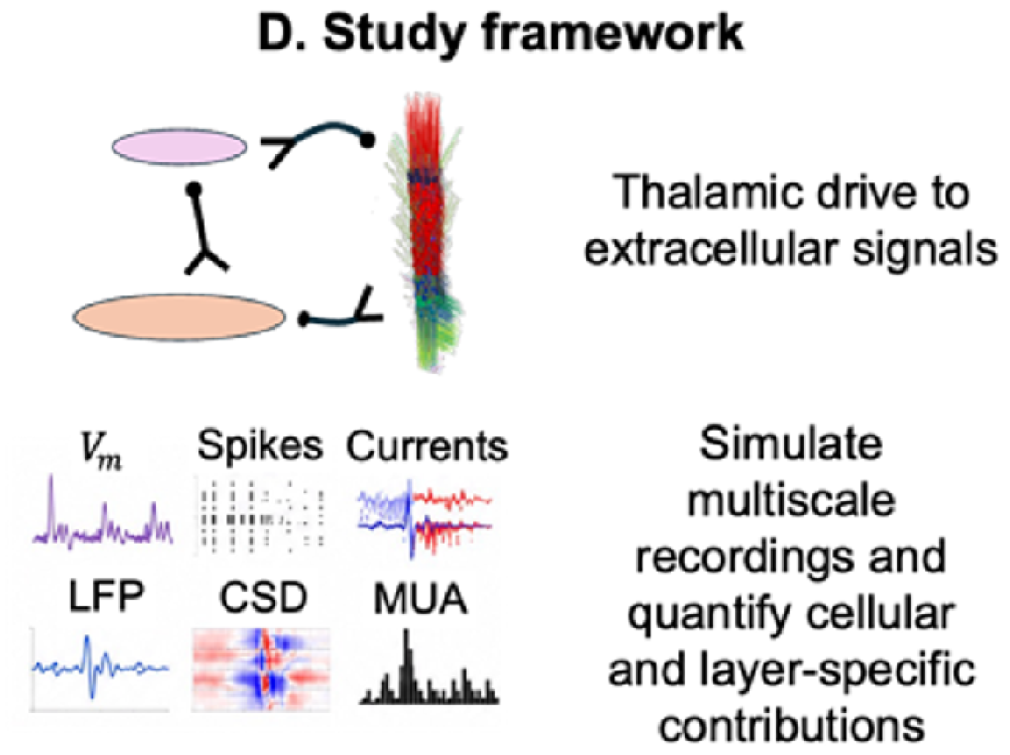

(d) electrode and experiment illustrations

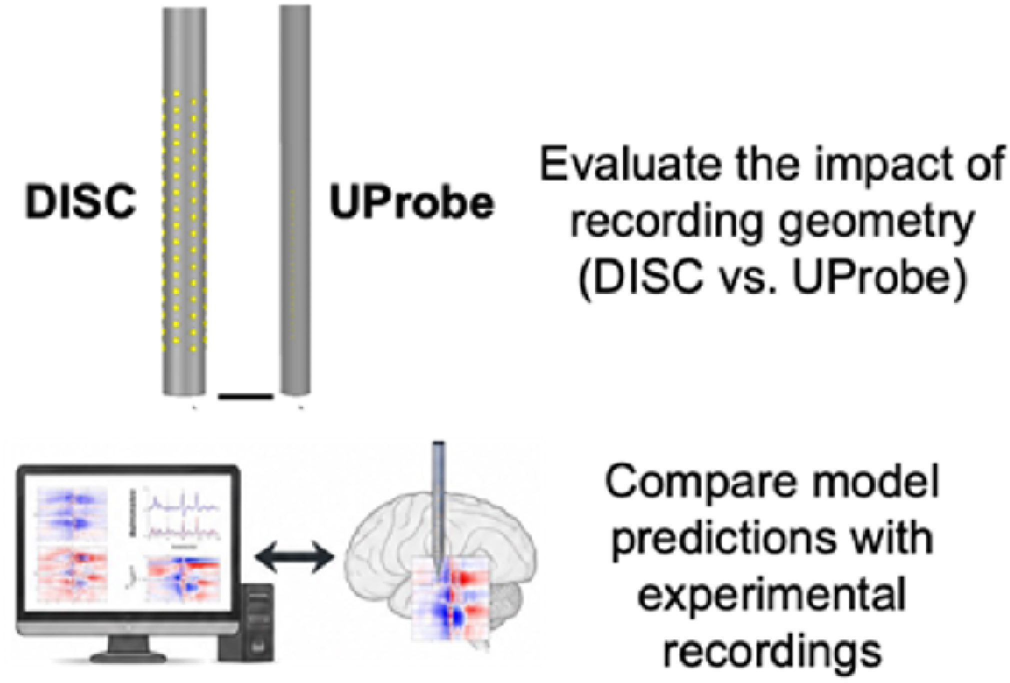

### S2. Thalamic input population responses

TC, HTC and TCM spike rasters for one example realization per condition (left) and population spike-rate profiles across 25 background-input realizations per condition (right). Rates use 1-ms bins and are normalized per neuron. Shading shows SEM. Time zero denotes thalamic input onset. State-wise axes share a scale within each population.

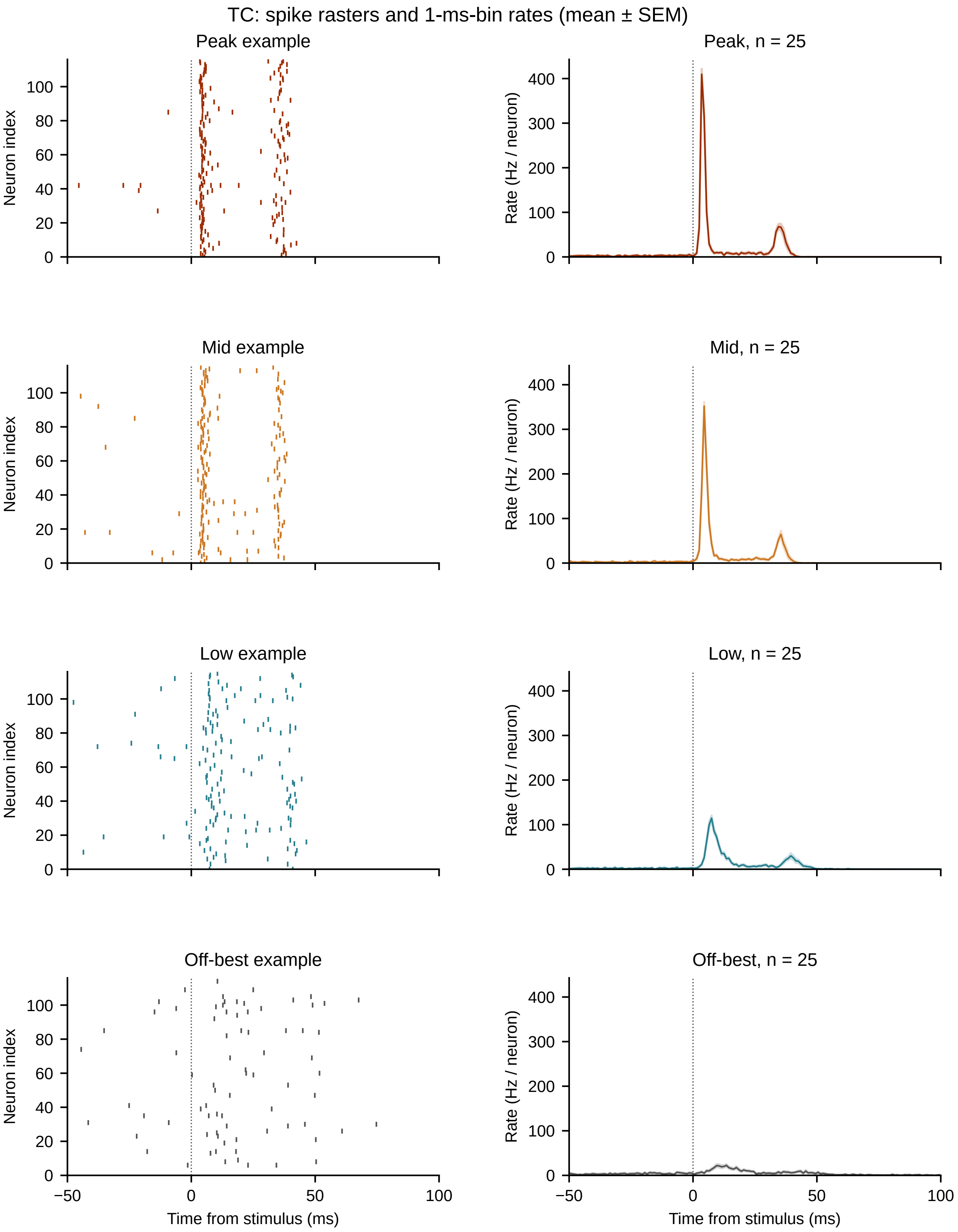
TC: spike rasters and 1-ms-bin rates (mean ± SEM)
Peak example
Peak, n = 25
Mid example
Mid, n = 25
Low example
Low, n = 25
Off-best example
Off-best, n = 25
Neuron index
Rate (Hz / neuron)
Time from stimulus (ms)
−50
0
50
100
20
40
60
80
200
300
400

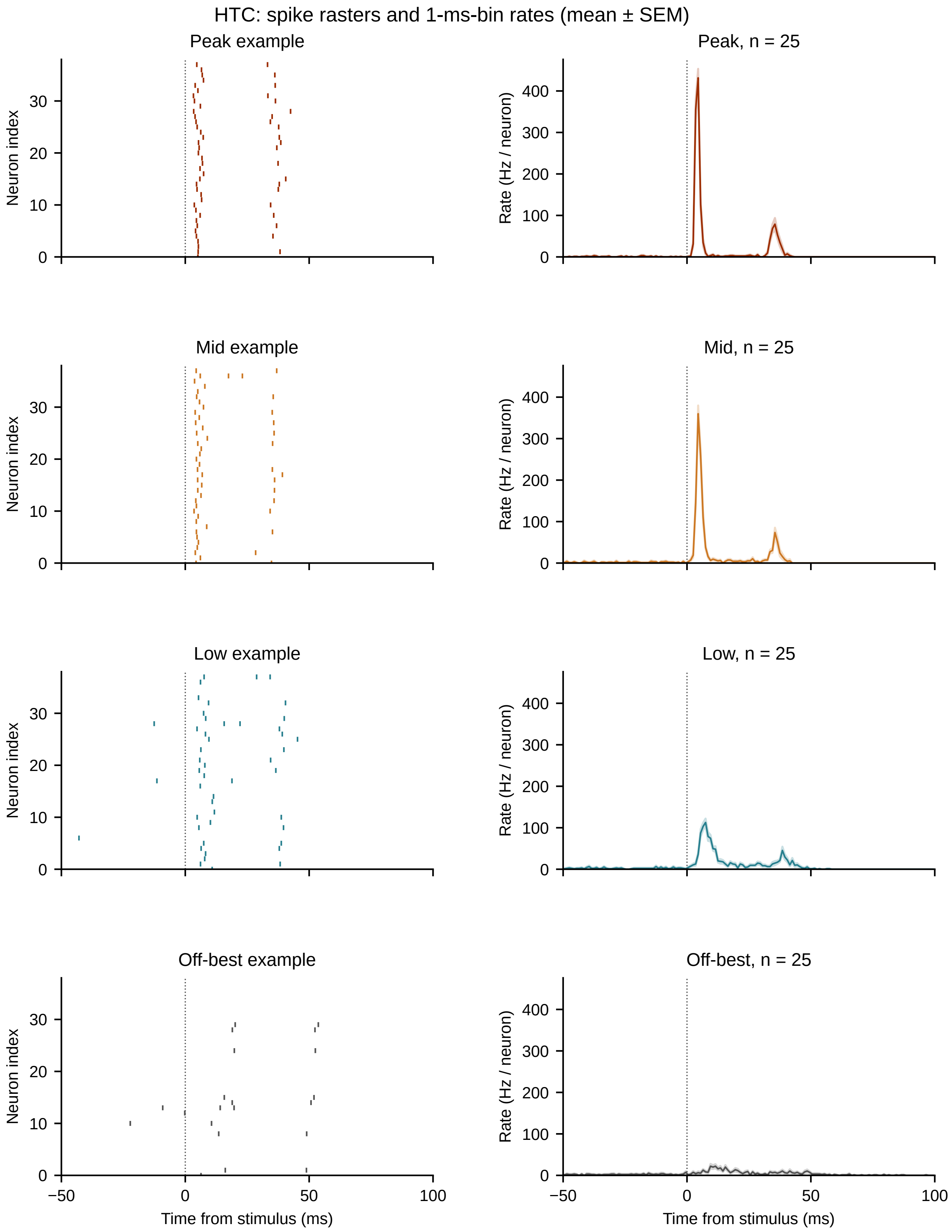
HTC: spike rasters and 1-ms-bin rates (mean ± SEM)
Peak example
Peak, n = 25
Mid example
Mid, n = 25
Low example
Low, n = 25
Off-best example
Off-best, n = 25
Neuron index
Rate (Hz / neuron)
Time from stimulus (ms)
0
10
20
30
100
200
300
400
−50
50

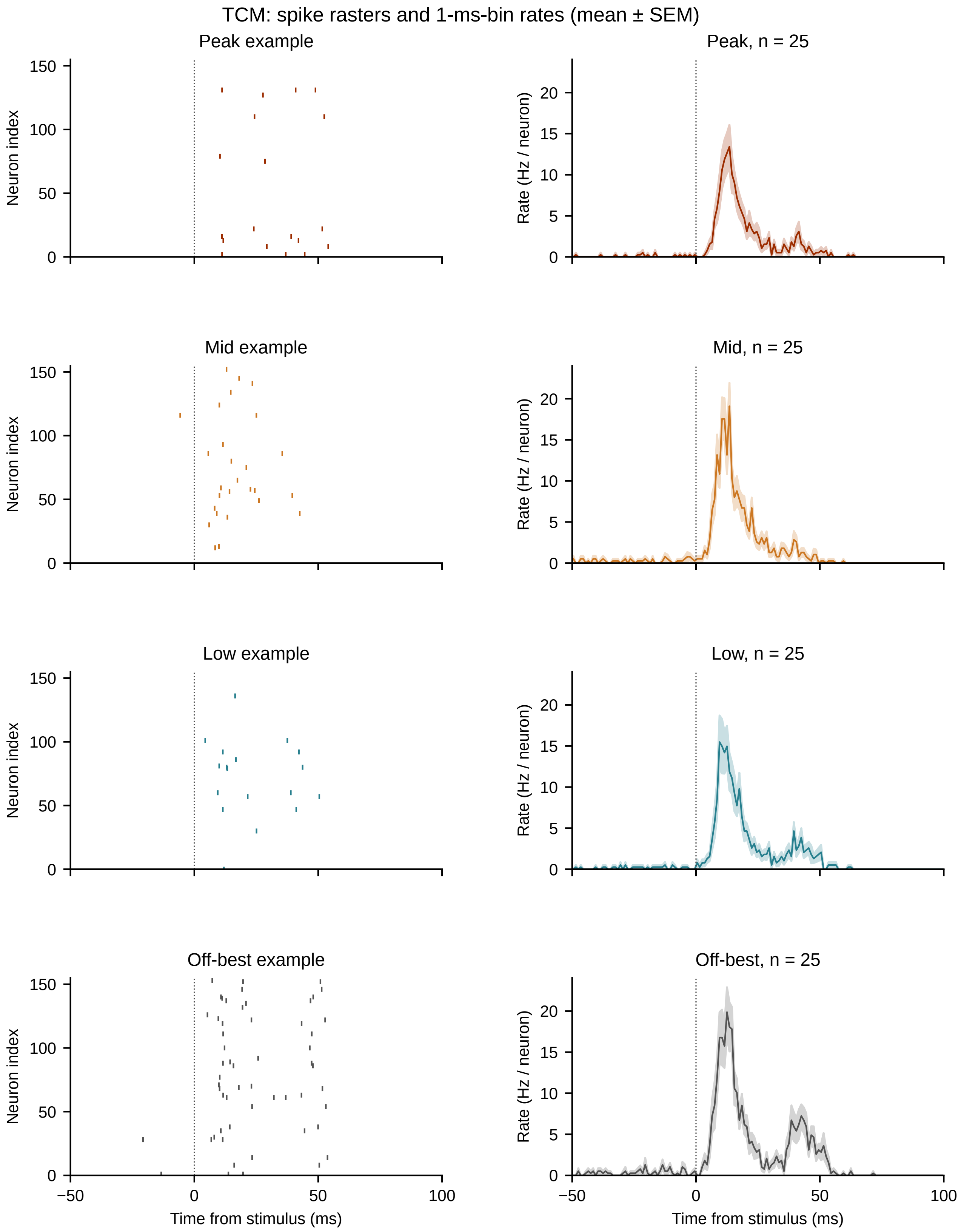
TCM: spike rasters and 1-ms-bin rates (mean ± SEM)
Peak example
Peak, n = 25
Mid example
Mid, n = 25
Low example
Low, n = 25
Off-best example
Off-best, n = 25
Neuron index
Rate (Hz / neuron)
0
50
100
150
5
10
15
20
−50
Time from stimulus (ms)

### S3. Peak–Mid early MUA-like envelope contrast

Points show 25 individual background-input realizations per state. Large markers and error bars show mean ± SEM. The right panel shows the mean difference and its 95% independent within-state bootstrap interval. Connectivity and drive differ together between states.

#### S3 Peak–Mid response contrast

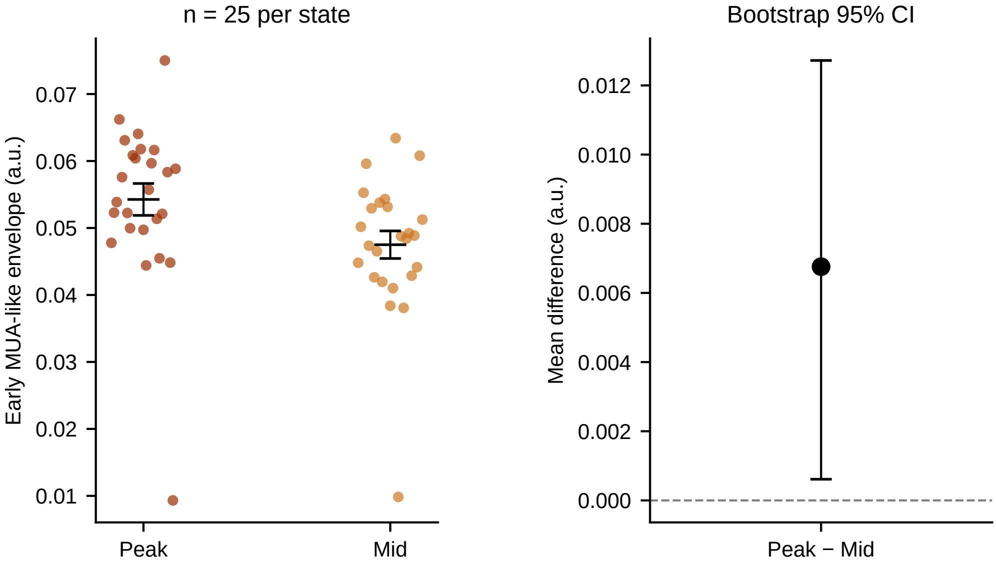


Welch p = 0.03695; Hedges g = 0.598.
Connectivity and drive differ between states.

### S4. Population spike-rate comparisons

Early-response spike rates and response-minus-baseline rates for ten cortical classes across four conditions. Symbols and error bars show mean ± SEM across 25 background-input realizations. Populations need not show a monotonic ordering even when extracellular response amplitudes broadly follow the imposed condition hierarchy.

**S4** Population spike rates

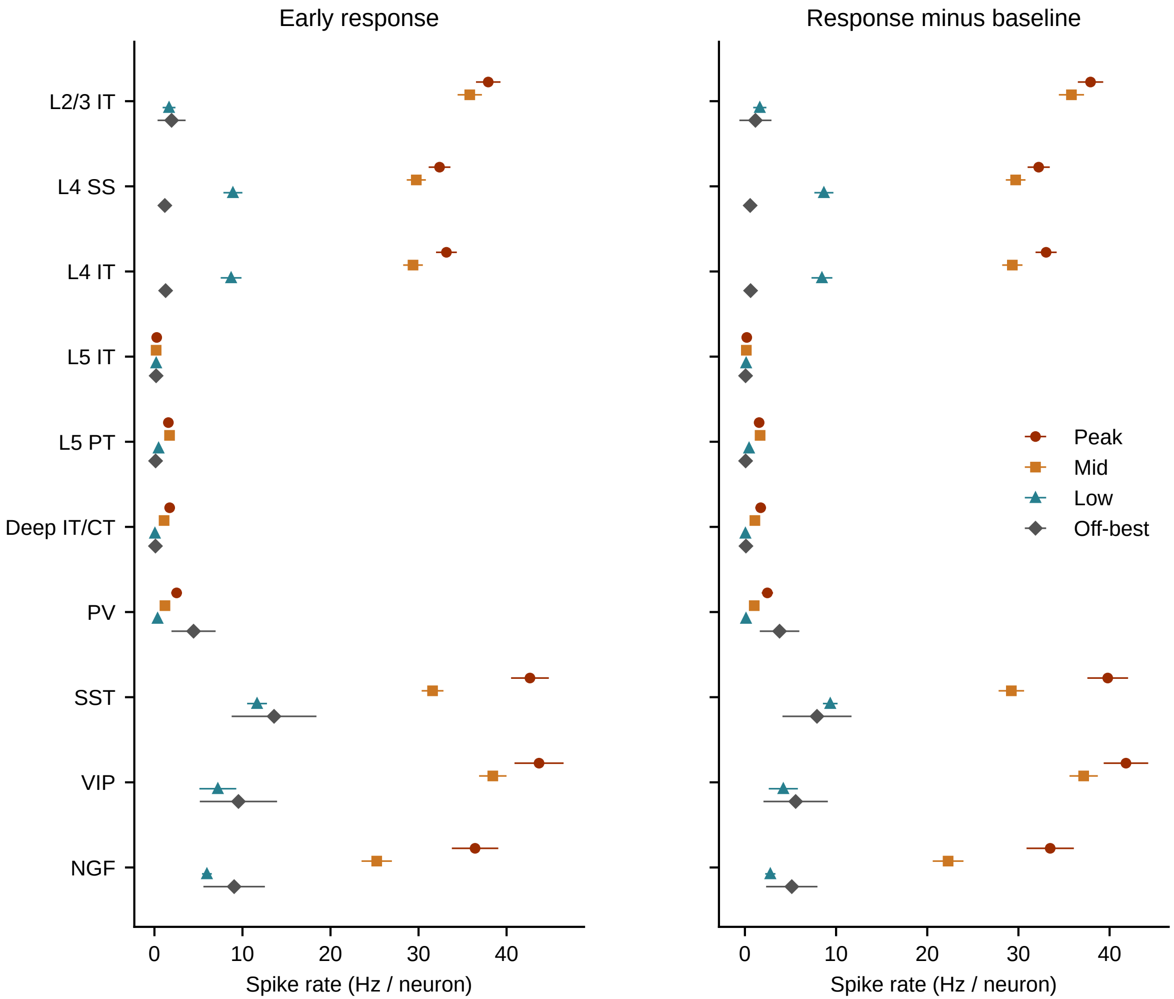

### S5. Complete DiSc frequency maps and direction profiles

Each penetration has fifteen maps (fourteen tones and BBN), sharing one color scale without per-tone normalization. Direction/depth indices range from 0 to 7. Companion profiles use one depth selected from the mean tone response for all frequencies; colors give trial-bootstrap support for the observed maximum-direction index. Magnitude is the mean of the strongest 16 contacts selected per tone, with trial SEM conditional on those contacts. All six penetrations are included.

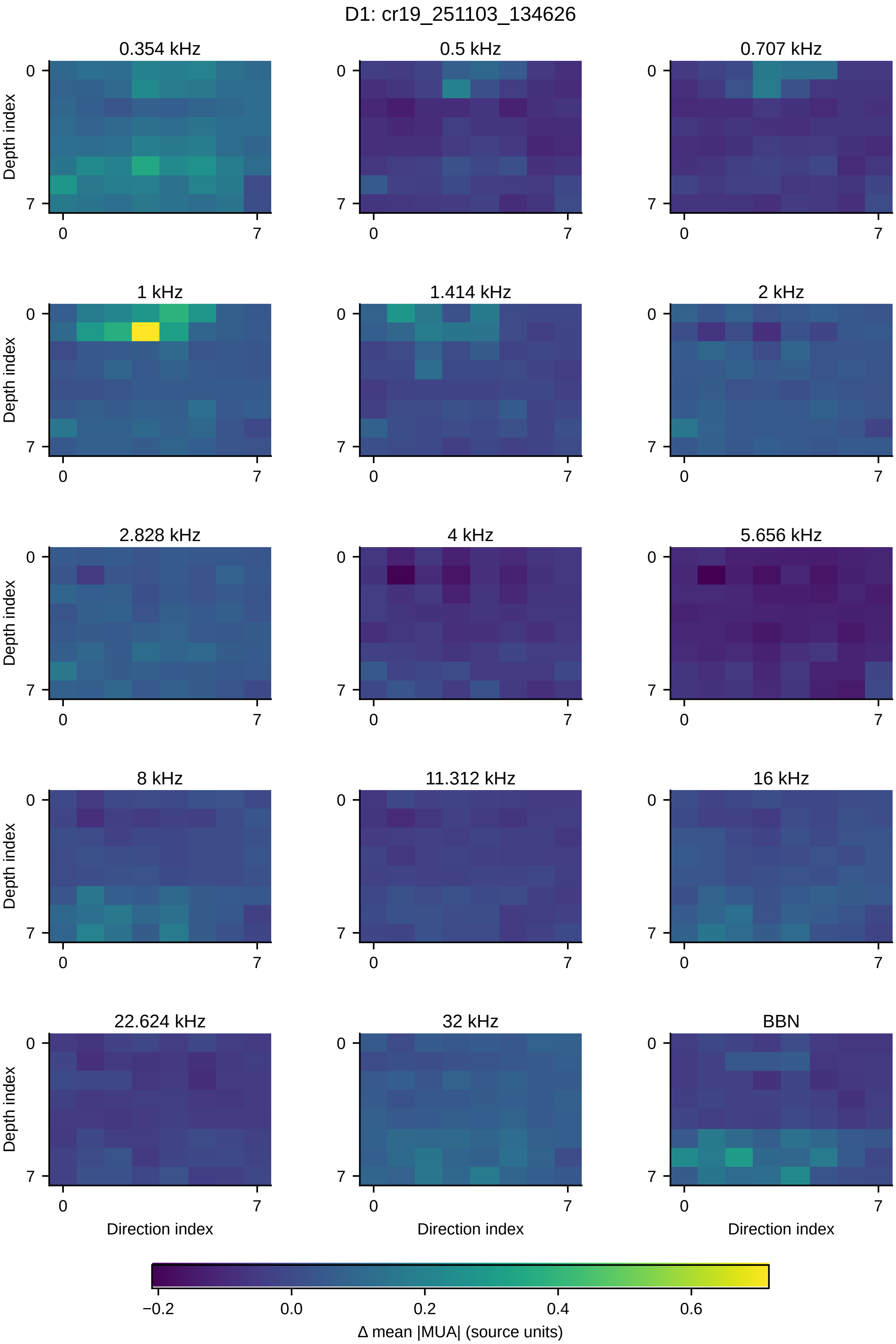
D1: cr19_251103_134626
0.354 kHz
0.5 kHz
0.707 kHz
1 kHz
1.414 kHz
2 kHz
2.828 kHz
4 kHz
5.656 kHz
8 kHz
11.312 kHz
16 kHz
22.624 kHz
32 kHz
BBN
Depth index
Direction index
0
7
−0.2
0.0
0.2
0.4
0.6
Δ mean |MUA| (source units)

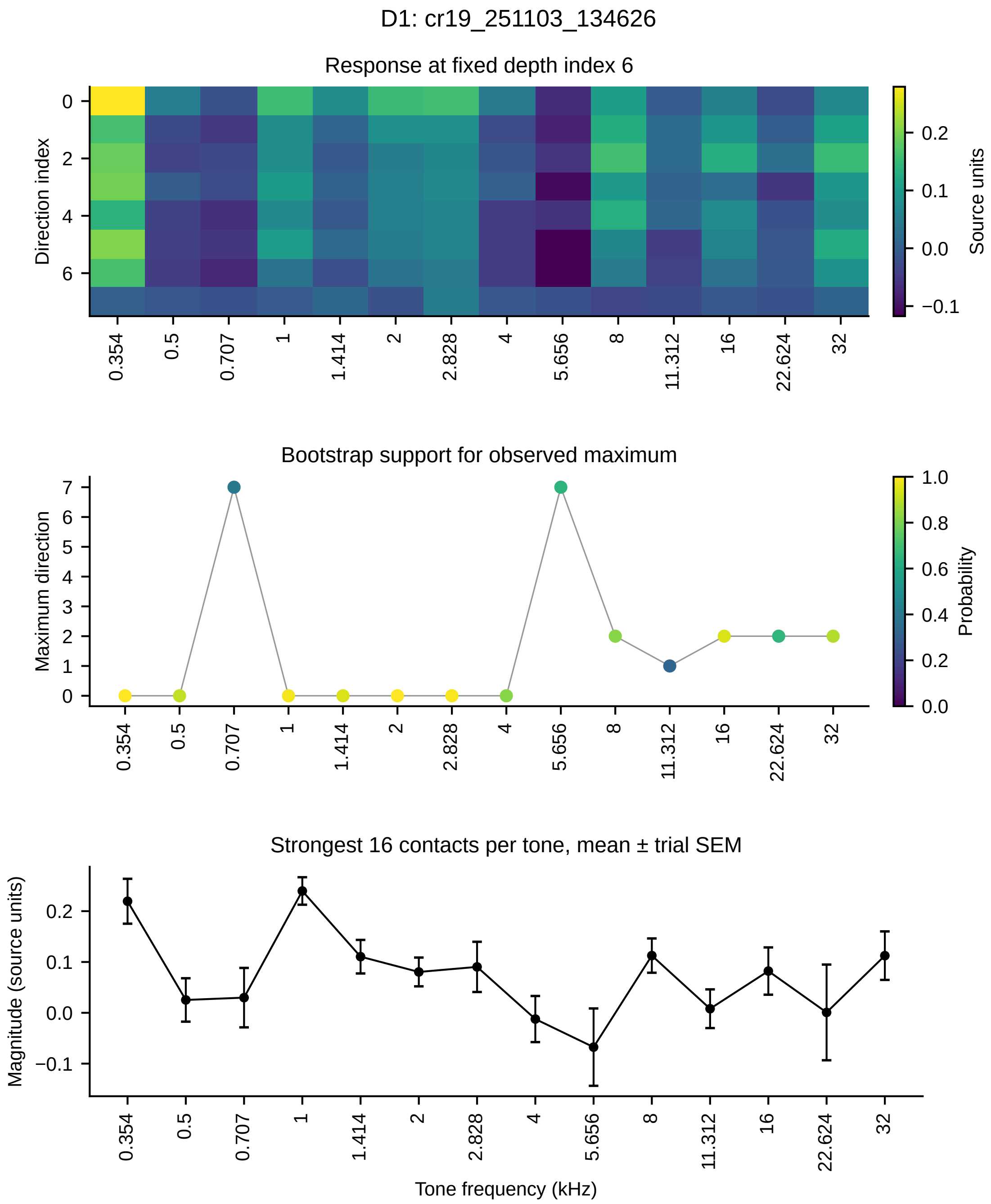
D1: cr19_251103_134626
Response at fixed depth index 6
Direction index
Source units
Bootstrap support for observed maximum
Maximum direction
Probability
Strongest 16 contacts per tone, mean ± trial SEM
Magnitude (source units)
Tone frequency (kHz)
0.354
0.5
0.707
1
1.414
2
2.828
4
5.656
8
11.312
16
22.624
32

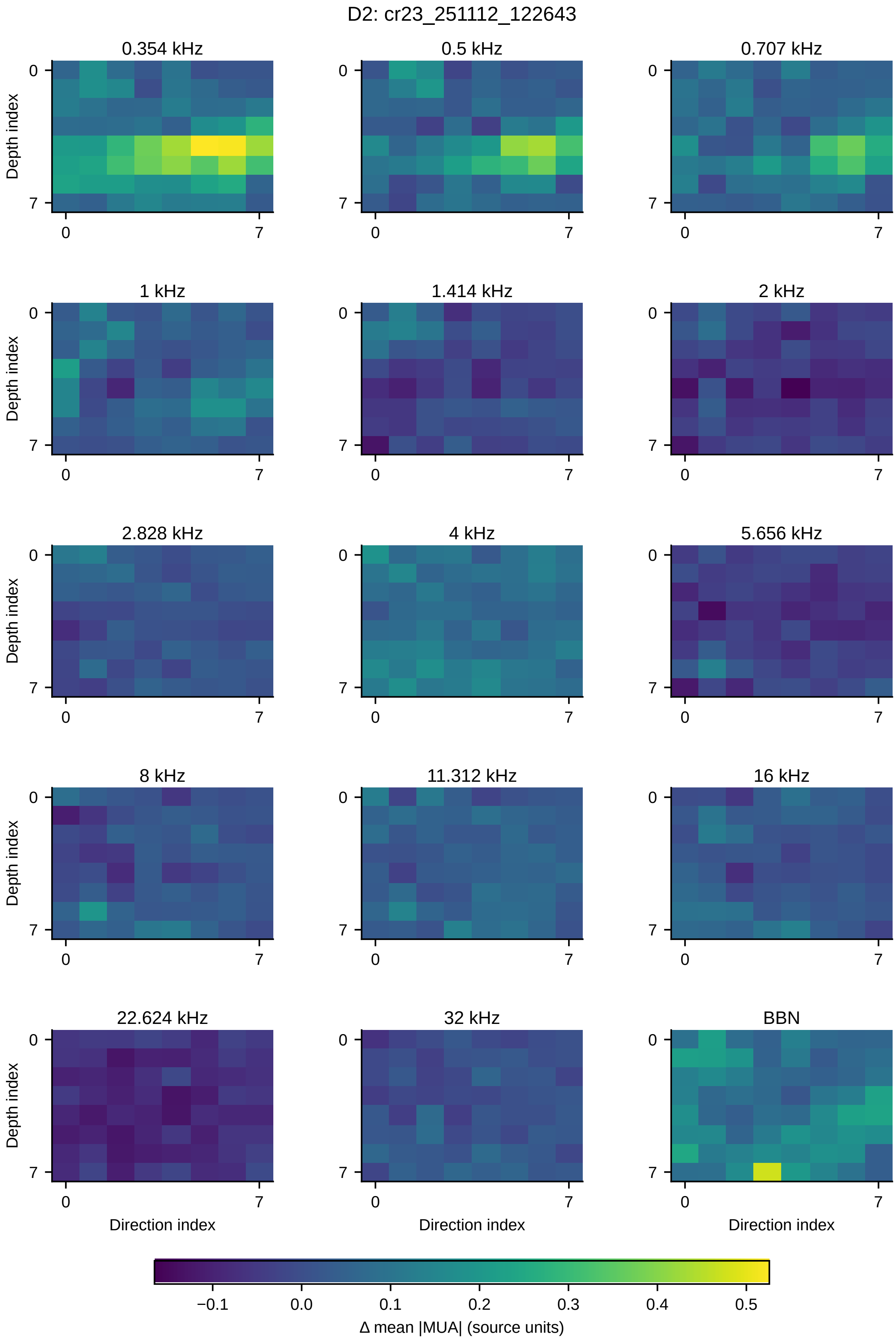
D2: cr23_251112_122643
0.354 kHz
0.5 kHz
0.707 kHz
1 kHz
1.414 kHz
2 kHz
2.828 kHz
4 kHz
5.656 kHz
8 kHz
11.312 kHz
16 kHz
22.624 kHz
32 kHz
BBN
Depth index
Direction index
0
7
−0.1
0.0
0.1
0.2
0.3
0.4
0.5
Δ mean |MUA| (source units)

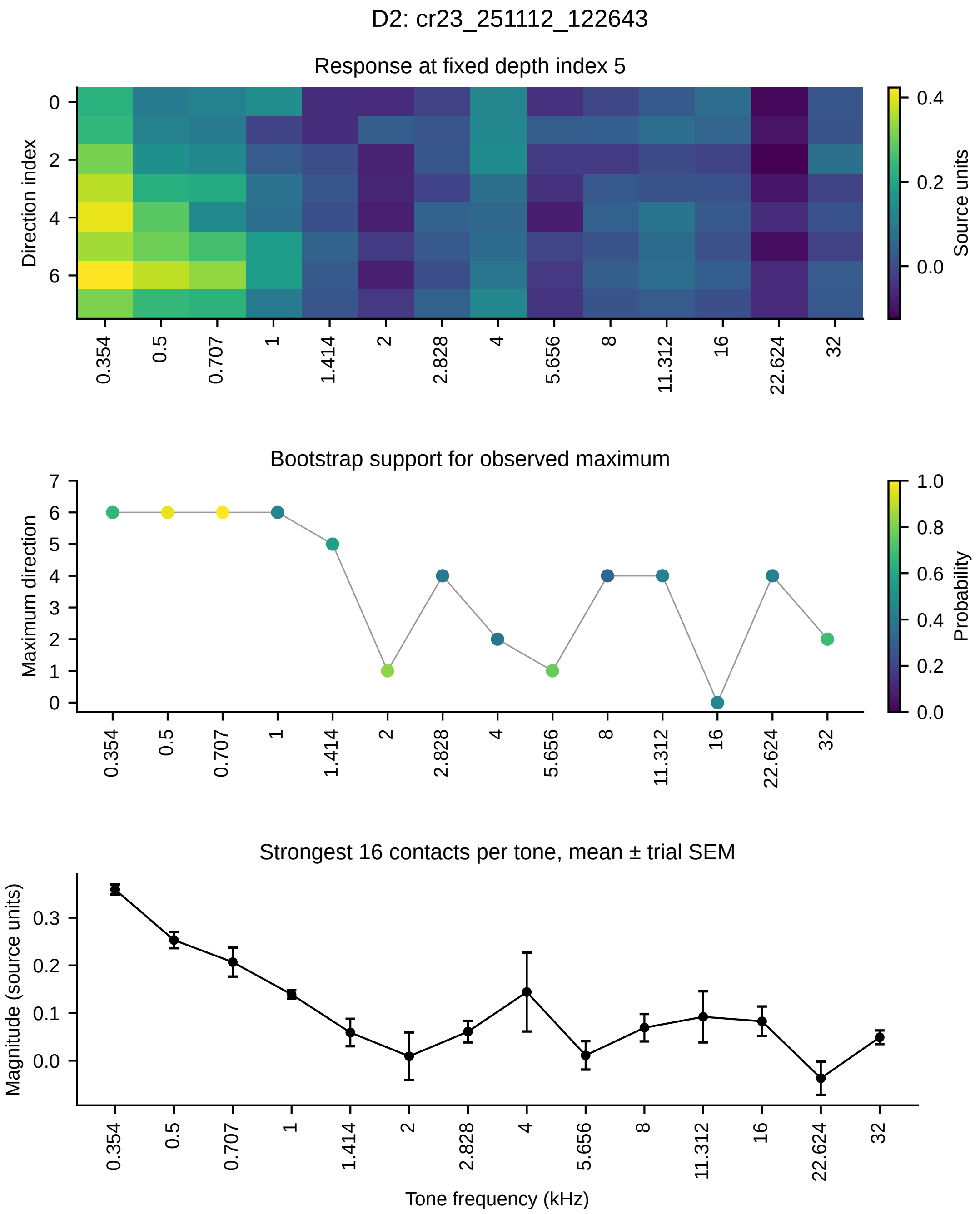
D2: cr23_251112_122643
Response at fixed depth index 5
Direction index
Source units
0
2
4
6
0.4
0.2
0.0
0.354
0.5
0.707
1
1.414
2
2.828
4
5.656
8
11.312
16
22.624
32
Bootstrap support for observed maximum
Maximum direction
Probability
0
1
2
3
4
5
6
7
1.0
0.8
0.6
0.4
0.2
0.0
Strongest 16 contacts per tone, mean ± trial SEM
Magnitude (source units)
0.3
0.2
0.1
0.0
Tone frequency (kHz)

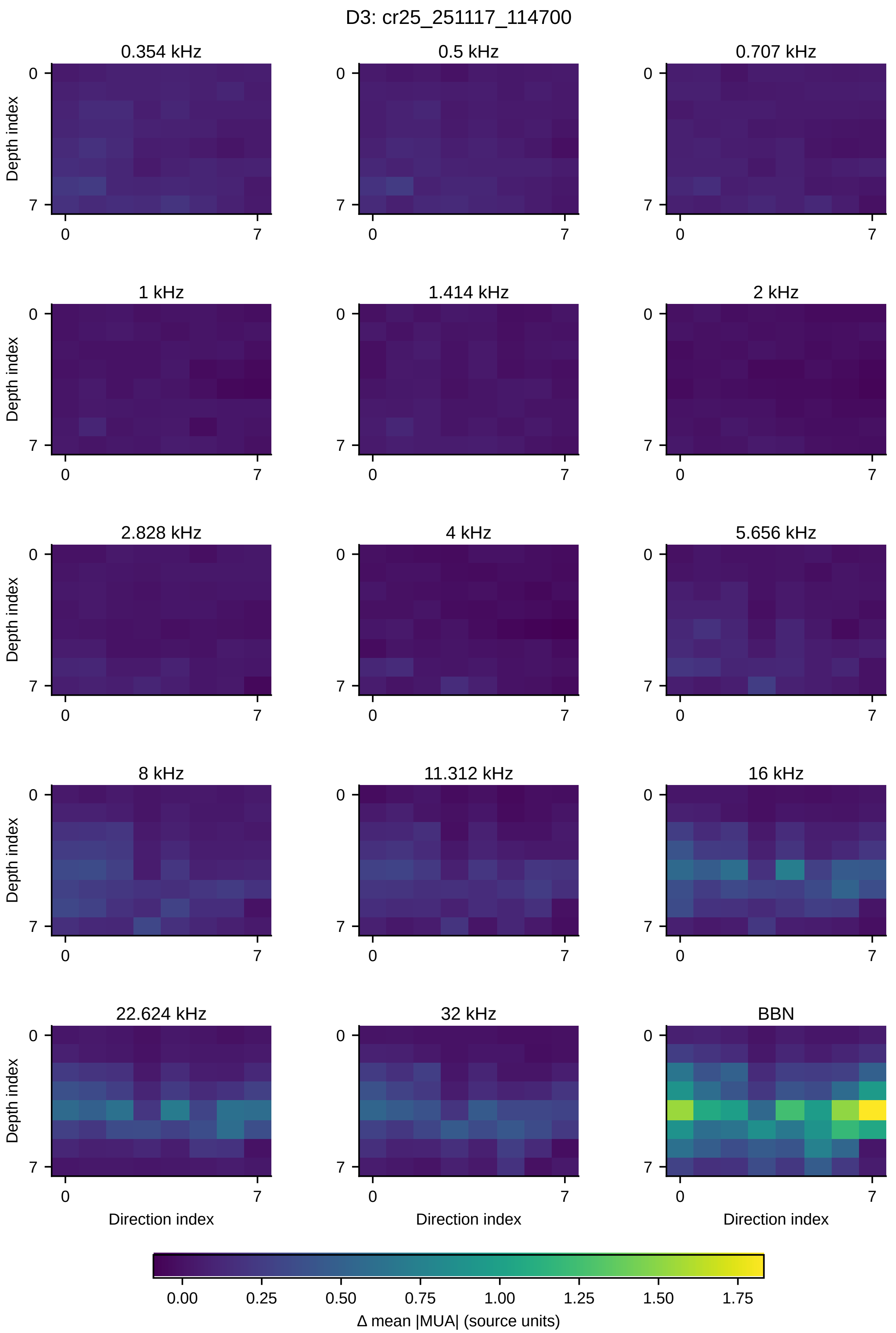
D3: cr25_251117_114700
0.354 kHz
0.5 kHz
0.707 kHz
1 kHz
1.414 kHz
2 kHz
2.828 kHz
4 kHz
5.656 kHz
8 kHz
11.312 kHz
16 kHz
22.624 kHz
32 kHz
BBN
Depth index
Direction index
0
7
0.00
0.25
0.50
0.75
1.00
1.25
1.50
1.75
Δ mean |MUA| (source units)

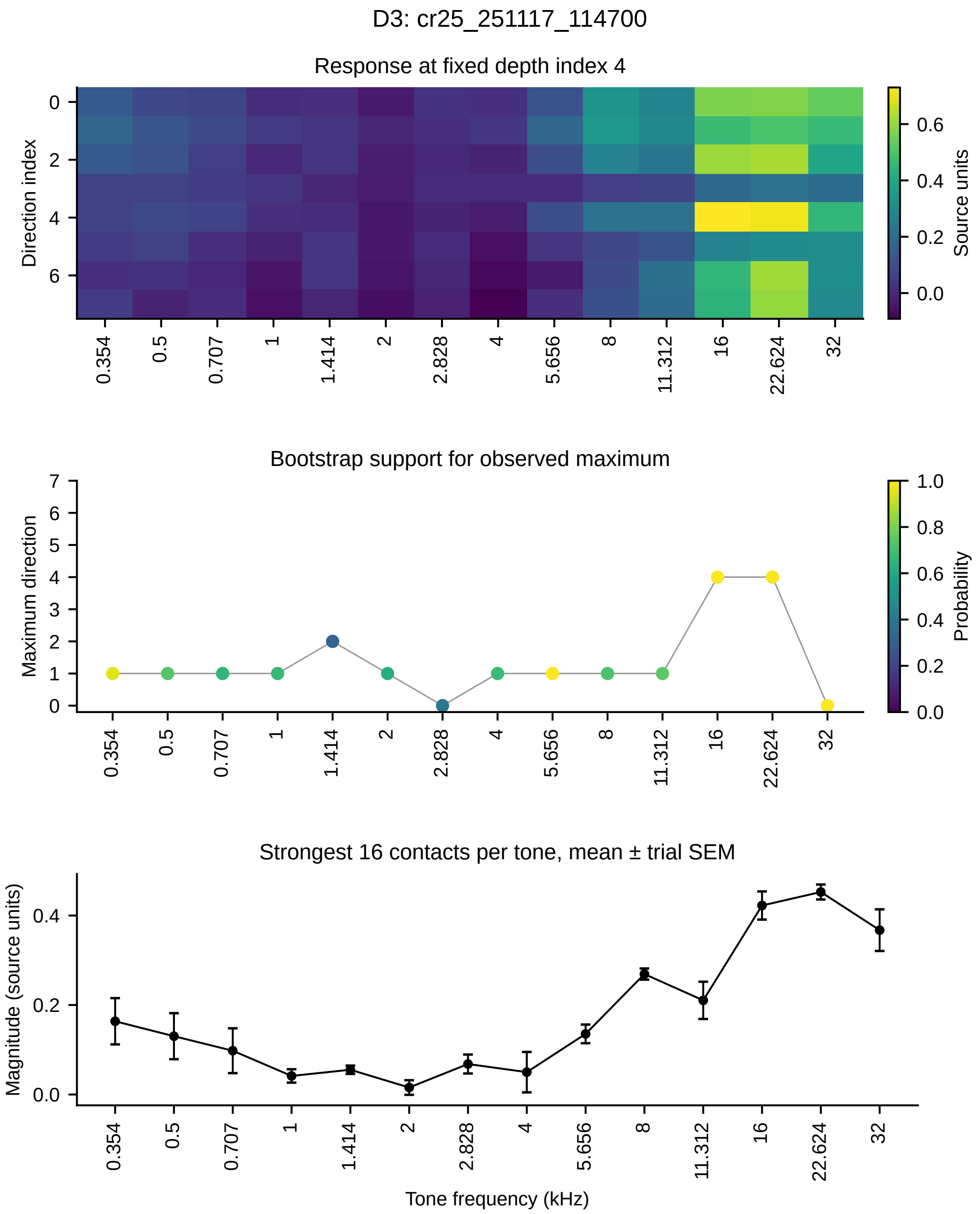
D3: cr25_251117_114700
Response at fixed depth index 4
Direction index
0
2
4
6
Source units
0.6
0.4
0.2
0.0
Bootstrap support for observed maximum
Maximum direction
7
6
5
4
3
2
1
0
Probability
1.0
0.8
0.6
0.4
0.2
0.0
Strongest 16 contacts per tone, mean ± trial SEM
Magnitude (source units)
0.4
0.2
0.0
0.354
0.5
0.707
1
1.414
2
2.828
4
5.656
8
11.312
16
22.624
32
Tone frequency (kHz)

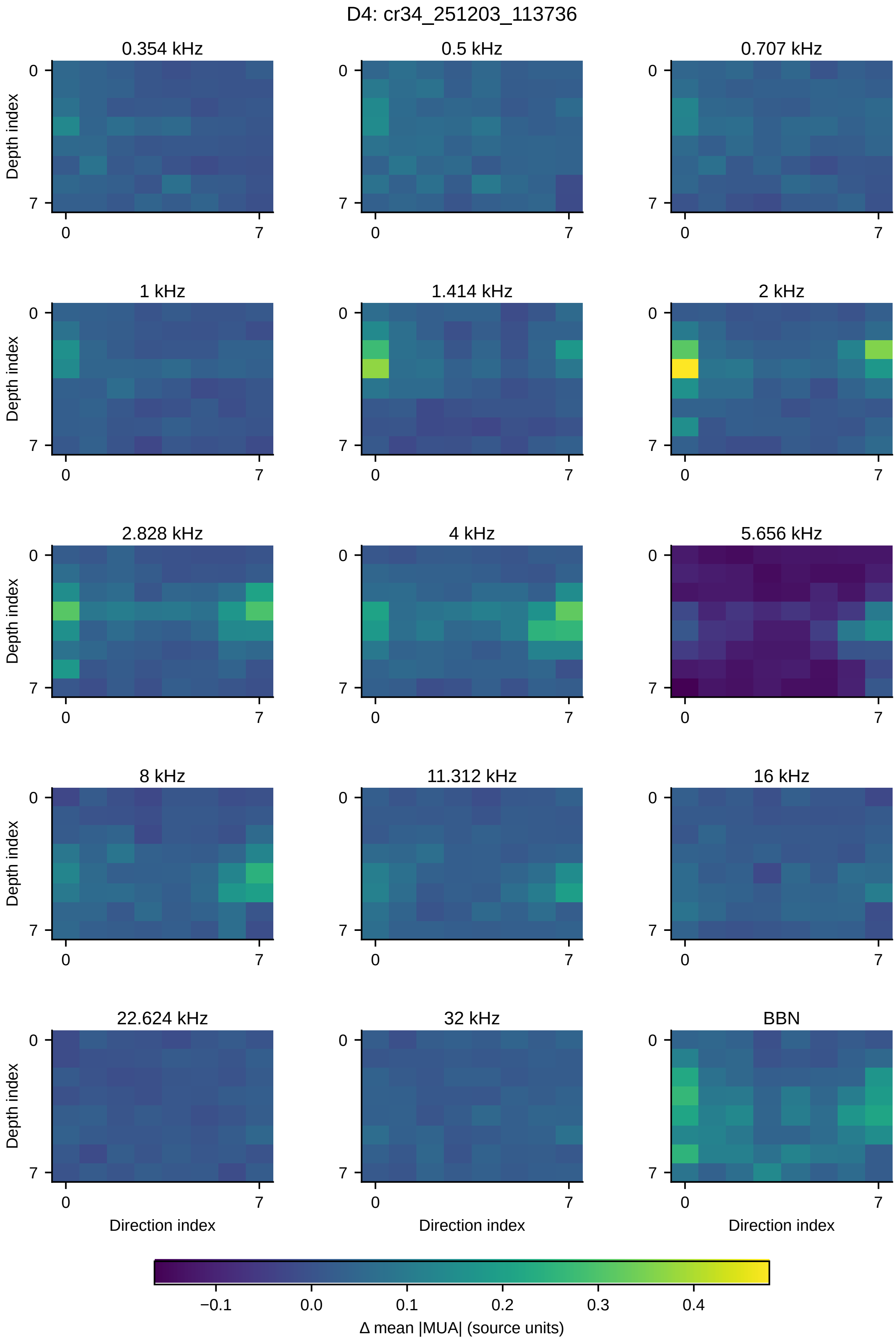
D4: cr34_251203_113736
0.354 kHz
0.5 kHz
0.707 kHz
1 kHz
1.414 kHz
2 kHz
2.828 kHz
4 kHz
5.656 kHz
8 kHz
11.312 kHz
16 kHz
22.624 kHz
32 kHz
BBN
Depth index
Direction index
0
7
−0.1
0.0
0.1
0.2
0.3
0.4
Δ mean |MUA| (source units)

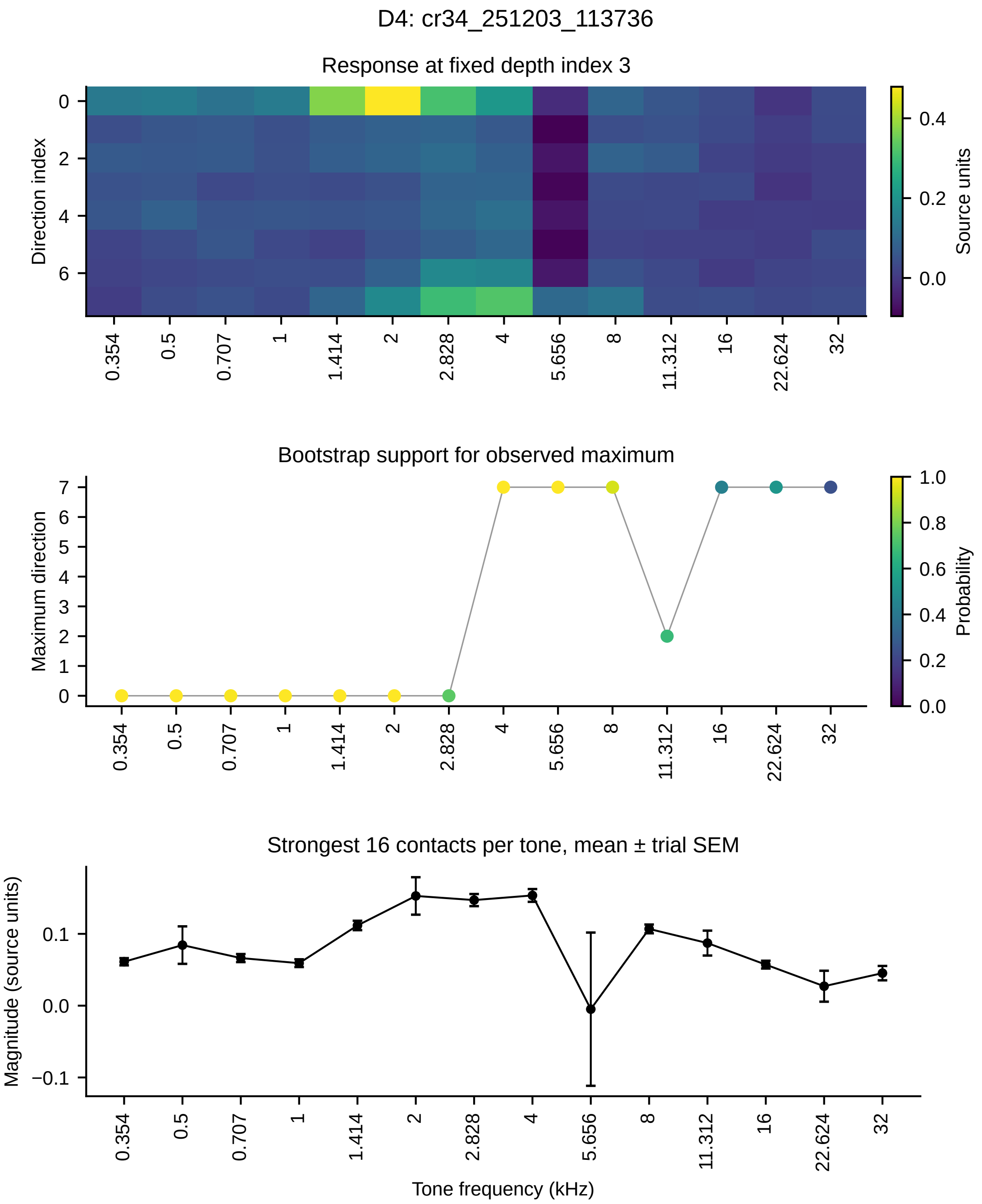

D4: cr34_251203_113736
Response at fixed depth index 3
Direction index
Source units
Bootstrap support for observed maximum
Maximum direction
Probability
Strongest 16 contacts per tone, mean ± trial SEM
Magnitude (source units)
Tone frequency (kHz)
0.354
0.5
0.707
1
1.414
2
2.828
4
5.656
8
11.312
16
22.624
32

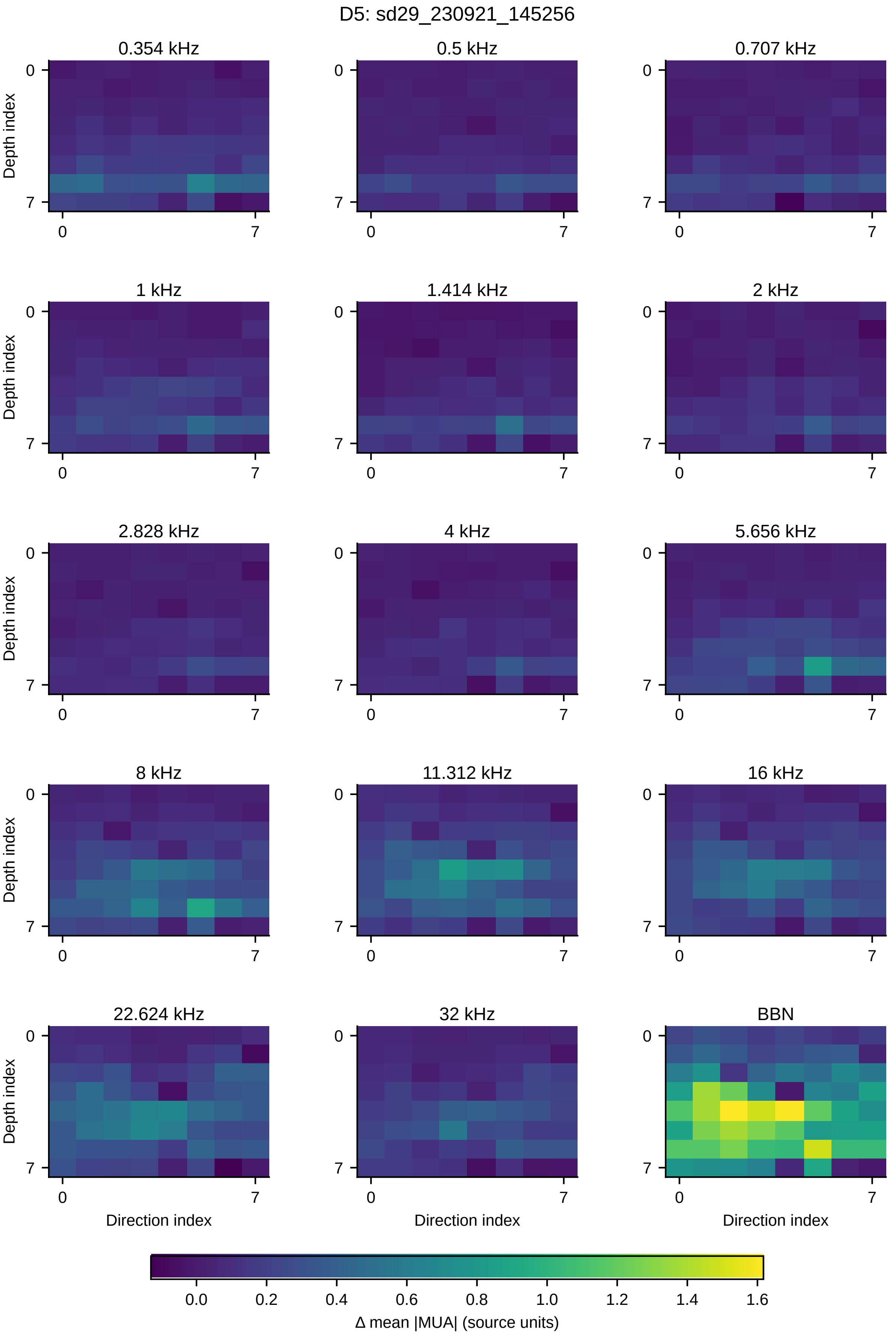
D5: sd29_230921_145256
0.354 kHz
0.5 kHz
0.707 kHz
1 kHz
1.414 kHz
2 kHz
2.828 kHz
4 kHz
5.656 kHz
8 kHz
11.312 kHz
16 kHz
22.624 kHz
32 kHz
BBN
Depth index
Direction index
0
7
0.0
0.2
0.4
0.6
0.8
1.0
1.2
1.4
1.6
Δ mean |MUA| (source units)

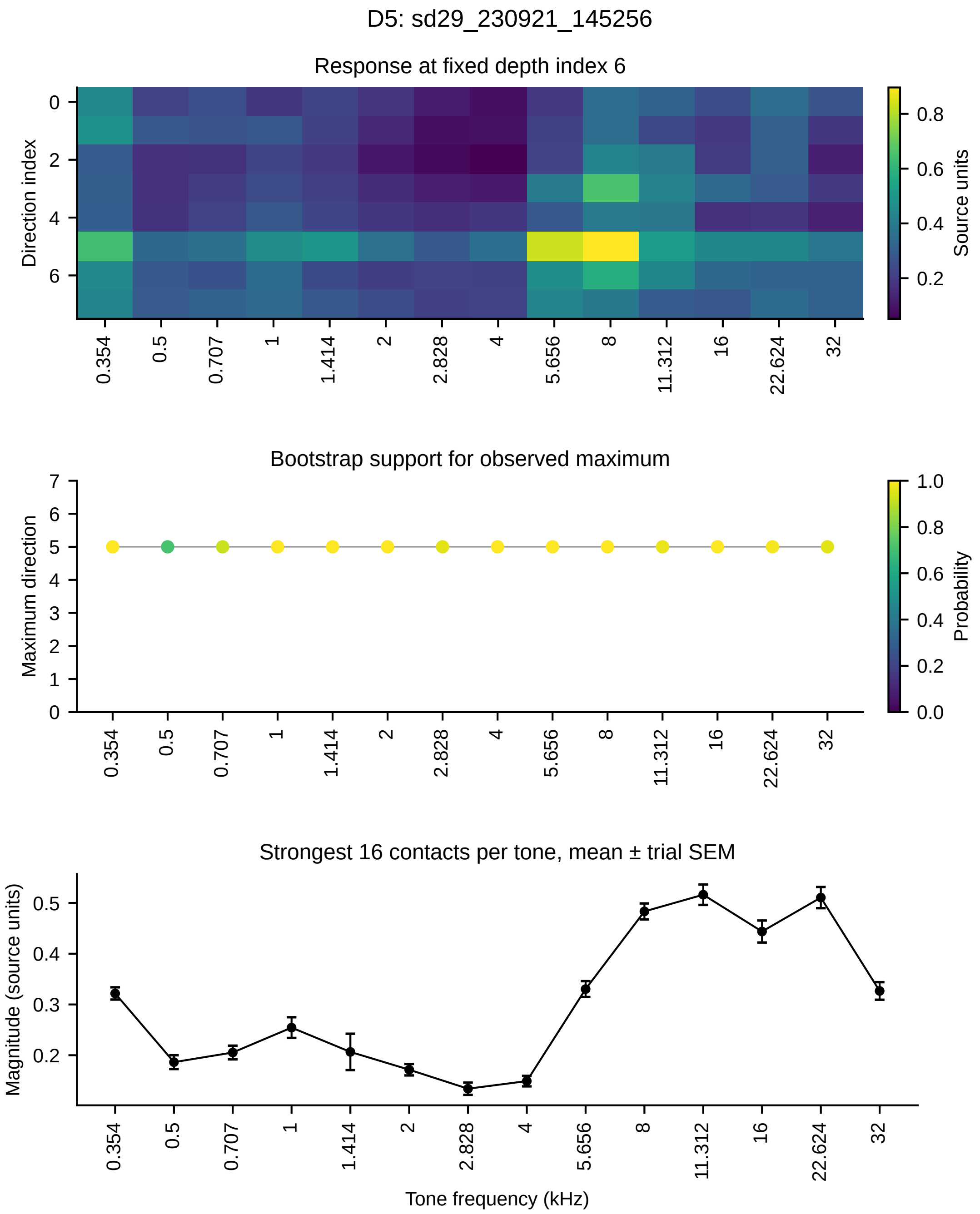
D5: sd29_230921_145256
Response at fixed depth index 6
Direction index
Source units
Bootstrap support for observed maximum
Maximum direction
Probability
Strongest 16 contacts per tone, mean ± trial SEM
Magnitude (source units)
Tone frequency (kHz)
0.354
0.5
0.707
1
1.414
2
2.828
4
5.656
8
11.312
16
22.624
32

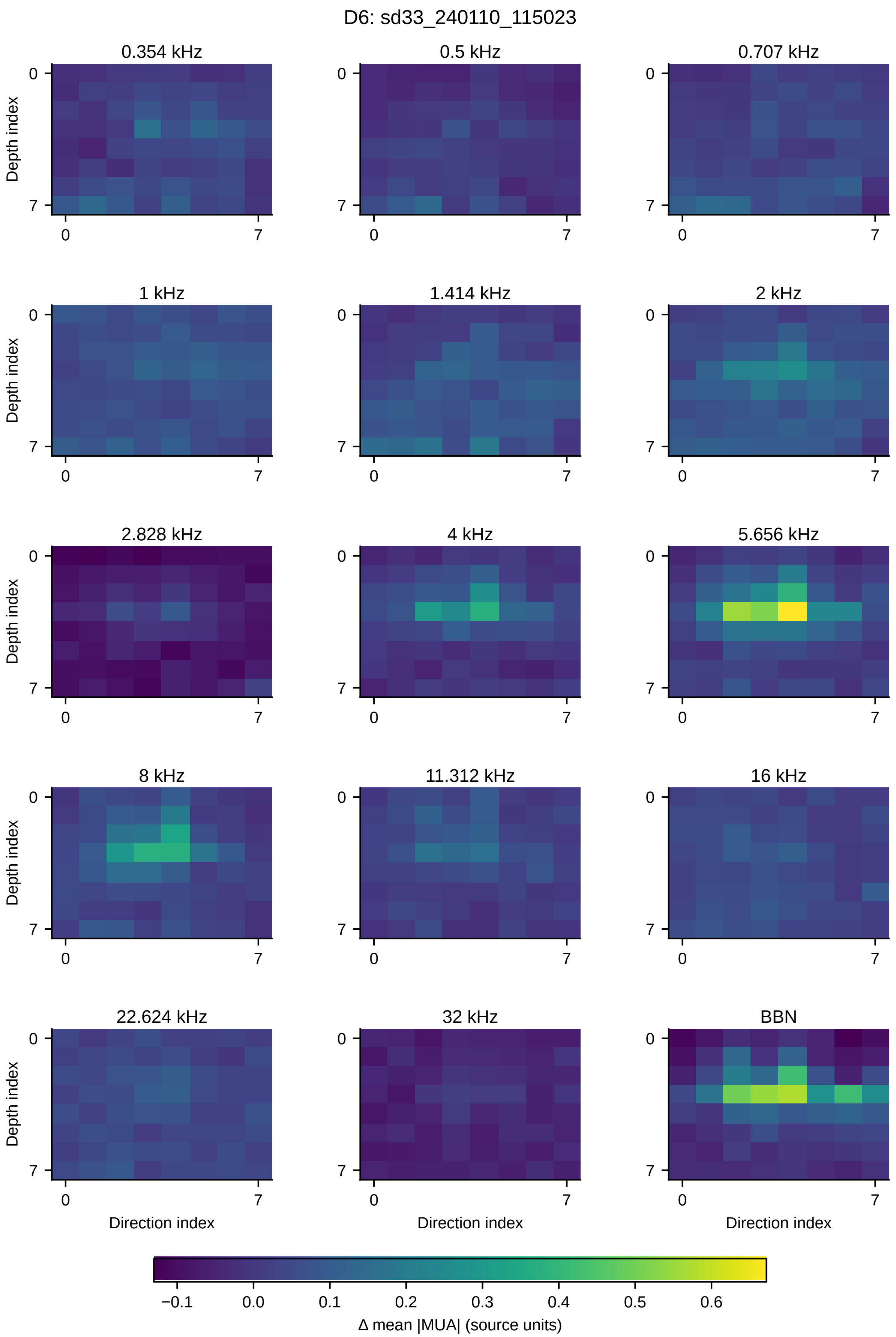
D6: sd33_240110_115023
0.354 kHz
0.5 kHz
0.707 kHz
1 kHz
1.414 kHz
2 kHz
2.828 kHz
4 kHz
5.656 kHz
8 kHz
11.312 kHz
16 kHz
22.624 kHz
32 kHz
BBN
Depth index
Direction index
0
7
−0.1
0.0
0.1
0.2
0.3
0.4
0.5
0.6
Δ mean |MUA| (source units)

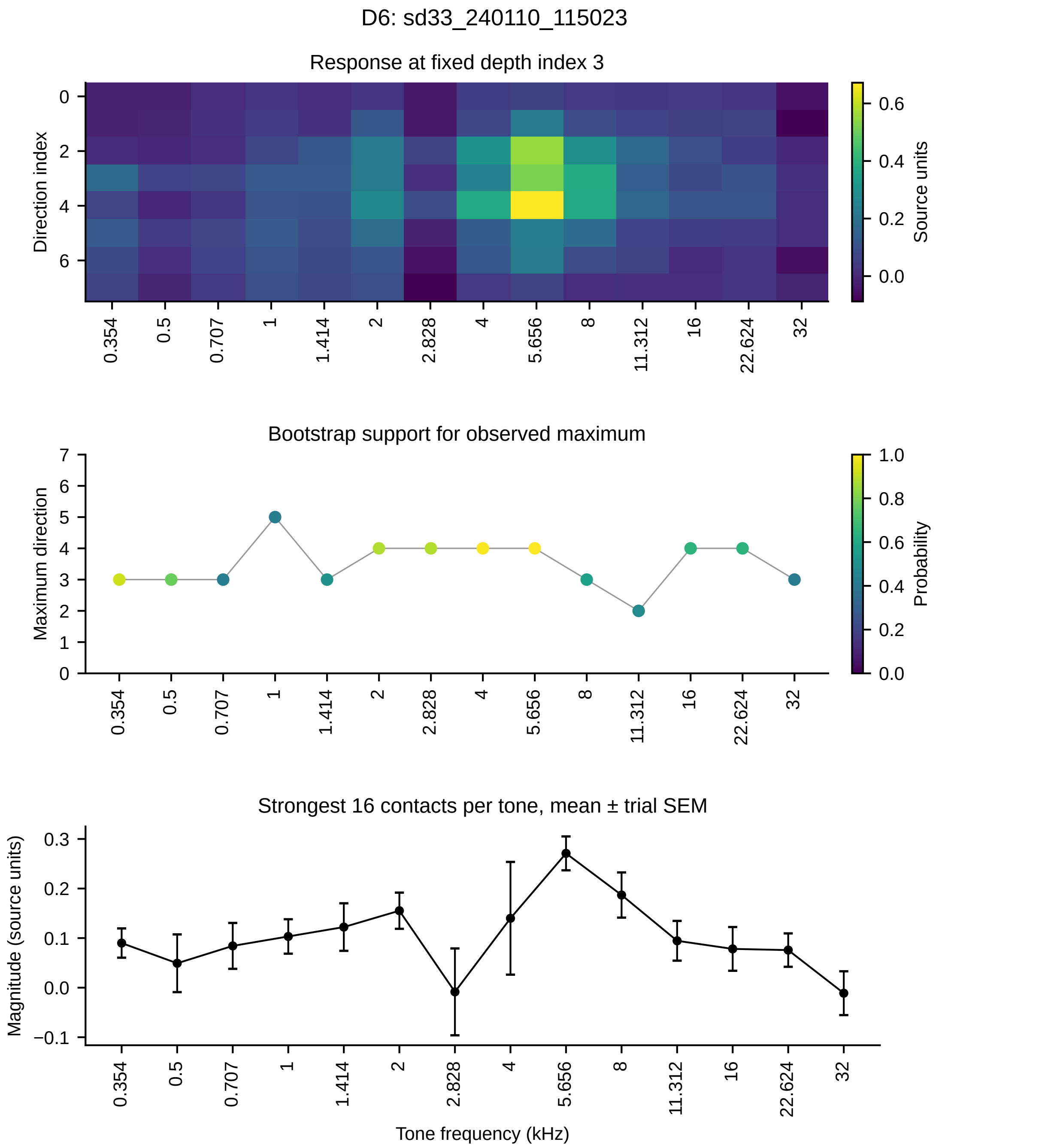
D6: sd33_240110_115023
Response at fixed depth index 3
Direction index
Source units
Bootstrap support for observed maximum
Maximum direction
Probability
Strongest 16 contacts per tone, mean ± trial SEM
Magnitude (source units)
Tone frequency (kHz)
0.354
0.5
0.707
1
1.414
2
2.828
4
5.656
8
11.312
16
22.624
32

#### S6. Complete UPROBE frequency-response profiles

All eight UPROBE penetrations are shown, with acoustic tone frequencies labeled in kHz. **(A)** Heat maps show baseline-subtracted mean absolute MUA responses across all 23 contacts for fourteen tones and broadband noise (BBN); dashed lines mark the contiguous ten-contact region selected from the signed mean-tone profile. **(B)** Correlations between BBN and tone spatial-response profiles across all 23 contacts; shading shows 95% trial-bootstrap intervals. **(C)** Tone-response magnitudes within the selected region, calculated after negative baseline-subtracted contact-mean responses are set to zero. Error bars show trial SEM conditional on the selected region and the contacts with positive mean responses. The same region is used for every tone within a penetration. These SEM values do not include uncertainty from choosing the region or setting negative values to zero. Amplitudes are in stored acquisition counts. Frequencies are plotted in ascending order, although stimuli were delivered in a fixed repeated sequence.

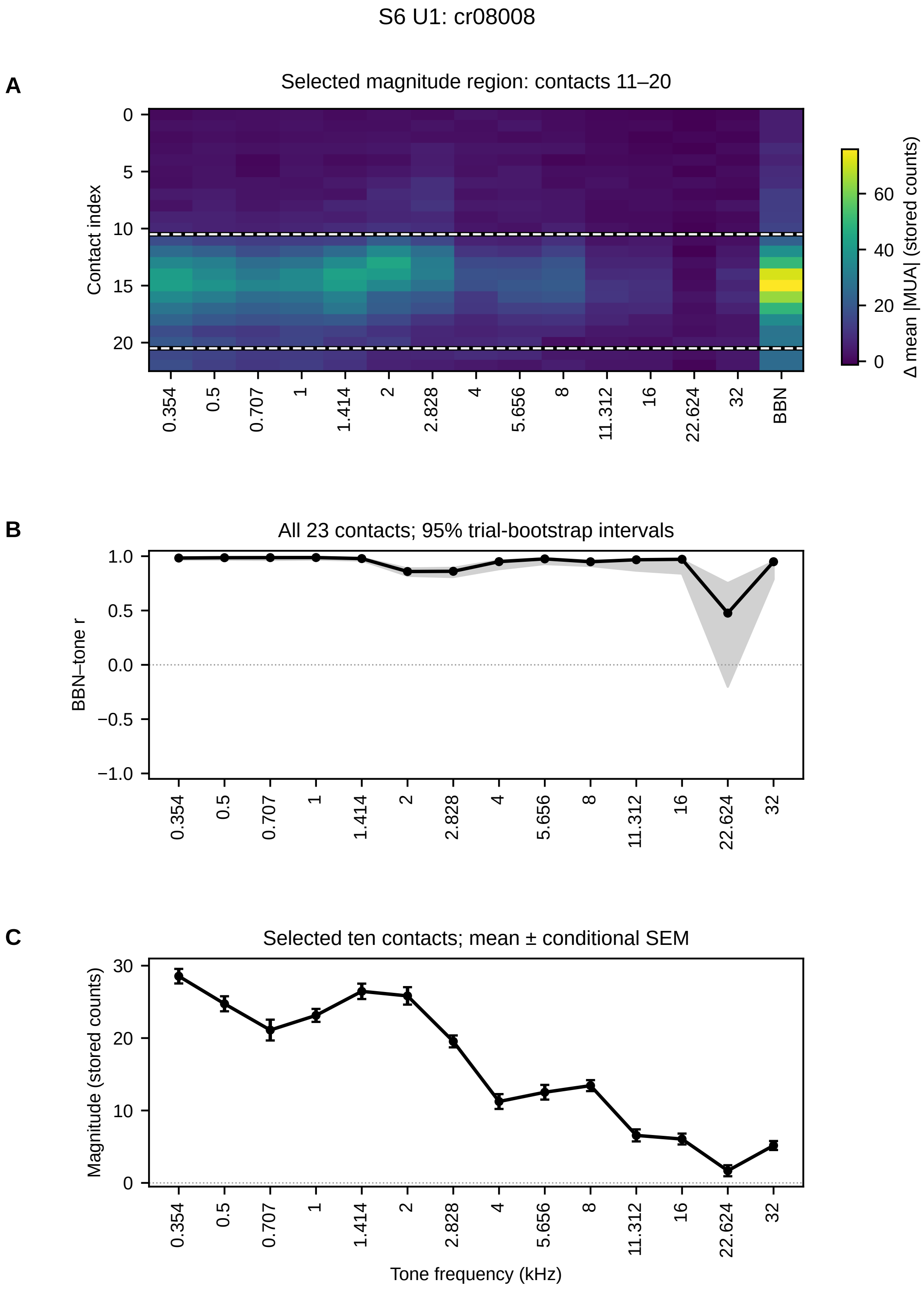
S6 U1: cr08008
A
Selected magnitude region: contacts 11–20
Contact index
Δ mean |MUA| (stored counts)
0.354
0.5
0.707
1
1.414
2
2.828
4
5.656
8
11.312
16
22.624
32
BBN
B
All 23 contacts; 95% trial-bootstrap intervals
BBN–tone r
C
Selected ten contacts; mean ± conditional SEM
Magnitude (stored counts)
Tone frequency (kHz)

S6 U2: cr11005

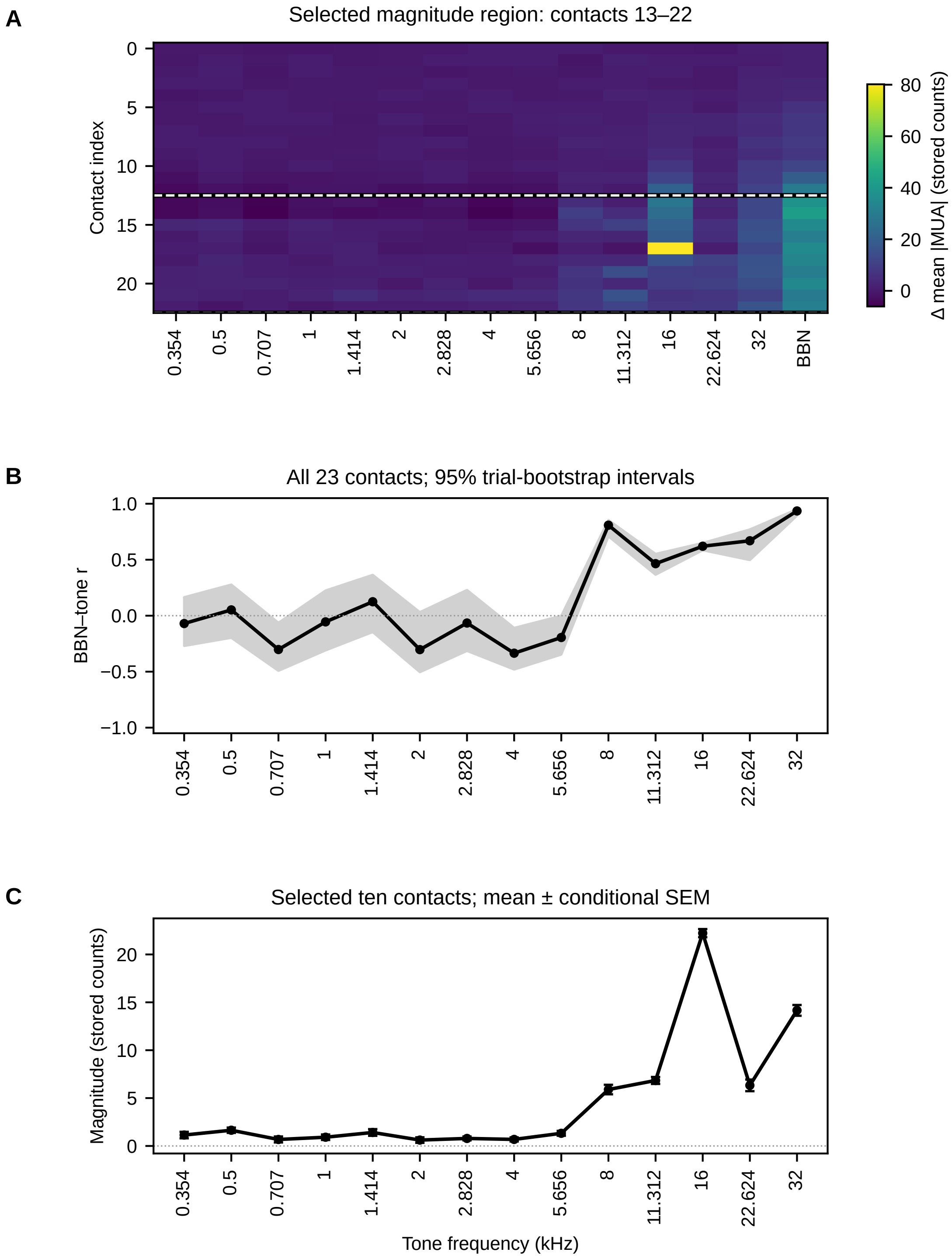

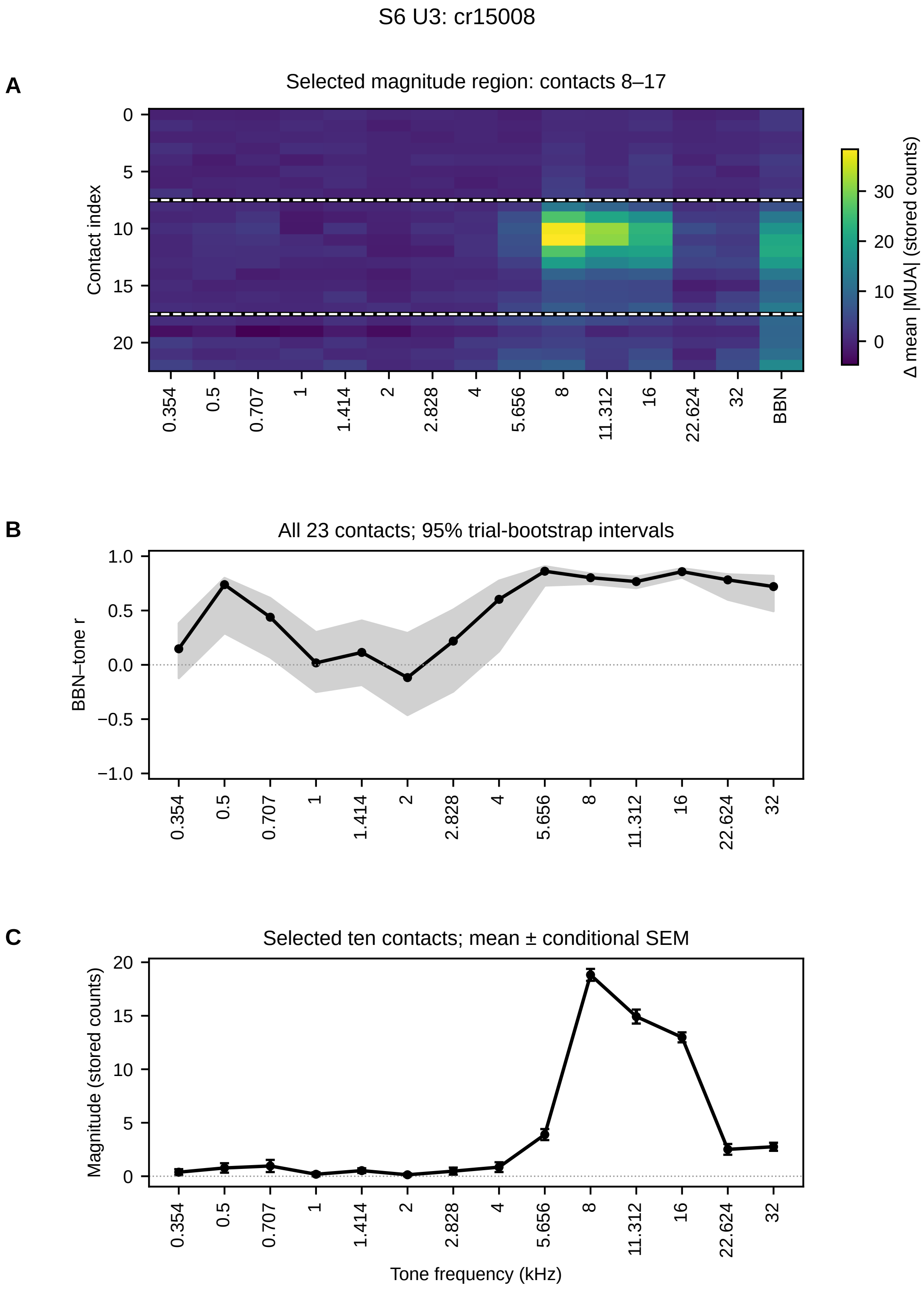
S6 U3: cr15008
A
Selected magnitude region: contacts 8–17
Contact index
Δ mean |MUA| (stored counts)
BBN
B
All 23 contacts; 95% trial-bootstrap intervals
BBN–tone r
C
Selected ten contacts; mean ± conditional SEM
Magnitude (stored counts)
Tone frequency (kHz)

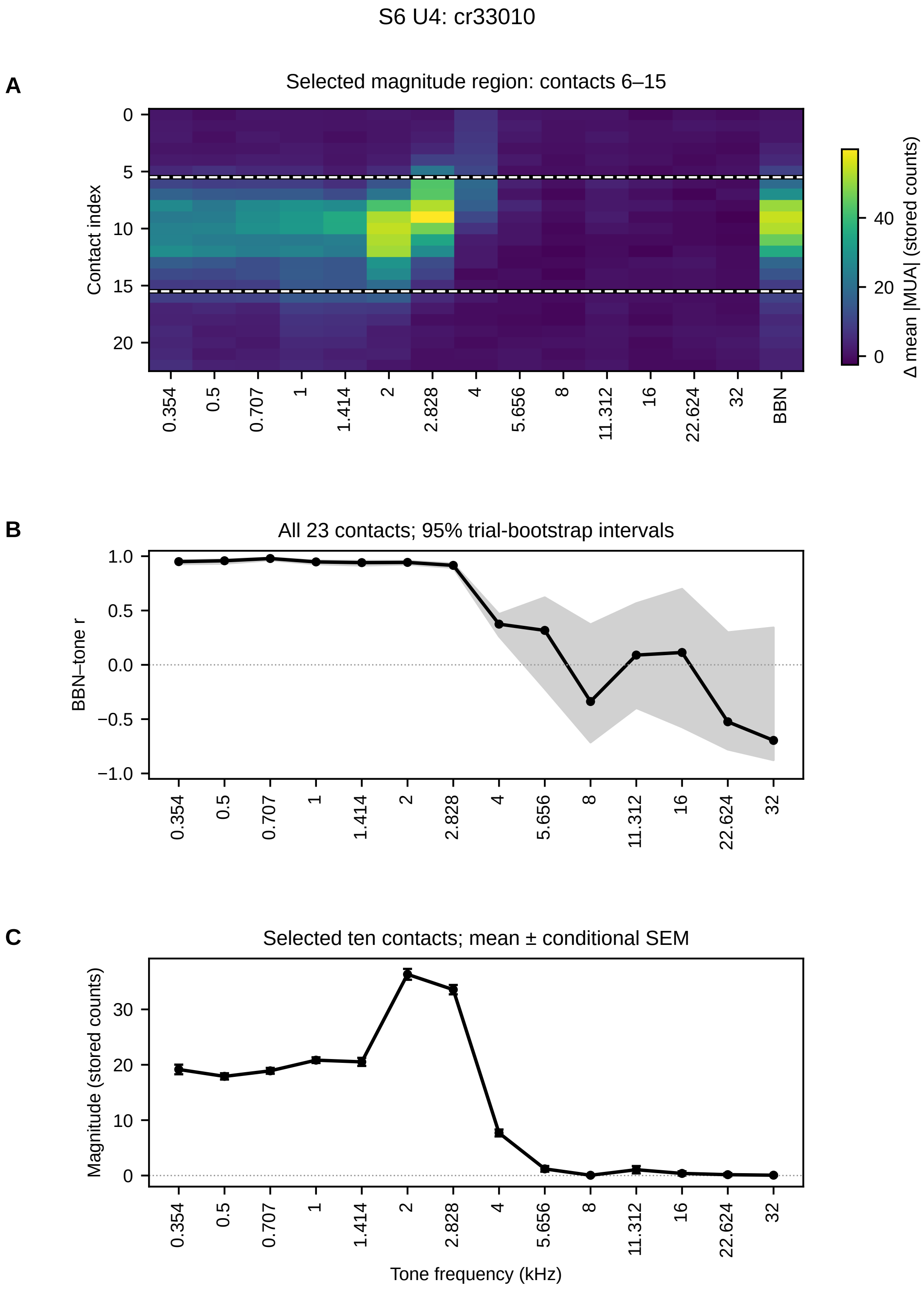
S6 U4: cr33010
A
Selected magnitude region: contacts 6–15
Contact index
0
5
10
15
20
0.354
0.5
0.707
1
1.414
2
2.828
4
5.656
8
11.312
16
22.624
32
BBN
Δ mean |MUA| (stored counts)
0
20
40
B
All 23 contacts; 95% trial-bootstrap intervals
BBN–tone r
1.0
0.5
0.0
−0.5
−1.0
C
Selected ten contacts; mean ± conditional SEM
Magnitude (stored counts)
30
20
10
0
Tone frequency (kHz)

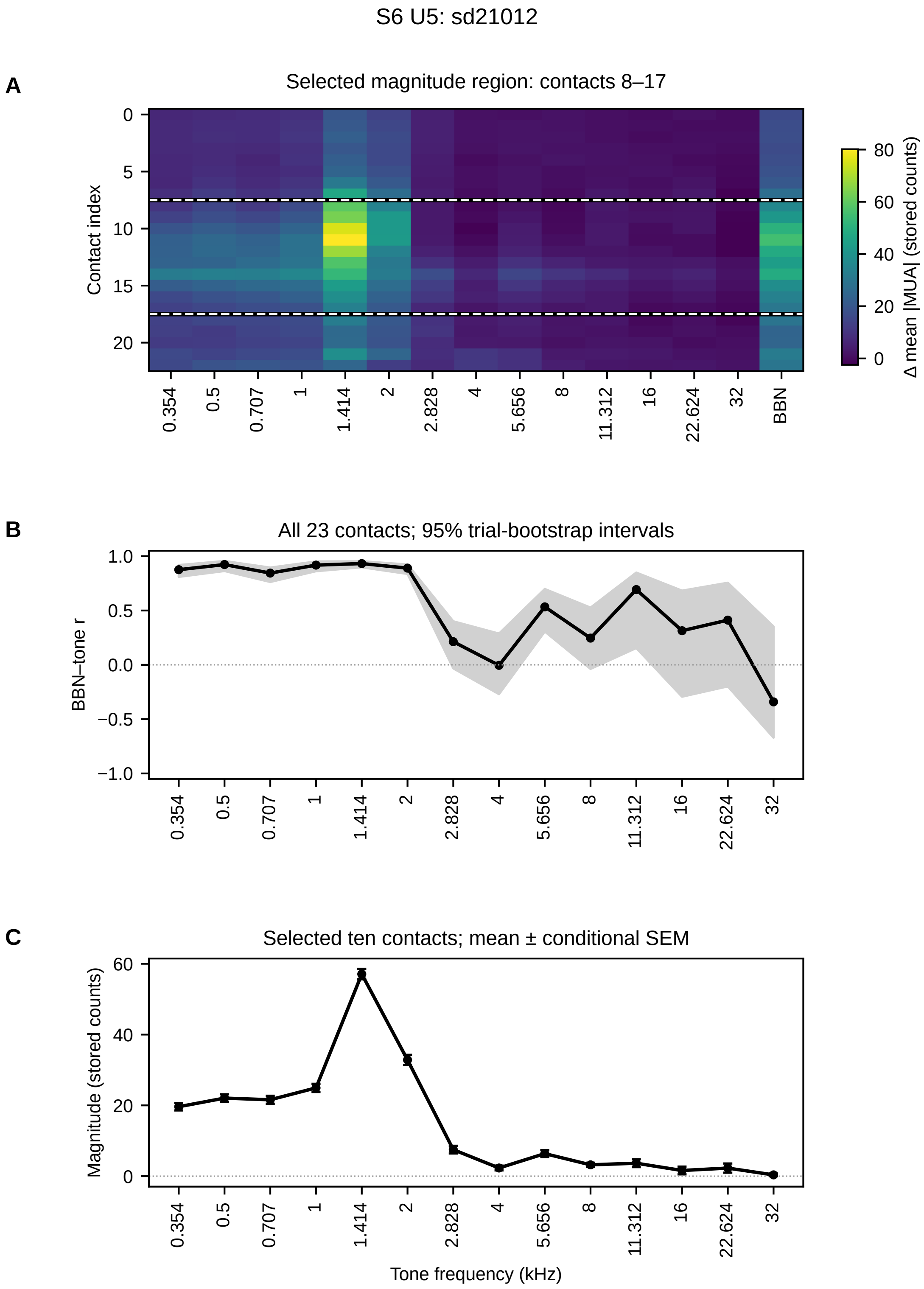
S6 U5: sd21012
A
Selected magnitude region: contacts 8–17
Contact index
Δ mean |MUA| (stored counts)
BBN
B
All 23 contacts; 95% trial-bootstrap intervals
BBN–tone r
C
Selected ten contacts; mean ± conditional SEM
Magnitude (stored counts)
Tone frequency (kHz)
0.354
0.5
0.707
1
1.414
2
2.828
4
5.656
8
11.312
16
22.624
32

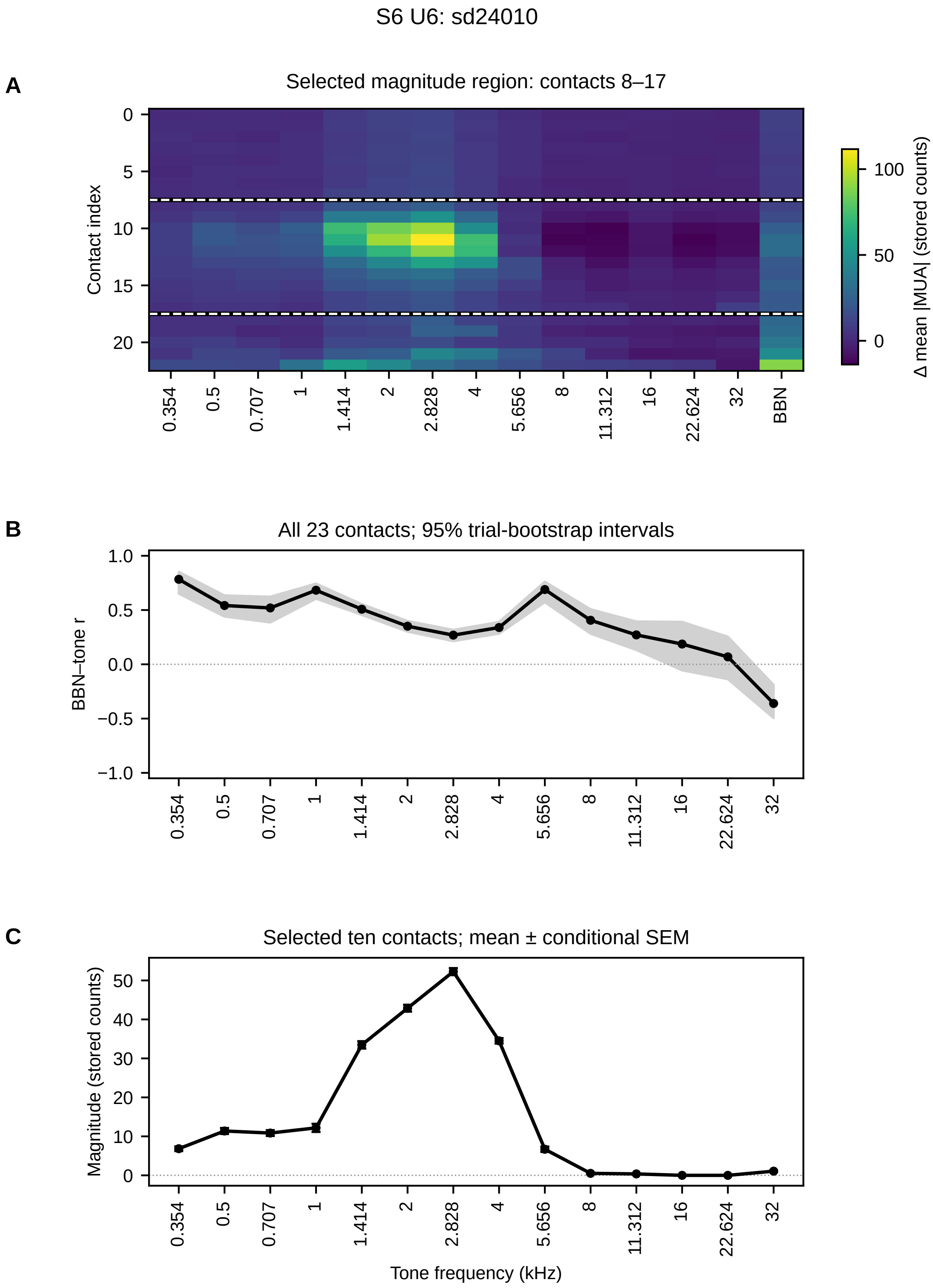
S6 U6: sd24010
A
Selected magnitude region: contacts 8–17
Contact index
Δ mean |MUA| (stored counts)
0
5
10
15
20
0.354
0.5
0.707
1
1.414
2
2.828
4
5.656
8
11.312
16
22.624
32
BBN
100
50
0
B
All 23 contacts; 95% trial-bootstrap intervals
BBN–tone r
1.0
0.5
0.0
−0.5
−1.0
C
Selected ten contacts; mean ± conditional SEM
Magnitude (stored counts)
50
40
30
20
10
0
Tone frequency (kHz)

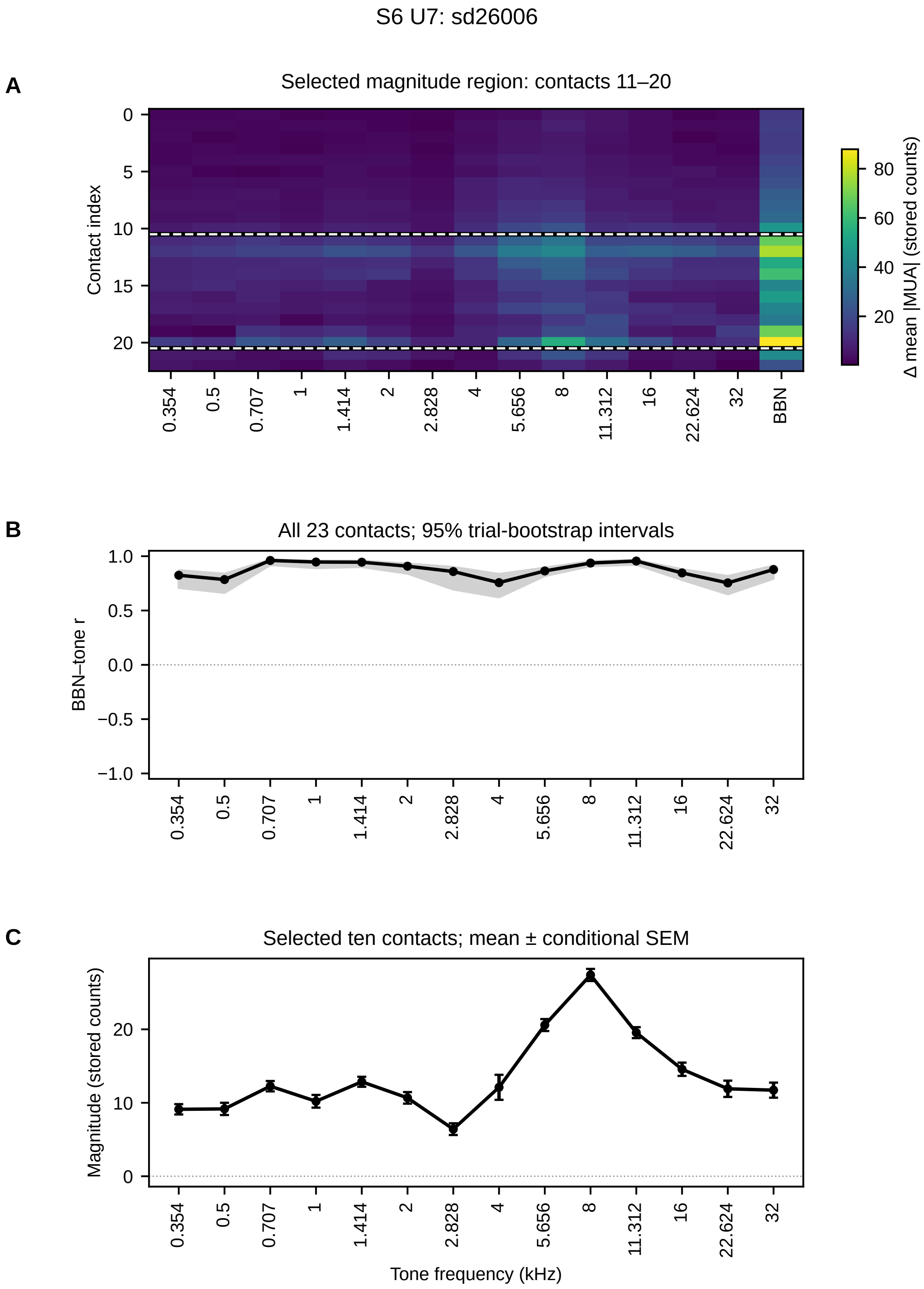
S6 U7: sd26006
A
Selected magnitude region: contacts 11–20
Contact index
0
5
10
15
20
0.354
0.5
0.707
1
1.414
2
2.828
4
5.656
8
11.312
16
22.624
32
BBN
80
60
40
20
Δ mean |MUA| (stored counts)
B
All 23 contacts; 95% trial-bootstrap intervals
BBN–tone r
1.0
0.5
0.0
−0.5
−1.0
C
Selected ten contacts; mean ± conditional SEM
Magnitude (stored counts)
20
10
0
Tone frequency (kHz)

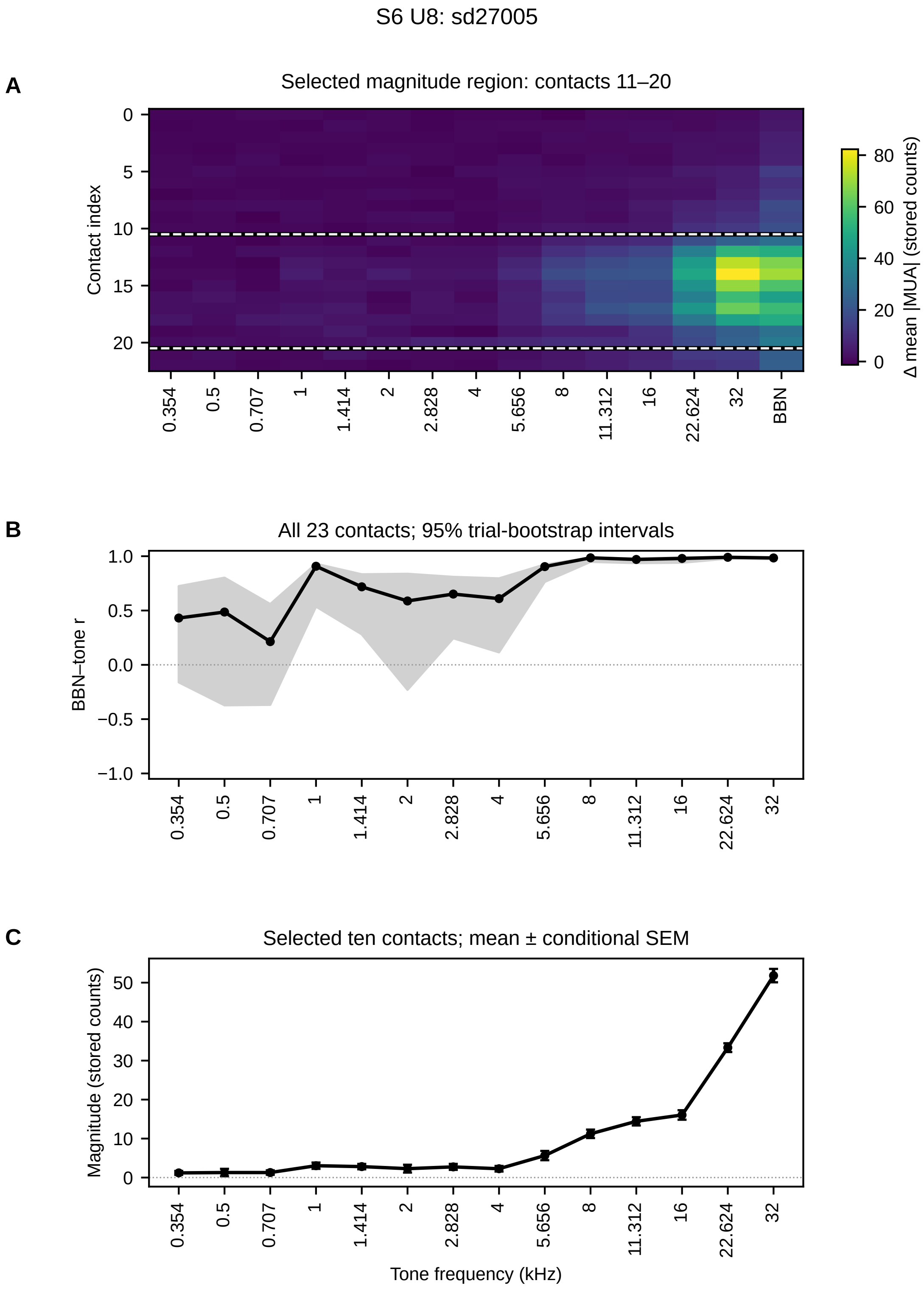
S6 U8: sd27005
A
Selected magnitude region: contacts 11–20
Contact index
0
5
10
15
20
0.354
0.5
0.707
1
1.414
2
2.828
4
5.656
8
11.312
16
22.624
32
BBN
Δ mean |MUA| (stored counts)
80
60
40
20
0
B
All 23 contacts; 95% trial-bootstrap intervals
BBN–tone r
1.0
0.5
0.0
−0.5
−1.0
C
Selected ten contacts; mean ± conditional SEM
Magnitude (stored counts)
50
40
30
20
10
0
Tone frequency (kHz)

### S7. UPROBE frequency-order comparison

Gray dashed curves show correlations in the former tone-label order, while blue curves show the same responses arranged by actual acoustic frequency. The response values are unchanged; their order along the frequency axis differs. Although this ordering reduces the apparent alternation between neighboring tones, frequency remains associated with stimulus position and preceding sound in the fixed repeated playback sequence.

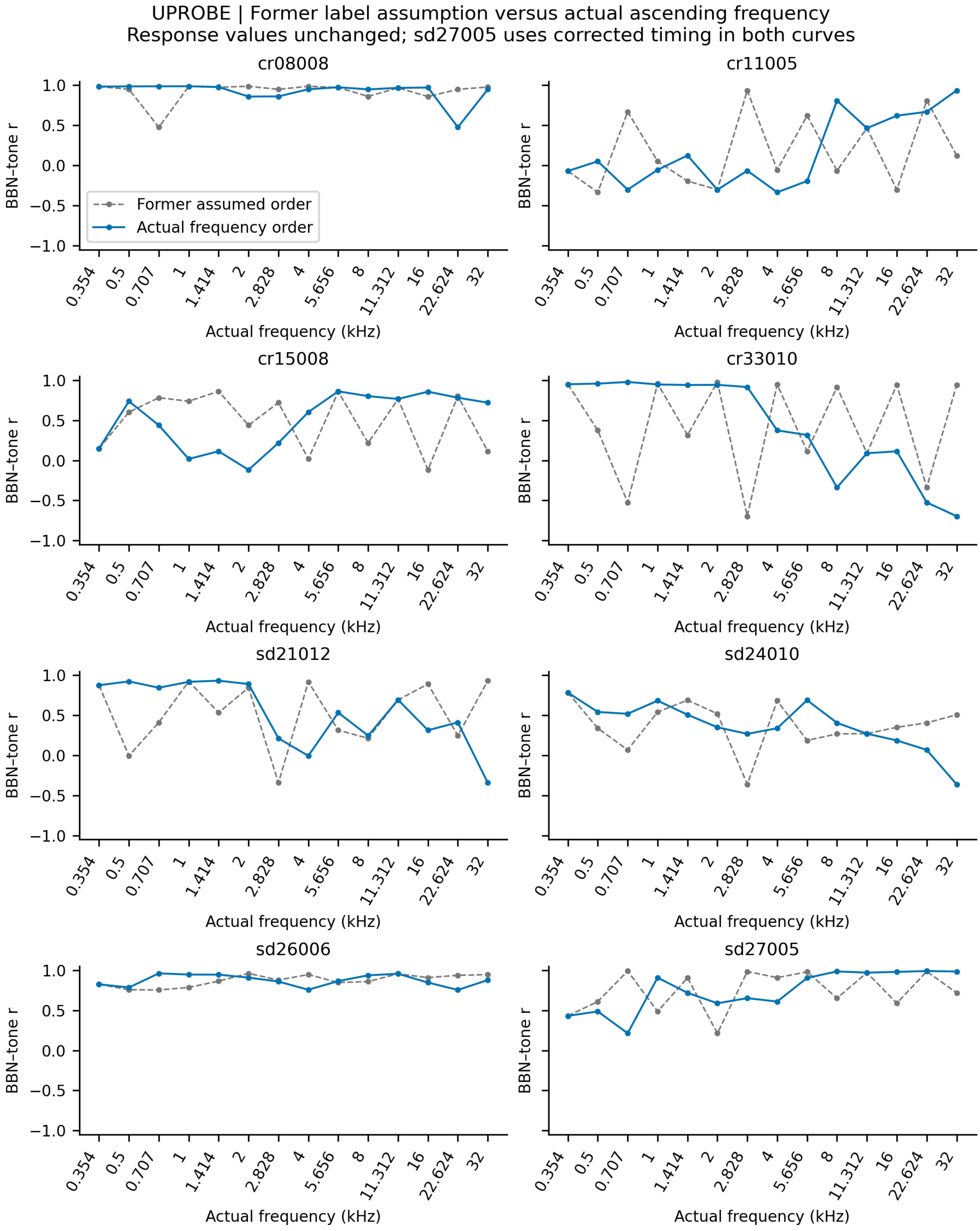
UPROBE | Former label assumption versus actual ascending frequency
Response values unchanged; sd27005 uses corrected timing in both curves
cr08008
cr11005
cr15008
cr33010
sd21012
sd24010
sd26006
sd27005
Former assumed order
Actual frequency order
BBN–tone r
Actual frequency (kHz)
1.0
0.5
0.0
−0.5
−1.0
0.354
0.5
0.707
1
1.414
2
2.828
4
5.656
8
11.312
16
22.624
32

### S8. Absolute population-response magnitudes

LFP and CSD RMS magnitudes and MUA-like mean envelope magnitudes over [0, 25) ms, after the signal-specific baseline procedures described in Methods. Symbols show mean ± SEM across 25 realizations per condition. Units follow the simulation forward calculation (mV for LFP and envelopes, mV/µm² for the unscaled coordinate-gradient CSD). These absolute magnitudes differ from percentages normalized across ten cortical classes within each realization. Individual and summary values are supplied as CSV tables.

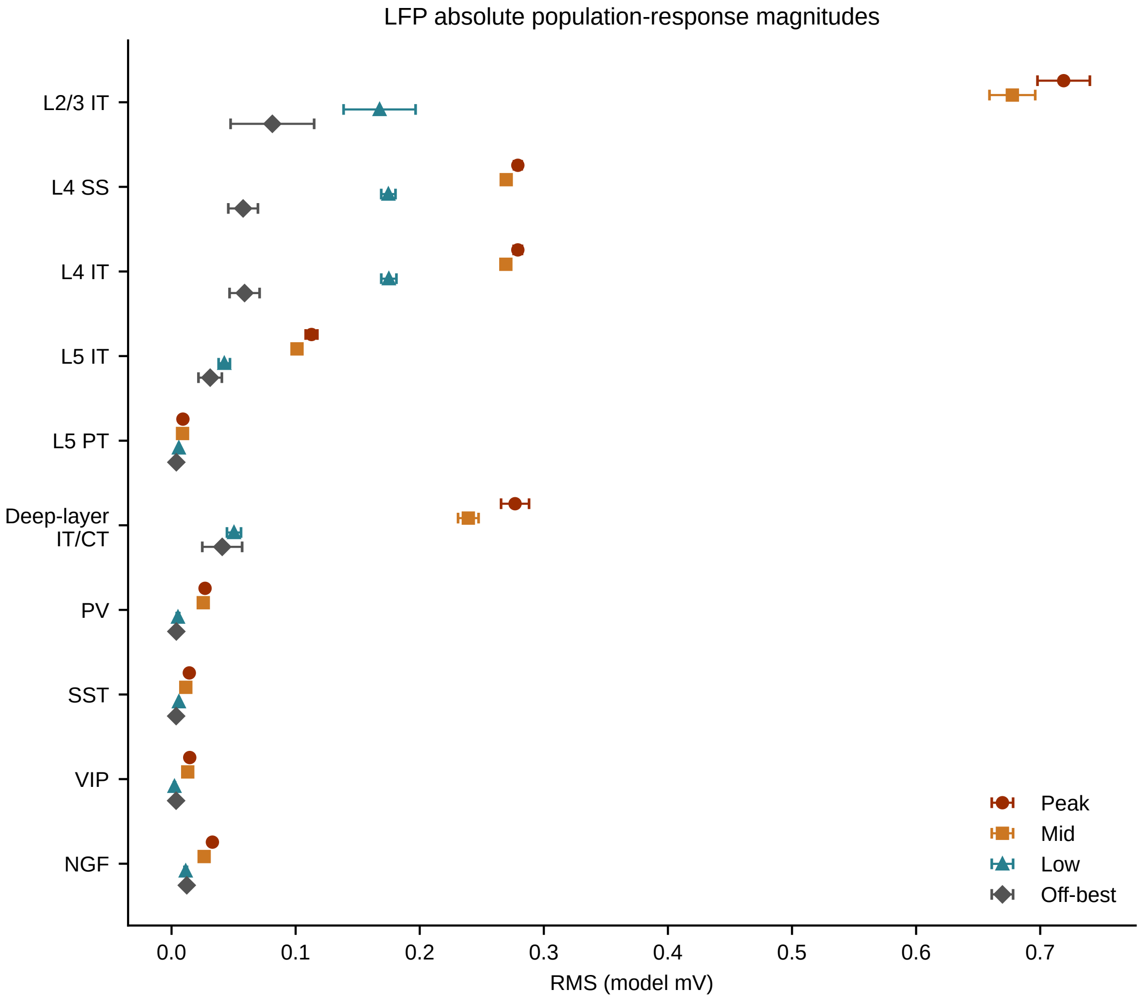
LFP absolute population-response magnitudes
L2/3 IT
L4 SS
L4 IT
L5 IT
L5 PT
Deep-layer IT/CT
PV
SST
VIP
NGF
0.0
0.1
0.2
0.3
0.4
0.5
0.6
0.7
RMS (model mV)
Peak
Mid
Low
Off-best

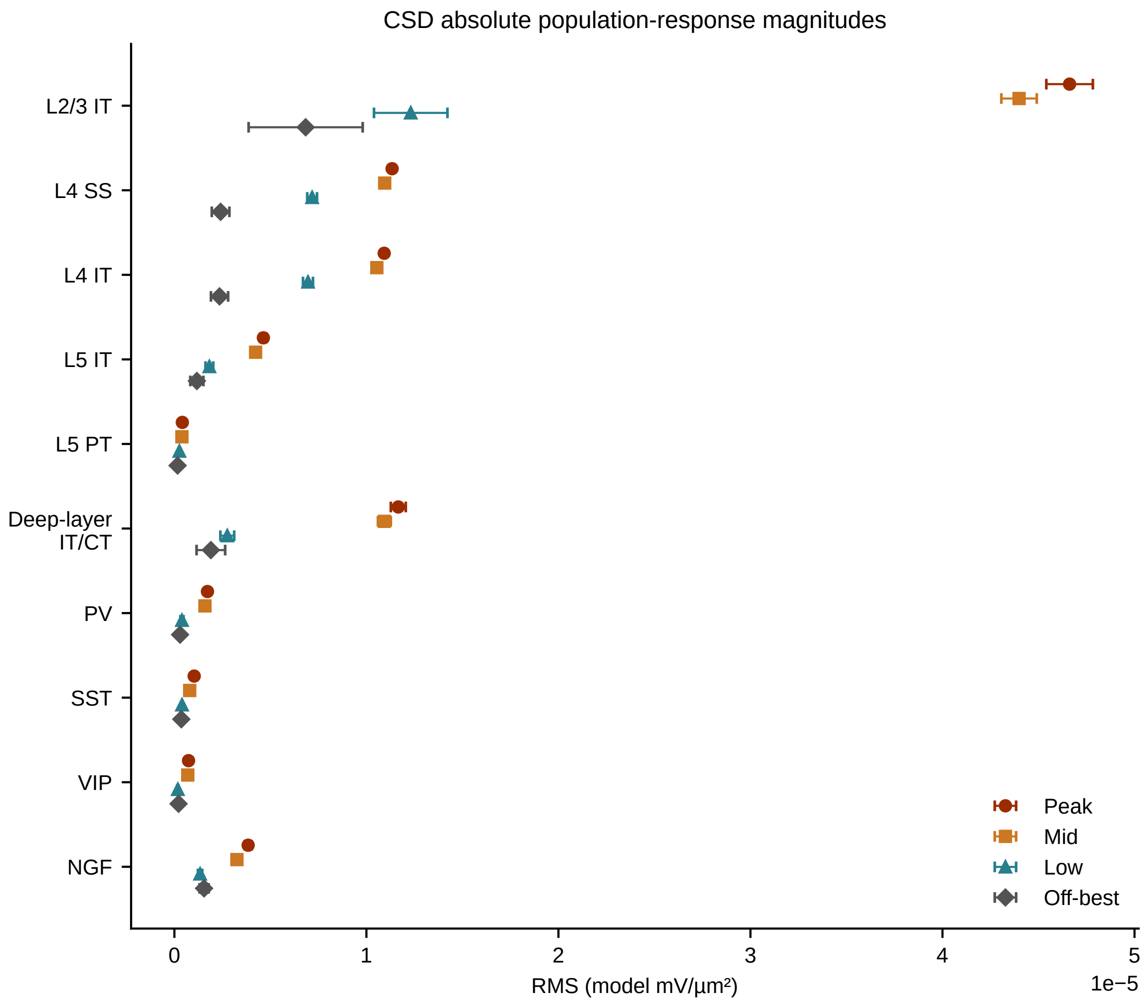
CSD absolute population-response magnitudes
L2/3 IT
L4 SS
L4 IT
L5 IT
L5 PT
Deep-layer IT/CT
PV
SST
VIP
NGF
0
1
2
3
4
5
1e−5
RMS (model mV/µm²)
Peak
Mid
Low
Off-best

**S8** MUA-like envelope magnitudes

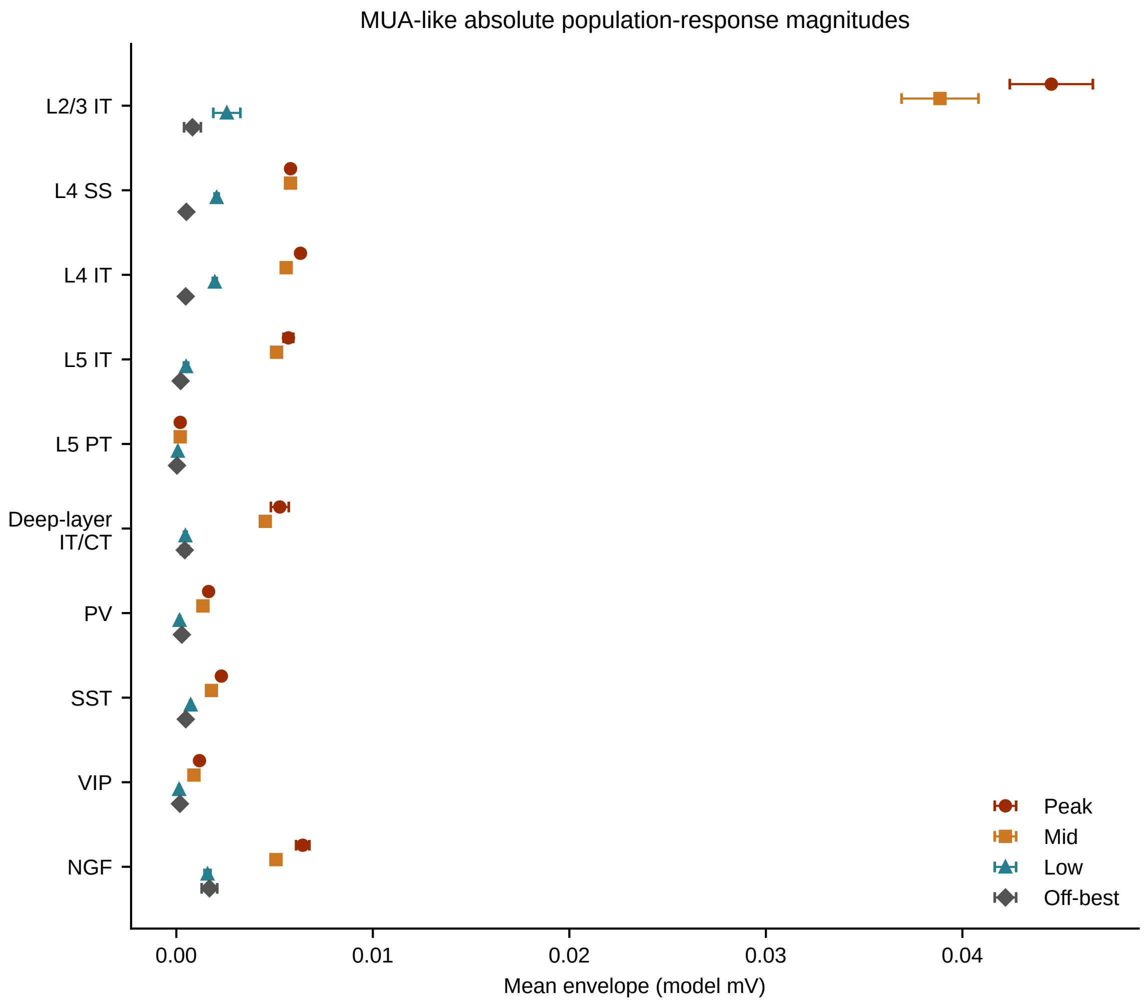

### S9. UPROBE contact orientation variation and region separability

The UPROBE replicates shown in Figure 4 were built with the contact face of the 23-contact laminar shank pointing along +y of the layout, perpendicular to the x axis along which the three Region layouts differ (Region 1 centred at x = −1.0 mm, Region 2 at 0, Region 3 at +1.0 mm). Because Regions 1 and 3 are mirrored about the device, the default placement aligns with the axis of symmetry. To test whether the orientation of the face is what limits the Uprobe, each replicate was rebuilt with the face rotated 90° about the shank so that it points along −x, towards the Region 1 centre. Everything else was held fixed. To keep the computation tractable, both orientations of all nine replicates (3 per Region) were built from a fixed pool of 20 of the 100 simulation trials, each column being assigned to the pool trial of its own condition. The two orientations of a replicate therefore record identical source columns with identical trials and are compared pairwise. The pool reproduces the main Uprobe replicates closely (per-contact LFP pattern cosine similarity 0.999, minimum 0.997, gain 0.90 of the main analysis, range 0.84–0.96, n = 9).

**(A)** Region-by-Region cosine distance between the per-contact LFP amplitude vectors of **(B)**, averaged over replicate pairs and expressed as a multiple of the within-Region distance, for the symmetric UPROBE, rotated UPROBE and DiSc (128 contacts). This compares directly to Figure 4 E&F. The outlined cell is the mirror pair R1–R3 and the star marks the largest off-diagonal distance. For UPROBE in either orientation the mirror pair remains the closest pair of Regions (R1–R3 distance 1.8 × within-Region for the symmetric case, 2.0 × for the rotated case), whereas for DiSc the mirror pair is the most distant (105 × within-Region). Region silhouette on the same distances: 0.48 (symmetric case), 0.47 (rotated case), and 0.98 (DiSc) against 0.51 for UPROBE in the main analysis.

**(B)** Per-contact LFP response amplitude (1–300 Hz, common average reference. RMS over 0–100 ms after stimulus onset) against depth for the symmetric orientation (solid orange) and the rotated orientation (dashed orange) as mean ± SD over three replicates per Region. Refer to Figure 4 for more detailed region depictions. Both devices shown to the right of figures are sized approximately to scale by depth. The DiSc columns that face the same directions are also included (Col 7 in solid blue; Col 1 in dashed blue). Dotted horizontal lines mark the cortical layer boundaries. Facing Region 1 raised the overall LFP gain there to 1.33 × the symmetric orientation (SD 0.05) and lowered it in Region 2 to 0.80 × (SD 0.03) and in Region 3 to 0.88 × (SD 0.05), so that the mirror-pair gain ratio gain(R3)/gain(R1) fell from 0.97 to 0.64. The same direction of change was seen for CSD, broadband high-frequency activity and MUA. The depth profile itself was little changed. The two DiSc columns show the same two behaviors, where the facing of Col 1 amplifies the region 1 and 3 difference. Rotating the face thus trades gain between the Regions, mainly by reducing the pattern difference between Region 2 and the lateral Regions, without creating a laminar pattern that distinguishes Region 1 from Region 3. Resolving the mirror pair requires sampling around the circumference, as with DiSc, where multiple column facings are recorded at once.

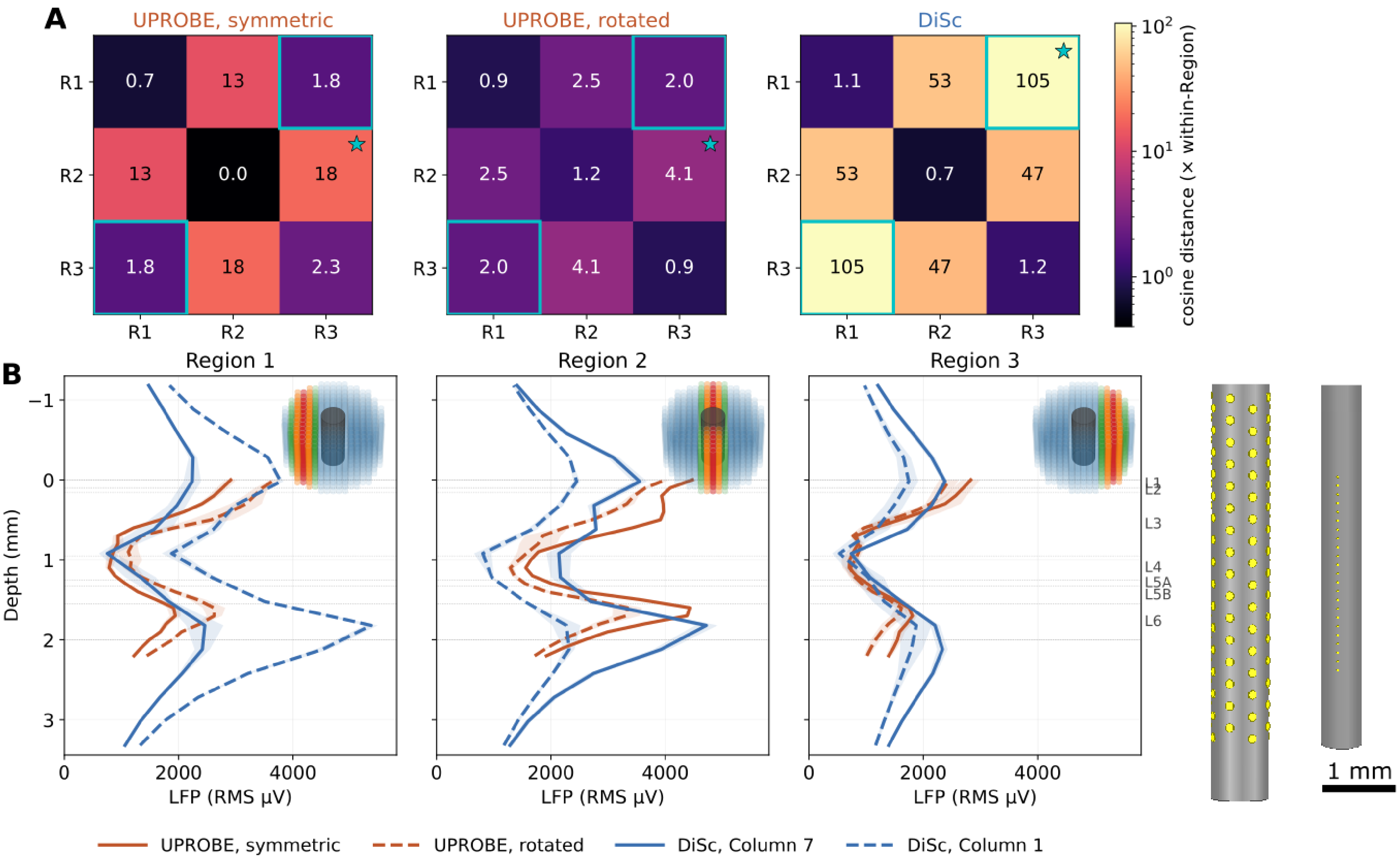
A
UPROBE, symmetric
UPROBE, rotated
DiSc
R1
R2
R3
0.7
13
1.8
0.0
18
2.3
0.9
2.5
2.0
1.2
4.1
1.1
53
105
0.7
47
1.2
cosine distance (× within-Region)
B
Region 1
Region 2
Region 3
Depth (mm)
LFP (RMS μV)
L1
L2
L3
L4
L5A
L5B
L6
1 mm
UPROBE, symmetric
UPROBE, rotated
DiSc, Column 7
DiSc, Column 1
